# Identifying JNK-regulated phosphoproteome markers of anxiety-like behaviour in mouse hippocampus

**DOI:** 10.1101/2025.02.06.636849

**Authors:** Ye Hong, Valentina Siino, Dani Flinkman, Prasannakumar Deshpande, Sylvia Ortega Martinez, Veronica Fagerholm, Artemis Varidaki, Pierre Heemeryck, Christel Sourander, Peter James, Eleanor Coffey

**Author notes:** Shared first author (Ye Hong, Valentina Siino).

## Abstract

JNKs mediate neuronal damage in neurodegenerative disease and inhibitors of JNK1 have shown anxiolytic and anti-depressive effects in mice. Here, we analyze the phosphoproteomes of hippocampus and nucleus accumbens from DJNKI-1 (JNK inhibitor)-infused mice. We correlate phospho-site changes with anxiety-like behaviours in the elevated plus maze and light-dark test and identify unique changes in responder mice. Among the DJNKI-1 regulated phosphosites, several lie within GSK3 motifs and are exclusively down-regulated. Consistent with this, GSK3β is inhibited and AKT activated. Importantly, we detect multilevel regulation of glucose metabolism enzymes including increased PDPK1-S241 phosphorylation, and a 5-fold increase in pyruvate dehydrogenase (PDHA1)-S-293 phosphorylation, signifying its inhibition. This suggests that JNK inhibitor drives a metabolic transformation to the neuronal Warburg response. This and a range of identified synaptic and cytoskeletal protein phosphosite changes are discussed in the context of JNK-regulated anxiety responses. The annotated hippocampal and nucleus accumbens phosphoproteomes described here will support a mechanistic understanding of the JNK pathway for future studies of brain disorders.

## Introduction

c-Jun amino terminal kinases (JNKs) are enzymatic sensors that respond to cellular stressors, including cytokines, endocrine signals, mechanical and ribotoxic stress. Upon activation, they phosphorylate target proteins that elicit diverse responses such as autophagy, microtubule stabilisation, cellular transport, insulin homeostasis, inflammation or neurodegeneration (Antoniou et al., 2011; Chang et al., 2003; Fu and Holzbaur, 2013; Hotamisligil and Davis, 2016; Hsu et al., 2010; Iordanov et al., 1997; Karpac et al., 2009; Kyriakis and Avruch, 2012; Morfini et al., 2009; Pereira et al., 2011; Vind et al., 2024; Xu et al., 2011; Zeng et al., 2022). Despite this important role, the phosphorylation targets of JNK and its precise mechanisms of action have not been systematically identified.

In the brain, JNKs are widely recognised for their role in mediating neuronal death in models of stroke, Alzheimer’s, Parkinson’s and Huntington’s diseases. Notably, JNK inhibitors have been shown to be neuroprotective in an expansive range of studies (Kumari et al., 2023; Repici and Borsello, 2006). In Alzheimer’s disease for example, beta-amyloid drives JNK activity leading to hyperphosphorylation of Tau which in turn contributes to neurofibrillary tangles, the pathological hallmark. Similarly, in Parkinson’s disease, JNK mediates key inflammatory signalling (Antoniou et al., 2011; Gehringer et al., 2015; Repici and Borsello, 2006; Solas et al., 2023).

More recently, JNK has been implicated in anxiety- and depressive like behaviours in animal studies. Thus inhibition or genetic deletion of *Jnk1* (*Mapk8*) induces a low-anxiety, low-depressive-like phenotype in rodents (Hollos et al., 2018; Meng et al., 2021; Mohammad et al., 2018; Zhao et al., 2017; Zhou et al., 2023). In a zebrafish phenotypic screen for anxiety lowering drugs, JNK inhibitors recapitulate the behavioural signature of classical anti-depressant drugs (Hong et al., 2024). The reduced anxiety/reduced depression/anhedonia phenotype upon JNK inhibitor treatment is associated with increased hippocampal neurogenesis, suggesting a potential mechanism (Castro-Torres et al., 2021; Mohammad et al., 2018).

In patients, JNK1 pathway genes have been shown to confer risk for psychosis (McCarthy et al., 2009; Winchester et al., 2012) and loss of function mutation in *JNK1 (MAPK8)* and truncations in *JNK3 (MAPK10)*, have been linked to intellectual disability (Harripaul et al., 2018; Kunde et al., 2013; Shoichet et al., 2006). Furthermore, upstream regulators on the JNK pathway confer risk for psychosis (McCarthy et al., 2009; Winchester et al., 2012). Beyond its role in neurogenesis, JNK contributes to immune dysfunction, which can also impact mental state (Arthur and Ley, 2013; Guma et al., 2011).

Despite the enormous literature describing the role of JNK in regulating stress responses, its *in vivo* impact on protein phosphorylation remains relatively unexplored at the proteome level. Understanding the downstream effects of JNK activity in brain is critical for determining how it contributes to these diverse functions. In this study, we address this gap using DJNKI-1, a well characterized peptide inhibitor of JNK (Borsello and Bonny, 2004; Borsello et al., 2003). We investigate how JNK inhibition affects behavior and protein phosphorylation in two key brain regions; the hippocampus, associated with learning, memory and mood regulation, and the nucleus accumbens, a central hub for reward and motivation. The results comprise an annotated view of the DJNKI-1 regulated phosphoproteome from hippocampus and nucleus accumbens. The data highlights JNK regulation of enzymes that control insulin signalling and glucose metabolism as well as extensive regulation of microtubule and actin regulatory and synaptic proteins.

## Material and methods

### Chronic intracerebroventricular (ICV) infusion

This procedure was carried out essentially as previously described (Mohammad et al., 2018). For analgesia, adult C57Bl/6J mice (2-month-old) were injected intraperitoneally with 0.1□mg□kg−1 of buprenorphine hydrochloride (Reckitt Benckiser Pharmaceuticals, Basel, Switzerland) and anaesthetized with 4 % isoflurane and maintained with 2.5 % isoflurane. Local anaesthetic (0.1□ml of 1□mg per 1□μg lidocaine) was subcutaneously injected, before cutting the scalp. A 100µl mini-osmotic pump (Alzet Model 2006, Cupertino, CA, USA) was inserted subcutaneously at the back of the neck and attached to a cannula that was implanted cerebroventricularly. Mini-osmotic pumps infused DJNKI-1 or TAT (100□μM) at a rate of 0.15□μl□h−1. After 6 weeks, mice underwent behavioural testing. DJNKI-1 peptide inhibitor of JNK was synthesized by Gene Cust, Custom Service for Research (Laboratoire de Biotechnologie du Luxembourg, Dudelange, Luxembourg).

### Behavioural tests

At 14 weeks of age, mice maintained on a 12 h light/dark cycle and supplied with food and water *ad libitum*, underwent behavioural testing. Tests were conducted 0800 and 1500 h in the order described below with 2-day interval between tests. The experimenter was blind to the treatment. *Elevated plus maze -* Unconditioned anxiety-like behaviour was monitored using an elevated plus maze (EPM) with two closed (35 × 5 × 15 cm) and two open (35 × 5 cm) arms made of opaque grey plastic elevated at a height of 50 cm above the floor. Mice were placed in the centre of the maze facing a closed arm and allowed to explore freely for 5 min. Behaviour was recorded using the Ethovision video tracking system (Noldus Information Technology, Wageningen, The Netherlands). Automated tracking was used to analyse time spent in the open and closed arms. *Light/dark -* test was performed using a box (30 × 45 × 30□cm), partitioned equally, with opaque black plastic walls with a roof (dark arena) and white walls without roof (light arena) and an opening (5.5 × 7□cm) between partitions. To initiate the test, mice were positioned in the light arena and monitored for 10□min. Time spent in the light and dark area was using video tracking with Ethovision software.

### Ethical statement

Animal procedures were conducted with approval from ELLA, the authority responsible for these procedures in Finland, licence number: ESAVI/1897/04.10.07/2015.

### Tissue dissection

Two months of age mice underwent cervical dislocation. The brain was rapidly removed from the skull and transferred to a petri dish containing ice cold PBS, pH 7.4 on ice. Brains were sectioned coronally using an adult mouse brain slicer (Zivic Instruments) at 2.7 and at -4.6 mm from Bregma. NAc (approximately 10 mg) was dissected from the second rostral slice, and the hippocampus from the third. NAc was isolated with inner diameter of 2.0 mm from Bregma 1.54 mm to Bregma 1.04 mm, including both core and shell regions. Afterwards, the third rostral slice was cut by the midline to separate both hemispheres. Hippocampi from each hemisphere, visually recognizable, were isolated. Samples were immediately frozen in liquid nitrogen and stored at -80°C until homogenization within 3 min of sacrifice.

### Immunohistochemistry

Wild type C57Bl/6J and *Jnk1-/-* mice at 8 to 12 weeks and originating from the same genetic background were anesthetized with pentobarbital (Mebunat vet, Orion Pharma, Finland) and transcardialy perfused with 4% paraformaldehyde in phosphate buffered saline (PBS), pH 7.4. Brains were dissected and postfixed in the same solution overnight and then transferred to 20% sucrose in PBS for one day. Thereafter, brains were frozen by immersion in isopentane chilled with CO_2_ ice. Brain sections, 14 µm thick, were prepared using a Leica CM1950 cryostat (Leica Biosystems, Germany) and mounted onto microscope slides (Superfrost Ultra Plus, Menzel-Gläser, Hungary). Slides were dried overnight and stored frozen. After thawing, brain sections were rehydrated in PBS for 10 min. Heat induced antigen retrieval was performed in 10 mM sodium citrate + 0.05% Tween 20, pH 6.0 for 15 min at approximately 100°C. After rinsing with PBS, slides were incubated in 70% ethanol for 30 min followed by 3% H_2_O_2_ + 10% methanol for 30 min. After a quick rinse with PBS, slides were incubated in blocking solution (1% bovine serum albumin + 0.4% Triton X-100 in PBS) for 1 h. Primary antibody (mouse anti-human JNK1 IgG1, clone G151-333, BD Biosciences) was applied at a concentration of 1:15000 in blocking solution for three days at 4°C. After rinsing in PBS, the slides were incubated with secondary antibody (biotinylated anti-mouse IgG, BA-9200, Vector Laboratories, CA, USA) at a concentration of 1:1000 in blocking solution for 1 h. After rinsing in PBS, the slides were incubated with streptavidin-horseradish peroxidase conjugate (S911, Invitrogen Molecular Probes, OR, USA) at a concentration of 1:2000 in blocking solution for 1 h. After rinsing in PBS, the chromogenic signal was developed for 5 min using DAB (Dako Liquid DAB+ Substrate Chromogen System, Dako K3468, Agilent, CA, USA). The slides were rinsed 2 x 2 min in ultrapure water and were then dehydrated in ethanol, cleared in xylene, and cover-slipped. Representative images of wild type and *Jnk1-/-* brain sections were prepared with a Zeiss AxioVert 200M microscope and the Zeiss AxioVision software (Carl Zeiss, Germany). Staining was evaluated in both male and female mice. No difference was observed.

### Primary cell culture

For isolating cortical neurons, newly born Sprague Dawley rats (P0) were decapitated and cortices dissected in dissociated medium containing kynurenic acid in order to prevent activation of AMPA and NMDA receptors. Mechanical dissociation was performed by pipetting after treatment with papain (100 U) (Worthington) and followed by incubation with trypsin inhibitor (10mg/ml) (Sigma) to block papain activity. Cells were cultured on surfaces coated with poly-D-Lysine (50 μg/ml). Neurons were grown in MEM medium supplemented with 10 % Fetal Bovine Serum, 30 mM Glucose, 2 mM Glutamine and Penicillin (50 U/ml)/Streptomycin (50 μg/ml) at a density of 2700 cells/mm^2^. After 48 hours, 2.5 μM cytosine arabinofuranoside (Sigma, St. Louis, MO) was added to the cultures and cells were maintained in a humidified incubator with 5 % CO_2_ at 37°C.

### Cell treatment

Cortical neurons at 16 days *in vitro* were treated with 10 µM TAT or DJNKI-1 peptide (Gene Cust, Custom Services for Research, Laboratoire de Biotechnologie du Luxembourg, Dudelange, Luxemburg) for the times indicated. Cells were lysed in 1x *Laemmli* buffer followed by boiling for 5 min.

### Immunoblotting

Lysates were subjected to sodium dodecyl sulfate (SDS)-polyacrylamide gel electrophoresis. Protein transfer was performed using polyvinylidene fluoride (PVDF) membranes (Bio-Rad). Membranes were incubated with 1x PBS-T (PBS containing 0.05 % Tween-20) and 5 % skimmed milk, for 1 h at room temperature. Primary antibody incubation was performed overnight at 4°C. The following day, membranes were washed with 1x PBST and incubated with horse-radish peroxidase labelled goat anti-mouse or rabbit IgG (Millipore) (1:50.000) for 1 h at room temperature. Bands were exposed to SuperSignal West Femto Maximum Sensitivity Substrate (Thermo Scientific) and visualized with ChemiDocTM MP Imaging System (Bio-rad). Densitometry analysis was performed with ImageJ (NIH). Primary antibody dilutions were as follows: phospho-GSK3-β Ser9 (1:1000, (#9336S, Cell Signalling), GSK3-α (1:1000, #sc-5264, Santa Cruz), phospho-AKT-Ser473 (1:1000, #9271, Cell Signaling) and AKT1 (1:1000, #sc-1618, Santa Cruz).

### Brain homogenization and protein extraction

Hippocampi and NAc underwent heat stabilization using Stabilizor™ T1 (Denator AB, Sweden). An amount 10 times more than the weight of the sample of Lysis buffer (8M Urea/ 100 mM AMBIC) was added to each sample supplemented with phosphatase and protease inhibitors (Sigma-Aldrich, St. Louis, USA). Tissue was sonicated for 20 s on ice in small PCT MicroTubes for Barocycler NEP2320 (Pressure BioSciences, USA). Homogenized samples were processed according to Pressure Cycling Technology using Barocycler NEP 2320 (Pressure BioSciences, USA), using a program of 50 s high pressure (35,000 p.s.i.) and 10 s ambient pressure (14.7 p.s.i.) for 60 cycles to guarantee the total disruption of the membranes and efficient protein extraction (Powell B.S. et al 2012; Olszowy et al 2013). The solution was then transferred into a 1.5 ml tube and centrifuged at 16000 x g for 1 h at 4°C. Supernatant, containing only proteins, was transferred to a clean tube and protein quantification was performed using Total Protein Kit, Micro-Lowry, Petersońs Modification (Sigma-Aldrich, St. Louis, USA).

### SDS-page and protein digestion

All the chemicals for Mass Spectrometry analysis are from Sigma-Aldrich (St. Louis, USA). A total amount of 200 μg of protein extract from Hippocampi and NAc were loaded onto 12 % TGX Criterion Gel (Bio-Rad Laboratories, Hercules, Ca, USA) for in-gel proteomic analysis. Proteins were electrophoresed in the gel for 20 minutes. Gels were washed in milliQ water, stained for 30 minutes with GelCode (Bio-Rad Laboratories, Hercules, Ca, USA) and de-stained overnight in milliQ water. Each lane was then manually sliced. Gel slices were de-stained 3 times in 25 mM AMBIC/50% ACN and dried in a vacuum centrifuge (Speedvac, Thermo Scientific, Germany). Samples were reduced (10 mM DTT/100 mM AMBIC, 1 h at 56°C), alkylated (55mM IAA/100 mM AMBIC, 45 min at room temperature in the dark) and washed twice with 100 mM AMBIC and dehydrated using ACN. Samples were dried in the Speedvac. Slices were rehydrated with 12.5 μg/ml modified porcine trypsin (Promega, Madison, WI, USA) freshly diluted in 50 mM AMBIC and the digestion was performed overnight at 37°C. Peptides were eluted twice using 75 % ACN/ 5% FA, dried in the Speedvac (Thermo Fisher Scientific) until complete dryness and dissolved in 0.1 % FA before further analysis.

### Phospho-peptide enrichment

Peptides were phospho-enriched using TiO_2_ beads (TiO_2_ Mag Sepharose, GE Healthcare, Little Chalfont, Buckinghamshire,UK) according to manufacturer’s instructions and dried in the Speedvac (Thermo Fisher Scientific) until completely dry.

### LC-MS/MS and data acquisition

Dry peptides were re-suspended in 0.1 % FA and separated using a 2D Eksigent nano-LC system (Eksigent, Dublin, CA, USA) coupled to an LTQ-Orbitrap XL mass spectrometer (Thermo Fisher Scientific) operated in DDA mode. Unassigned charge states and singly charged ions were excluded from fragmentation. The dynamic exclusion list was limited to a maximum of 500 masses with a maximum retention time of 2 minutes and a relative mass window of 10 ppm. Xcalibur software version 2.0.7 (Thermo Fisher Scientific, Germany) was controlling the HPLC, mass spectrometer and data acquisition.

### Mass spectrometry data analysis and data pre-processing

LC-MS/MS raw files were analysed in MaxQuant (Cox 2008, PMID: 19029910) 1.6.17.0 against Mus musculus FASTA file with isoforms downloaded from UniProt(Uniprot in 2023 pmid 36408920). Cysteine carbamidomethylation was set to fixed modification, and methionine oxidation, n-terminal protein acetylation and serine, threonine, tyrosine phosphorylation were set as variable modifications. MaxQuant’s Match between run function was turned on. The resulting phospho (STY) table was processed in Perseus 1.6.15.0 (The Perseus computational platform for comprehensive analysis of (prote)omics data pmid 27348712). Rows that did not have 50% non-missing values in at least one condition were filtered away. Reverse and potential contaminant sites were filtered away. Raw site intensity values were log2 transformed and column median was subtracted from each sample, and median of all values before subtraction was added to all values to bring the values back to original scale. Missing values were imputed with Perseus replace values from the normal distribution function in total matrix mode. The values were back transformed for subsequent bioinformatics analyses. Data are available via ProteomeXchange (PRIDE database) with identifier PXD060057.

### Bioinformatics analysis

The MS/MS data was analysed using the PhosPiR analysis pipeline (Hong et al., 2022). PhosPiR is an automated proteomics/phosphoproteomics pipeline written in R, which includes a complete set of analysis tools ranging from data preprocessing to downstream analysis such as enrichment analysis and network analysis. We compared the phosphosite intensities from hippocampus and nucleus accumbens of mice treated with TAT or DJNKI-1 using PhosPiR’s statistical analysis package. For visualization, a circosplot of significantly changing phosphosites annotated according to functional groups was created using the “circlize” package in R (Gu et al., 2014).

### Pearson’s correlation analysis

Correlation plots are made for MS data and behaviour data. Data was separated into four groups, hippocampus treated with TAT (n=6), hippocampus treated with DJNKI-1 (n=6), nucleus accumbens treated with TAT (n=4), and nucleus accumbens treated with DJNKI-1 (n=3). MS data consisted of phosphosite intensity values, and behaviour data consisted of time scores in seconds for the EPM and light/dark behaviour tests. Pearson correlation was performed to explore the correlative relationship between behaviour and phosphorylation changes for treatment groups. Correlations between proteins for each treatment group was also compared. Pearsons’s correlation coefficients and corresponding p-values were calculated using R packages (“*stats”* and “*ggcorrplot”*, respectively). Correlation of behaviour data verses MS data was plotted using the *ggcorrplot* package, where a threshold of inclusion was set at correlation value of >0.5 in absolute value, and p-value of <0.05 for correlation. Phosphosites that had at least 1 correlation fitting these criteria with any of the 5 behaviour scores were plotted. For these phosphosites, all behaviour correlations were included, even if the correlation coefficient was lower than 0.5. Correlation of protein verses protein was plotted using the “*corrplot”* package. The same significance threshold was applied. Only significant correlations are represented (by colored circle) in the plot. The phosphoprotein cross-correlations were ordered using spectral clustering performed using the “spectrum” package in R. Normalized graph Laplacian was calculated from the correlation similarity matrix to generate the eigensystem, then k-means was applied on the top k eigenvectors to get the spectral clusters.

### Responder/non-responder split

From the DJNKI-1 treatment group, we further separated the data into responder and non-responder groups based on the light dark test results. Mice with time spent in the light measuring greater than 1 standard deviation above the mean of TAT-treated mice, were considered responders, while the rest JNKI-1 treated mice were considered non-responders. This left 6 control TAT treated mice and 3 mice in each of the categories responder and non-responder. Separate correlation to light/dark behaviour features and correlation p-values are calculated for responders and non-responders. A volcano plot was created with the “ggplot2” package in R to visualize the correlation values and correlation p-value of each phosphoprotein from both responders and non-responders. It should be noted that when the responder/non-responder split analysis was performed on the EPM test results, the split was 2 responders to 4 non-responders, therefore the correlations shown are only valid for correlation with a low anxiety phenotype in the light dark test.

### Motif analysis

Potential GSK3 phoshorylation sites were identified with Perseus motif linear motif analysis tool. Consensus motif S/TX_(2-3)_S/T,X/P was used to find potential GSK3 sites that were significantly changing (FC > 1.5 and ROTS p-value < 0.05). Potential GSK3 sites were required to have a second significant (FC > 1.5 and ROTS p-value < 0.05) co-detected on same peptides.

## Results

### Intracerebral infusion with JNK inhibitor induces phosphorylation changes in mouse hippocampus and nucleus accumbens

We previously demonstrated that intracerebral infusion of the DJNKI-1 peptide JNK inhibitor (Borsello and Bonny, 2004), induces anxiolytic-like behaviour in mice after six weeks (Mohammad et al., 2018). Here, we set out to determine the phosphoproteome changes elicited by this treatment in mouse brain. We first characterized the localization of constitutively active JNK1 using a JNK1-specific antibody (Coffey et al., 2000; Coffey et al., 2002). JNK1 was detected throughout the cortex and subcortical regions, with the most prominent signal in the hippocampus formation, particularly in the neuropil. Importantly, this signal was absent in *Jnk1* knockout mice, confirming its specificity (Fig. 1A, B). Based on the high level of JNK1 in hippocampus (HPC), a region integral to emotional regulation, we focused our analysis there. The nucleus accumbens (NAc), a key region for reward and motivation, was included for comparison.

**Figure 1.**
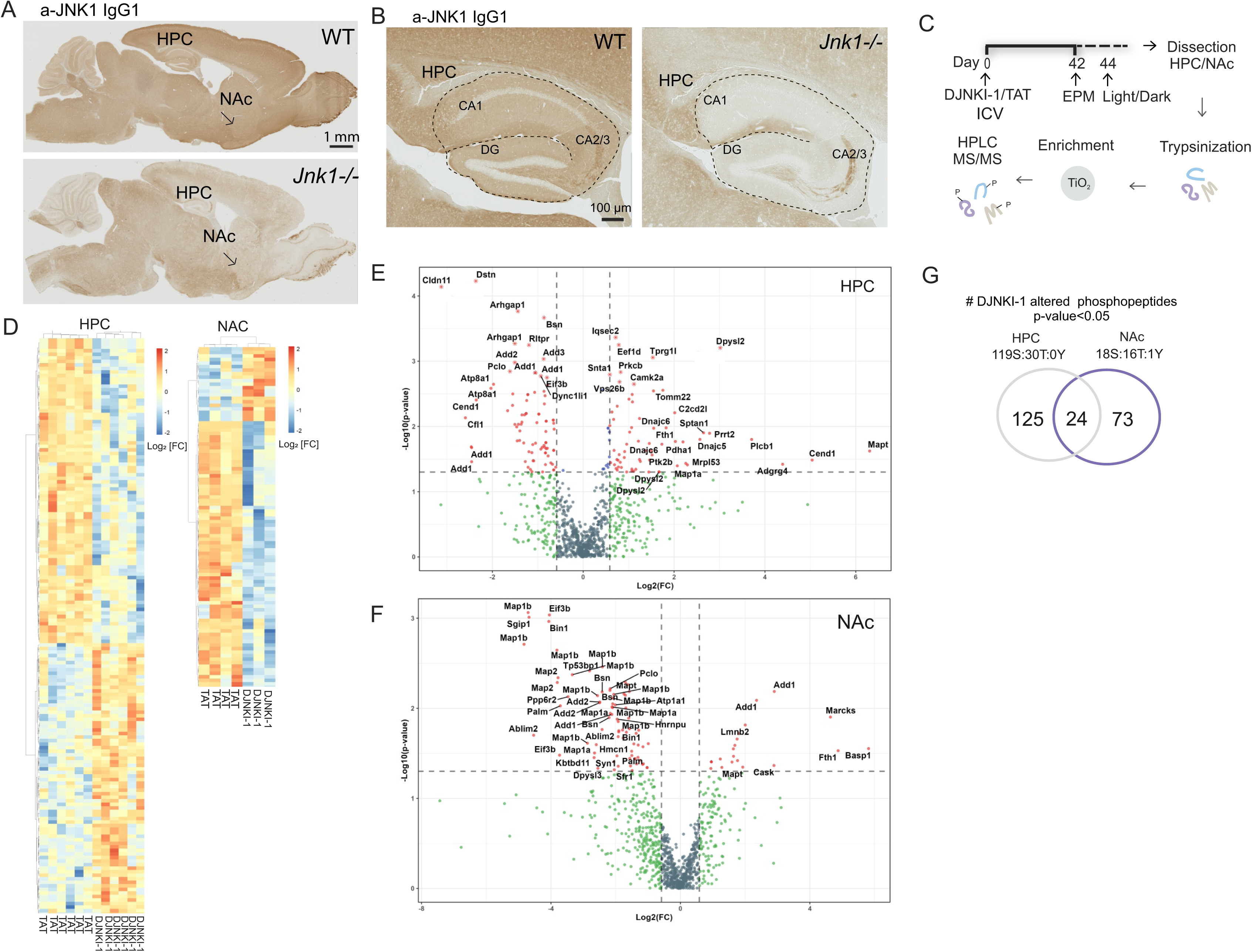
ICV infusion with DJNKI-1 induces phosphorylation changes in hippocampus and nucleus accumbens. A, B) Sagittal sections from wild type (WT) and *Jnk1 -/-* mouse brains stained with antibody against JNK1. Immunohistochemistry reveals the expression of JNK1 in WT brain. Signal specificity is validated using *Jnk1-/-* brain. C) Experimental timeline: mice were infused with DJNKI-1 or TAT for 6 wk followed by EPM and light/dark behaviour testing. Hippocampus (HPC) and nucleus accumbens (NAc) were dissected and underwent phosphoproteomic investigation. D) Euclidean distance and ward.D2 cluster analysis heatmaps show Log_2_-normalised phosphosite intensities for significant changes (ROTS p-value p<0.05, no fold cut-off). E, F) Volcano plots of phosphosite intensity changes in HPC and NAc following DJNKI-1 treatment. G) The number of significantly changing phosphosites with F.C.>2.0 and p-value < 0.05 are shown for HPC and NAc. HPC: TAT: n=6, DJNKI-1: n=6; NAc: TAT: n=4, DJNKI-1: n=3.

We next infused mice for six weeks with either DJNKI-1 or the TAT delivery peptide as a control, using mini-pumps, as previously described (Mohammad et al., 2018). Mice then underwent EPM and light/dark tests to assess anxiety-like behaviours. The HPC and NAc were then isolated for phosphopeptide enrichment and LC-MS/MS analysis (Fig. 1C). In total, 994 phosphosites were identified, accounting for multiplicity. From these 149 phosphosites were regulated by DJNKI-1 in the HPC (F.C. >1.5, p<0.05), and 97 in the NAc (Fig. 1D-F; Supplementary Table 1). Notably, only 24 of the regulated phosphosites overlapped between the two regions (Fig. 1G, Table 1). These included presynaptic regulators of neurotransmitter release (e.g. basoon and piccolo), proteins controlling lipid dynamics (e.g. phospholipid transporting ATPase 1A, paralemmin-1), and cytoskeletal proteins (e.g. MAP2, MAP1a, DPYSL3, ADD1 and 3).

**Table 1.**
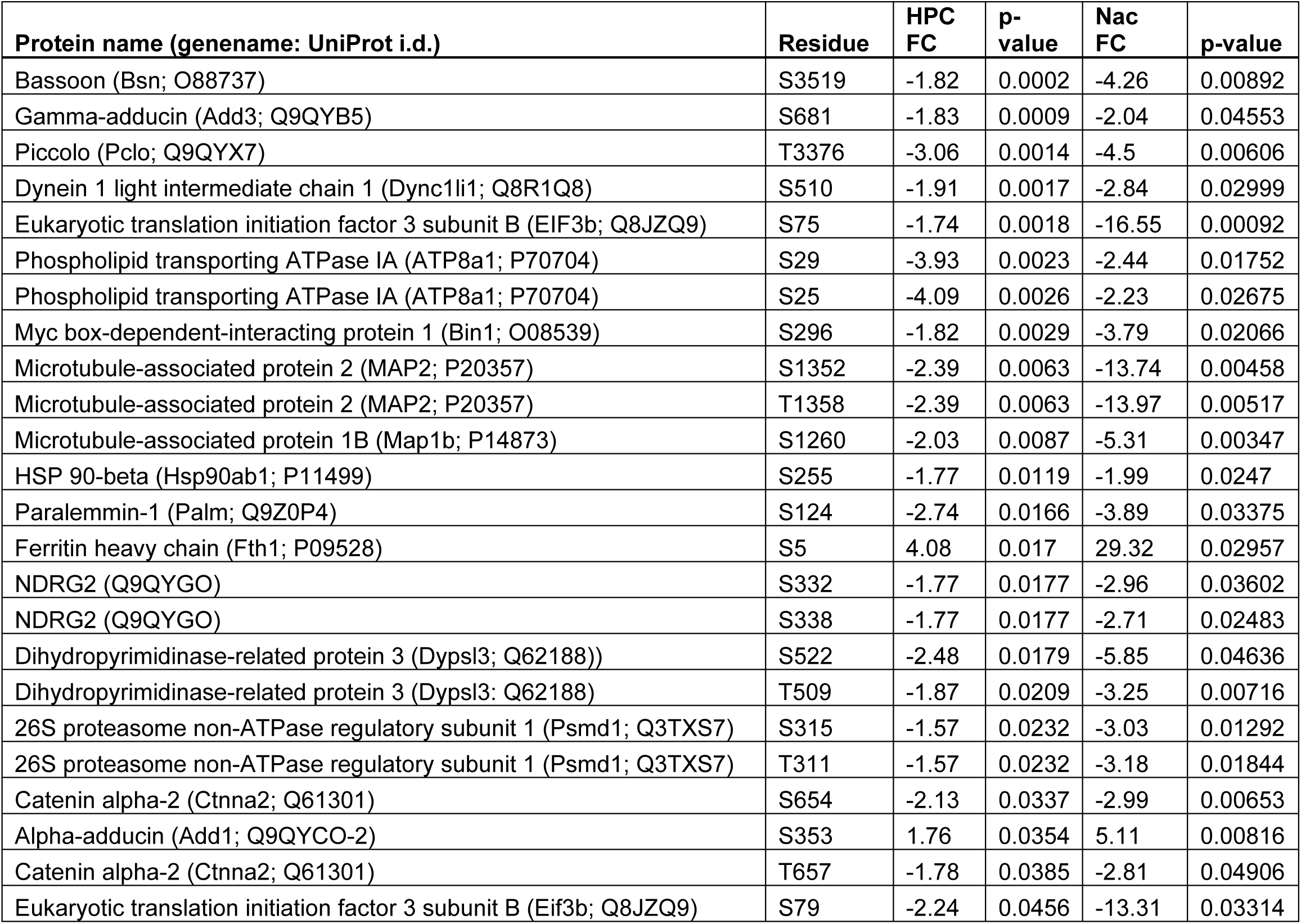
Overlapping phosphorylation sites that changed significantly both in hippocampus (HPC) and nucleus accumbens (NAc).

| **Protein name (genename: UniProt i.d.)** | **Residue** | **HPC**<br>**FC** | **p-**<br>**value** | **Nac**<br>**FC** | **p-value** |
| --- | --- | --- | --- | --- | --- |
| Bassoon (Bsn; O88737) | S3519 | -1.82 | 0.0002 | -4.26 | 0.00892 |
| Gamma-adducin (Add3; Q9QYB5) | S681 | -1.83 | 0.0009 | -2.04 | 0.04553 |
| Piccolo (Pclo; Q9QYX7) | T3376 | -3.06 | 0.0014 | -4.5 | 0.00606 |
| Dynein 1 light intermediate chain 1 (Dync1li1; Q8R1Q8) | S510 | -1.91 | 0.0017 | -2.84 | 0.02999 |
| Eukaryotic translation initiation factor 3 subunit B (EIF3b; Q8JZQ9) | S75 | -1.74 | 0.0018 | -16.55 | 0.00092 |
| Phospholipid transporting ATPase 1A (ATP8a1; P70704) | S29 | -3.93 | 0.0023 | -2.44 | 0.01752 |
| Phospholipid transporting ATPase 1A (ATP8a1; P70704) | S25 | -4.09 | 0.0026 | -2.23 | 0.02675 |
| Myc box-dependent-interacting protein 1 (Bin1; O08539) | S296 | -1.82 | 0.0029 | -3.79 | 0.02066 |
| Microtubule-associated protein 2 (MAP2; P20357) | S1352 | -2.39 | 0.0063 | -13.74 | 0.00458 |
| Microtubule-associated protein 2 (MAP2; P20357) | T1358 | -2.39 | 0.0063 | -13.97 | 0.00517 |
| Microtubule-associated protein 1B (Map1b; P14873) | S1260 | -2.03 | 0.0087 | -5.31 | 0.00347 |
| HSP 90-beta (Hsp90ab1; P11499) | S255 | -1.77 | 0.0119 | -1.99 | 0.0247 |
| Paralemmin-1 (Palm; Q9Z0P4) | S124 | -2.74 | 0.0166 | -3.89 | 0.03375 |
| Ferritin heavy chain (Fth1; P09528) | S5 | 4.08 | 0.017 | 29.32 | 0.02957 |
| NDRG2 (Q9QYGO) | S332 | -1.77 | 0.0177 | -2.96 | 0.03602 |
| NDRG2 (Q9QYGO) | S338 | -1.77 | 0.0177 | -2.71 | 0.02483 |
| Dihydropyrimidinase-related protein 3 (Dypsl3; Q62188) | S522 | -2.48 | 0.0179 | -5.85 | 0.04636 |
| Dihydropyrimidinase-related protein 3 (Dypsl3; Q62188) | T509 | -1.87 | 0.0209 | -3.25 | 0.00716 |
| 26S proteasome non-ATPase regulatory subunit 1 (Psmd1; Q3TXS7) | S315 | -1.57 | 0.0232 | -3.03 | 0.01292 |
| 26S proteasome non-ATPase regulatory subunit 1 (Psmd1; Q3TXS7) | T311 | -1.57 | 0.0232 | -3.18 | 0.01844 |
| Catenin alpha-2 (Ctnna2; Q61301) | S654 | -2.13 | 0.0337 | -2.99 | 0.00653 |
| Alpha-adducin (Add1; Q9QYCO-2) | S353 | 1.76 | 0.0354 | 5.11 | 0.00816 |
| Catenin alpha-2 (Ctnna2; Q61301) | T657 | -1.78 | 0.0385 | -2.81 | 0.04906 |
| Eukaryotic translation initiation factor 3 subunit B (Eif3b; Q8JZQ9) | S79 | -2.24 | 0.0456 | -13.31 | 0.03314 |

Some of these phosphorylation sites were linked to known regulatory functions. For example, phosphorylation of NDRG2-S322 by SGK1 primes it for phosphorylation by GSK3 (Murray et al., 2004). Similarly, DPYSL3 (CRMP4) phosphorylation at S522 by GSK-3 influences neurite length (Cole et al., 2006) (Table 1). These findings indicate that DJNKI-1 modulates neuronal cytoskeleton and synaptic function. Other sites with known function are described below. We plotted the top 20 most significant phosphoprotein changes per region in a Circos plot according to functional category (Fig. 2). Cytoskeletal proteins, particularly the microtubule stabilizers MAP1a, MAP1b and MAP2, showed large fold phosphorylation changes. MAP1b-S1260, a site critical for axon collateral branching (Scales et al., 2009; Ziak et al.) was significantly altered in both NAc and HPC, while 13 additional sites on MAP1b were regulated exclusively in NAc, despite being detectable in HPC (Fig. 2A, Supplementary Table 1). MAP2 phosphorylation decreased at S1352 and T1358 in both regions, with a more pronounced reduction in NAc (-13-fold, p= 0.004 and 0.005) compared to HPC (-2.4-fold, p= 0.006). Similarly, Tau (MAPT) phosphorylation on S692 (human S717) decreased 4-fold in NAc and 2-fold in HPC. S717 Tau, a site that is also subject to o-glycosylation, is known to influence fibril formation, a hallmark of Alzheimer’s disease (Liu et al., 2004; Pratt and Vocadlo, 2023).

**Figure 2.**
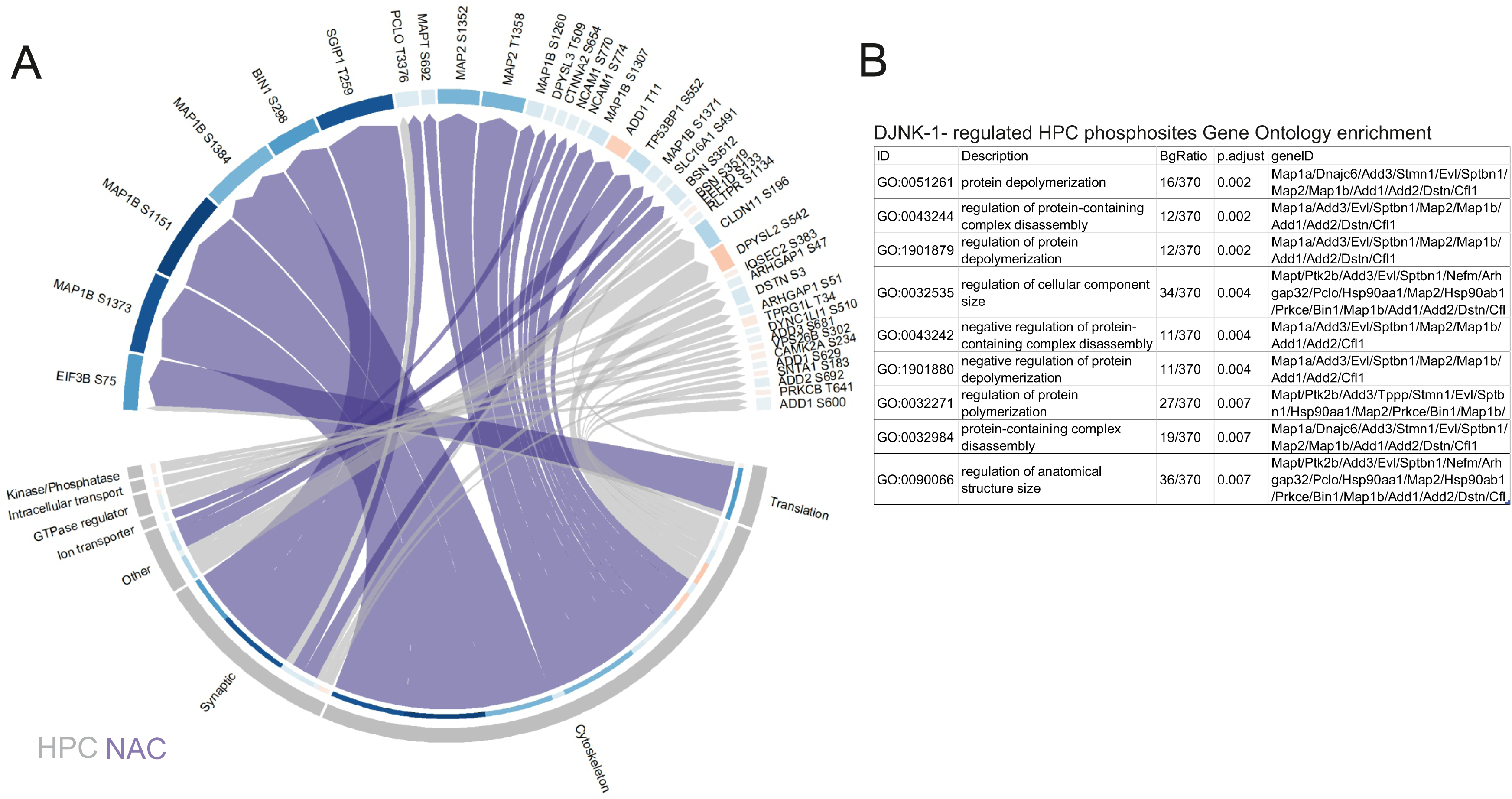
Inhibition of JNK leads to prominent decreases in cytoskeletal protein phosphorylation in both hippocampus and nucleus accumbens. A) Circos plot depicts significantly altered phosphosites with F.C. >2.0, F.D.R.< 0.05. DJNKI-1-regulated phosphoprotein sites are grouped by brain region (HPC=grey, NAc=purple) and by function. Link size and phosphoprotein sector color correspond with phosphorylation intensity fold change (blue represents decreasing phosphosite intensity and red increasing). Results shown are from the top 20 most significant changes in each brain region. For conciseness, gene names are used. Note DSTN (dystrophin) is otherwise known as cofilin-1 and Bin1 (bridging integrator 1) is otherwise known as amphiphysin-2. B) Gene ontology enrichment of significantly changing phosphoproteins in HPC of DJNKI-1-treated mice.

In HPC, kinase and phosphatase phosphorylation was notably regulated (Fig. 2; Supplementary Table 1). Significant changes were observed in PRKCB, AAK1, CamK2a, PRKCE, PDPK1, PRTK2b and PP1r7. Among these, PDPK1-S241 phosphorylation, a critical activation loop site that controls its enzymatic activity (Levina et al., 2022), increased 3-fold (p=0.0003), indicating enhanced PDPK1 activity. In contrast, no kinases or phosphatases were significantly altered in NAc. However, several synaptic proteins were regulated in both regions. These included bassoon (Bsn)-S3519 and piccolo (Pclo)-T3376. Synapsin 1 (Syn1), where S427 regulated in HPC, and S568 in NAc (Fig. 2A, Supplementary Table 1).

A gene ontology enrichment analysis highlighted changes related to “protein depolymerization and regulation of protein-containing complex disassembly” in the hippocampal DJNKI-1-regulated phosphoproteome (Fig. 2B), with no significant enrichments obtained in NAc. The Circos plot (Fig. 2A), shows the top 20 regulated phosphoproteins, while additional changes were detected in cytoskeletal regulatory proteins in HPC contributing to this enrichment (Supplementary Table 1).

### Protein phosphorylation changes in HPC and NAc correlate with high and low anxiety-like behaviours

We next examined the behaviour of DJNKI-1-treated mice in the EPM and light/dark tests, to replicate the anxiolytic effects observed previously with DJNKI-1 and in *Jnk1*-/-mice (Mohammad et al., 2018). As before, DJNKI-1-infused mice spent more time in the open arms of the EPM and the light area in the light/dark test (Fig. 3A, B). Thus DJNKI-1 treatment induced a similar low anxiety-like phenotype as reported earlier.

**Figure 3.**
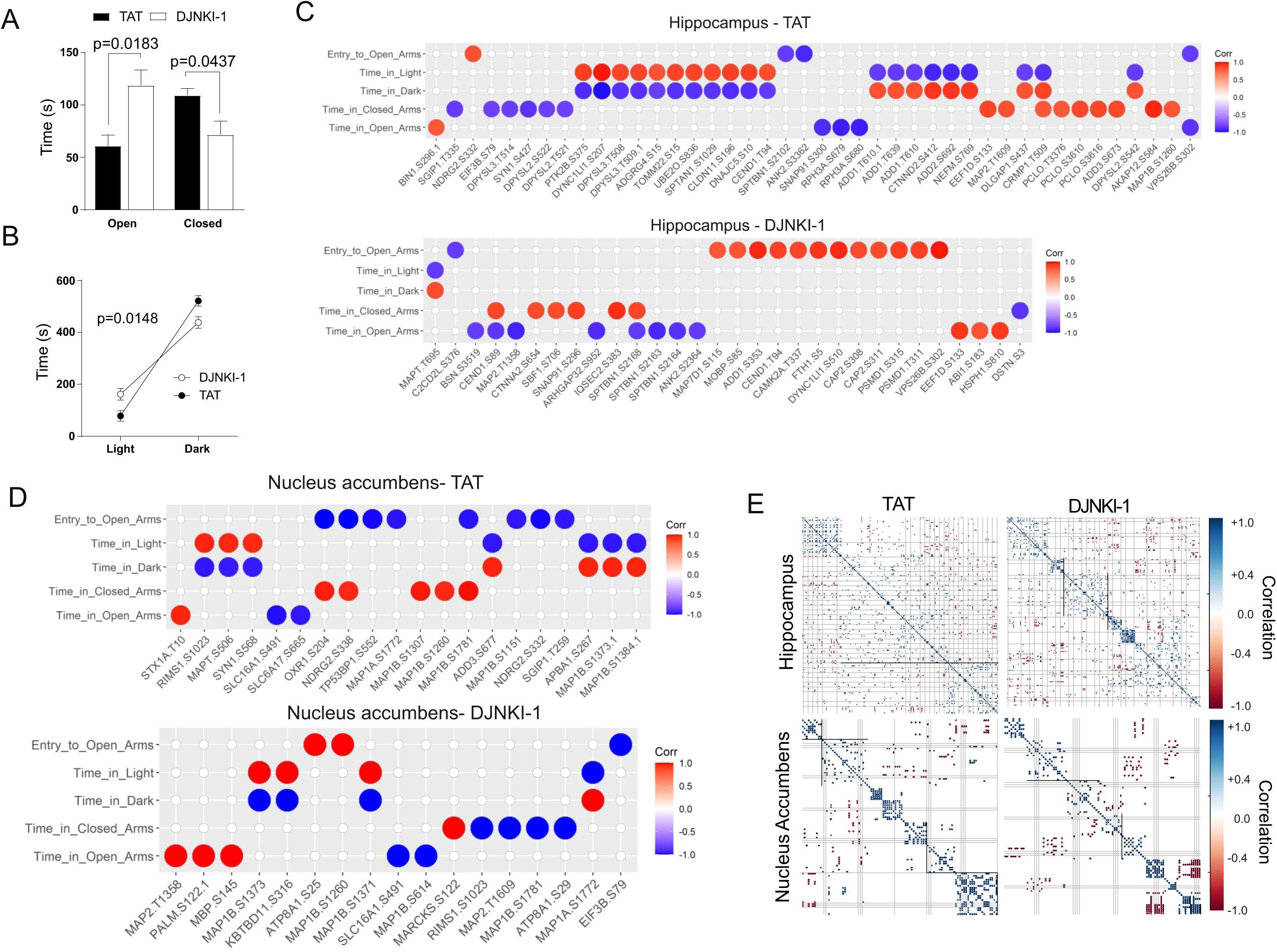
DJNKI-1 treated mice show reduced anxiety like behaviours that correlate significantly with multiple phosphorylation events in hippocampus. A-B) Behaviour results from the EPM and light-dark test following 6 weeks treatment with TAT (control peptide) (n=8) or DJNKI-1 (n=7). P-values were calculated using Student’s T-test. C) Dot plots show the correlation between behaviours and phosphosite intensities, F.C.>2, p-value <0.05 in HPC and NAc. Pearson’s coefficients are depicted with red or blue circles for positive and negative correlation respectively. Only correlations > 0.5 and p-value < 0.05 are included in the plots. E) Spectral clustering maps depict cross-correlating phosphoprotein sites in HPC and NAc. Only significant correlations (p<0.05) are included. A difference in cluster pattern between TAT- and DJNKI-1-treated mice is observed. Nodes of cross-correlating phosphoproteins change in both HPC and NAc following DJNKI-1 treatment.

To identify which phosphosites were likely to affect the behaviours, we correlated phosphosite intensities with time spent in the aversive and non-aversive zones in the behaviour tests. In control mice, several phosphosites in HPC, including those on tubulin regulatory proteins DPYSL1, 2 and 3 (CRMP1, 2 and 4), as well as adducin 1, correlated bidirectionally with behaviour (Fig. 3C, TAT). Conversely, no such bidirectional correlations were found in the EPM of control mice.

In DJNKI-1-treated mice, an entirely new set of correlations emerged. Altogether 30 phosphorylation sites correlated with time spent in the open/closed arms, and entries to the open arms (Fig. 3C, HPC-DJNKI-1). The correlating phosphosites included axonal proteins regulating actin and microtubules (e.g. MAPT-S695, MAP2-T1358, ANK2-S2364 and SPTBN1-S2164), and synaptic proteins (e.g. BSN-S3519, DSTN-S3 and SNAP91-S296). Notably, destrin (DSTN or cofilin-1)-S3 phosphorylation correlated negatively with time spent in the closed arms of the EPM (Fig. 3C, DJNKI-1). Phosphorylation on this site decreased by 5-fold (p=0.00005) in DJNKI-1 treated mice and was the most significant change in the dataset. Cofilin-S3 phosphorylation has been shown to promote actin filament growth (Mizuno, 2013). Remarkably, in the light/dark test, most correlations disappeared, and only MAPT-T695 correlation remained. It correlated positively and negatively with time in the dark and light respectively (Fig. 3C, HPC-DJNKI-1).

In NAc of control peptide-treated mice, fewer significant correlations were observed overall. Notably, MAP2-T1358 phosphorylation correlated with time spent in the open arms in both brain regions after DJNKI-1 treatment, but the correlation was positive in NAc and negative in HPC. We applied spectral clustering to visualize phosphorylation synchronization with or without JNK inhibitor (Fig. 3E). In hippocampus, drug treatment increased the number of clusters of positively correlating phosphoproteins (Fig. 3E), possibly reflecting greater synchronization of signaling in HPC when JNK is inhibited.

### Hippocampus responder/non-responder analysis

Given that only 50% of DJNKI-1-treated mice showed an anxiolytic response, we reanalyzed the phosphoproteomic data for responder and non-responder mice separately, to identify phosphorylation changes that may determine behavioral outcome (Fig. 4). Volcano plots depicting phosphorylation changes in the hippocampus of mice that responded to the light/dark test with lower anxiety (responders), and those that did not (non-responders) are shown (Fig. 4A, B; Supplementary Table 2). Interestingly, only NCAM1-S774 changed significantly and in opposite directions in both the responder and non-responder group. It decreased by 3.4-fold (p=0.009) in responders and increased by 2.2-fold (p=0.0004) in non-responders. This site is known to regulate NCAM1 interaction with CRMP2 which in turn, promotes axonal growth (Karbe et al., 2013). There were several unique phosphorylation changes in responder and non-responder mice. Among these, phosphorylation of adducin-1 T610 and T639 decreased substantially (-20 and -17 fold respectively) only in responder mice. On the other hand, MAPT T695 (human T720) increased 125-fold only in non-responder mice. Similarly, Hsp90ab1 S255 phosphorylation decreased -2.5-fold only in non-responders. Phosphorylation on this site is required for apoptosome formation during apoptotic cell death (Kurokawa et al., 2008). Interestingly also, pyruvate dehydrogenase (PDHA1)-S293 phosphorylation increased by 6-fold in non-responder mice only, 5-fold in the mixed population. This site inhibits its enzymatic activity. Thus DJNKI-1 treatment is predicted to prevent pyruvate from entering the TCA cycle, and divert its use towards lactate production. Interestingly, phosphorylation on this site inversely correlates with neuronal activity (Yang et al., 2024).

**Figure 4.**
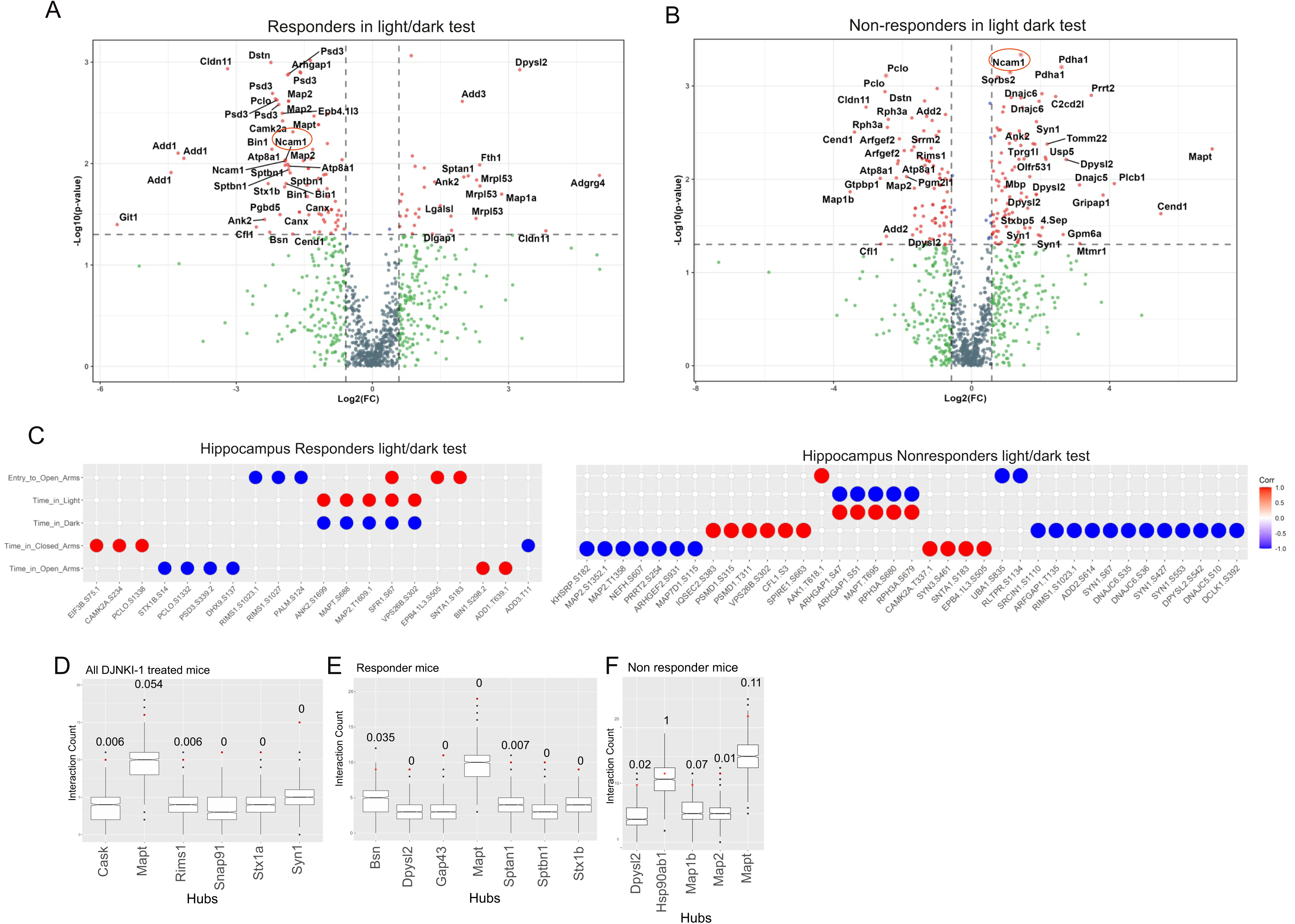
DJNKI-1-induced phosphorylation changes in responder and non-responder mice. A, B) Volcano plots depict phosphosite changes in responder and non-responder mice compared to control. N=6 for TAT-treated mice and n=3 respectively for responder and non-responders. C) Correlations plots of significantly changing (p<0.05 using ROTS) phosphosite intensities against behavioural metrics (number of entries to open arms, time in closed or open arms) for the EPM test and (time in the light/dark zones) for the light/dark test are shown. Pearson’s correlations for phosphosites that significantly change > 2-fold with a p-value <0.05 using the ROTS test are indicated. C) Protein phosphorylation changes that significantly correlate with either low or high anxiety behaviours in light-dark or EPM test are shown. Pearson’s r is plotted against –Log_2_ p-value. The phosphopeptide list showing corresponding phosphosite information can be found in Supplementary Table 2. D-F) Signalling hub analysis shows the most highly interacting signalling nodes in the phosphoproteomes of DJNKI-s-treated mice (D), responder mice (E) and non-responder mice (F). Horizontal black lines depict the median values for number of interacting proteins based on bootstrapping against all phosphoproteins identified in HPC. The red spot indicates the number of putative interacting proteins in the comparison group and the corresponding p-value.

Phosphorylation changes that correlated with anxiety responses were enriched for presynaptic proteins (Fig. 4C). For example, in responders, piccolo, amphiphysin-2, adducin 1 and spectrin-1 were regulated at multiple sites (Fig. 4A). Also, phosphorylation of STX1b decreased on S14, a site known to regulate neurotransmitter release (Shekar et al., 2023). In non-responders, phosphorylation of nerve terminal and cytoskeletal proteins (e.g. synapsin I/III, NEFH, MAP2, MAPT, CFL1, DPYSL2) correlated with light/dark test behaviour (Fig. 4C). These findings suggest presynaptic regulators of neurotransmitter release are likely drivers of anxiety-liked behaviours.

### Signalling hub connectivity increases in hippocampus of DJNKI-1 treated mice

To explore the connectivity of DJNKI-1-regulated phosphoproteins, we created interaction networks using the STRING database. Hubs were defined as phosphoproteins with connectivity greater than two standard deviations above the mean interaction count. To validate hub specificity, we generated 1,000 random background networks of equivalent size and compared interaction counts to identify statistically significant hubs. Our analysis suggests that CASK, MAPT, RIMS1, SNAP91, STX1A, and SYN1 function as hubs in the phosphoprotein network of DJNKI-1-treated mice (Fig. 4D-F).

Interestingly, when focusing exclusively on responder mice, a distinct set of hubs with significant connectivity emerged. These included DPYSL2 (CRMP2), GAP43, MAPT, spectrin beta (SPTBN1), spectrin alpha (SPTAN1), and STX1B (syntaxin 1b). This pattern suggests more robust hub activity in responder mice following DJNKI-1. These hubs may potentially drive the signalling underlying the low-anxiety phenotype. In contrast, there was an absence of specific signalling hubs in the non-responder group, indicating minimal changes in phosphoprotein connectivity relative to the random background networks.

### The AKT-GSK3 pathway is regulated in JNK inhibitor-treated mice

We previously described the phosphoproteome of *Jnk1-/-* mouse brain (Hong et al., 2024), identifying that JNK1 regulates AKT1 and GSK signaling hubs that contribute to anxiolytic behaviours (Hong et al., 2024). We therefore decided to examine GSK3 phosphorylation in our dataset, as GSK3 is independently implicated in anxiety regulation (Crofton et al., 2017; Mines et al., 2010),

GSK3 sites were predicted from the DJNKI-1-regulated hippocampal phosphoproteome using the motif S/T_X(2-3)_S/T, where the priming site is classically C-terminal to the GSK3 site (Fiol et al., 1987). We required both sites to be significantly regulated by >1.5 fold in the same direction. The results, shown in Table 2 reveal that all predicted GSK3 sites underwent negative fold-changes in DJNKI-1-treated mice, along with their predicted priming sites. In total, 18 GSK3 motifs showed decreased phosphorylation in hippocampus and 10 in nucleus accumbens (Table 2). Notably, these changes occurred in responder and non-responder mice, however only 30% occurred in both regions simultaneously. These findings suggest that GSK3 signaling is inhibited in these brain regions following DJNKI-1 treatment.

**Table 2.**
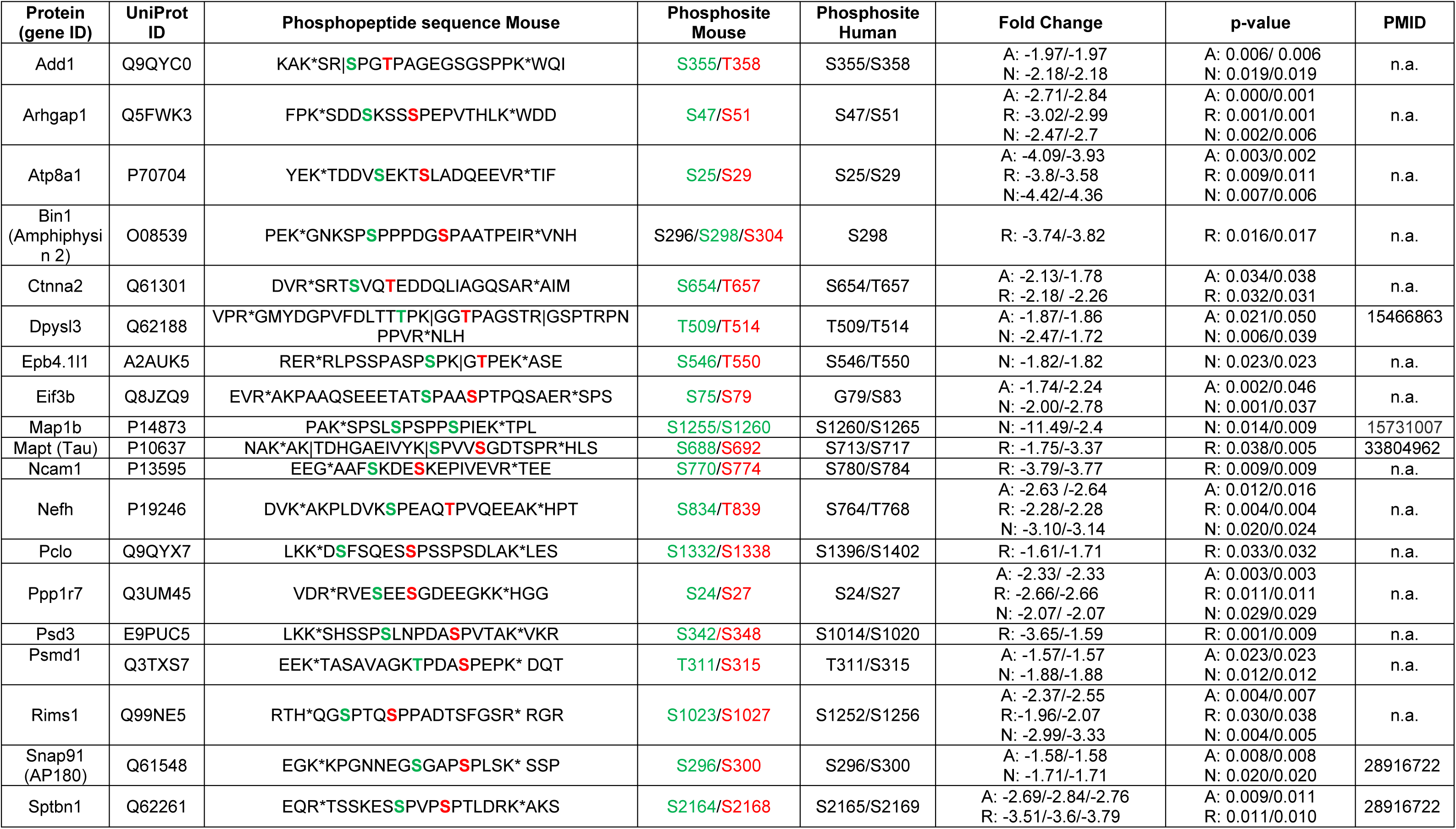
Potential GSK3 motifs identified from hippocampus of DJNKI-1-treated mice. A “relaxed” GSK3 consensus motif S/TX_(2-3)_S/T,X/P search was applied. Significantly changing phosphosites (FC>1.5, p<0.05) are shown only in cases where both priming site and GSK3 site (red and green) are altered relative to control, except for MAP1b and MAPT which are reportedly unprimed. Sites co-detected by MaxQuant are shown when significant and are bolded black. Miss-cleavage sites are indicated by |. FC and p-values are shown for site in all mice “A”, responder mice “R” and non-responder mice “N”. UniProt ID and corresponding gene names are given. Note, only sites that fit the motif are shown. In some cases, additional sites on a shown peptide underwent significant change. All phosphosite changes can be seen in supplementary Table 1.

### GSK3β is inhibited by JNK inhibitor treatment or Jnk1 deletion in HPC and NAc

We assessed whether DJNKI-1 treatment altered GSK3 activity using phospho-antibodies recognising the activation state of GSK3a and β. This revealed that 8 hours treatment of cortical neurons with DJNKI-1 (10 µM) led to inhibition of both GSK3 isoforms (Fig. 5A-C). We also measured the activity of AKT, which inhibits GSK3 by direct phosphorylation of S9 and S21 of GSK3b and a respectively. AKT-S473 phosphorylation increased, indicating activation (Fig. 5A, D). Similar regulation of GSK3□ and AKT activity was observed in the NAc and hippocampus of *Jnk1-/-* mice, though in hippocampus, only GSK3b was regulated (Fig. 5E-K). These results indicate that inhibition of JNK in mouse brain activates AKT and inhibits GSK3 activities.

**Figure 5.**
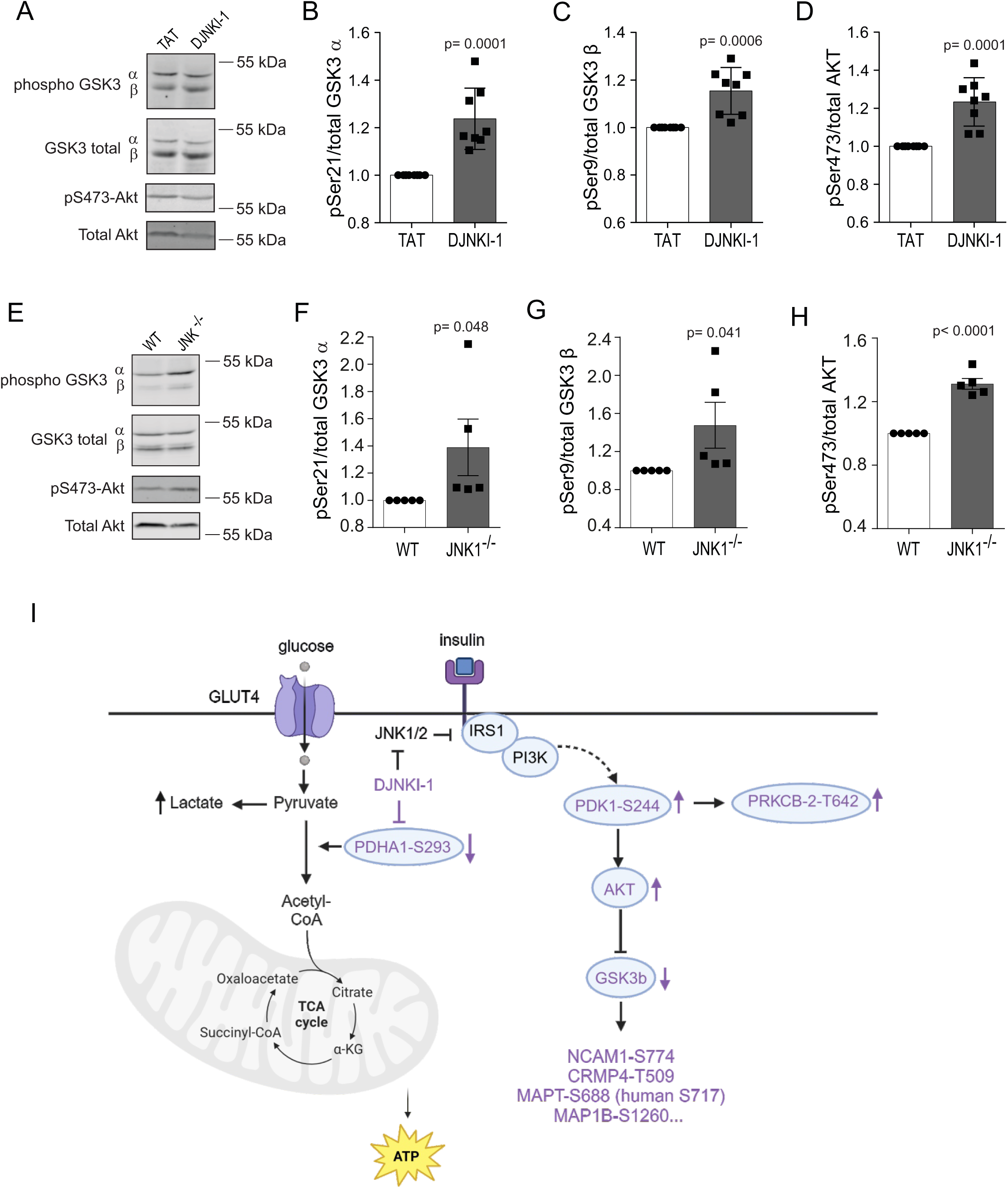
DJNKI-1 activates insulin pathway and neuronal Warburg effect signalling. A-C) Cortical neurons at 16 days *in vitro* were treated with DJNKI-1 (10 μM) or TAT carrier (10 µM) for 8 hours. Phosphorylation of GSK3-□/aon Ser9 or Ser21 is shown. Phosphorylation of GSK3a/b increases following DJNKI-1 indicating decreased activity. D) Quantitative data for phospho-Ser473-AKT is shown. E-H). Representative blots and quantitative analysis for GSK3a/b and AKT phospho-blotting in NAc of wild-type (WT) and *Jnk1-/-* mice. I-K) Phospho-Ser9 GSK3b and phospho-Ser473-AKT immunoblots and quantitative data are shown from hippocampus of WT and *Jnk1-/-* mice. Error bars represent standard errors of the mean. Significance was analysed by T-test. L) A scheme depicting the evidence from this study that DJNKI-1 regulates insulin signalling. In purple are shown the proteins where phosphosites with known function were regulated in DJNKI-1-treated mouse hippocampus. Arrows indicate the direction of activity regulation.

**Figure 6.**
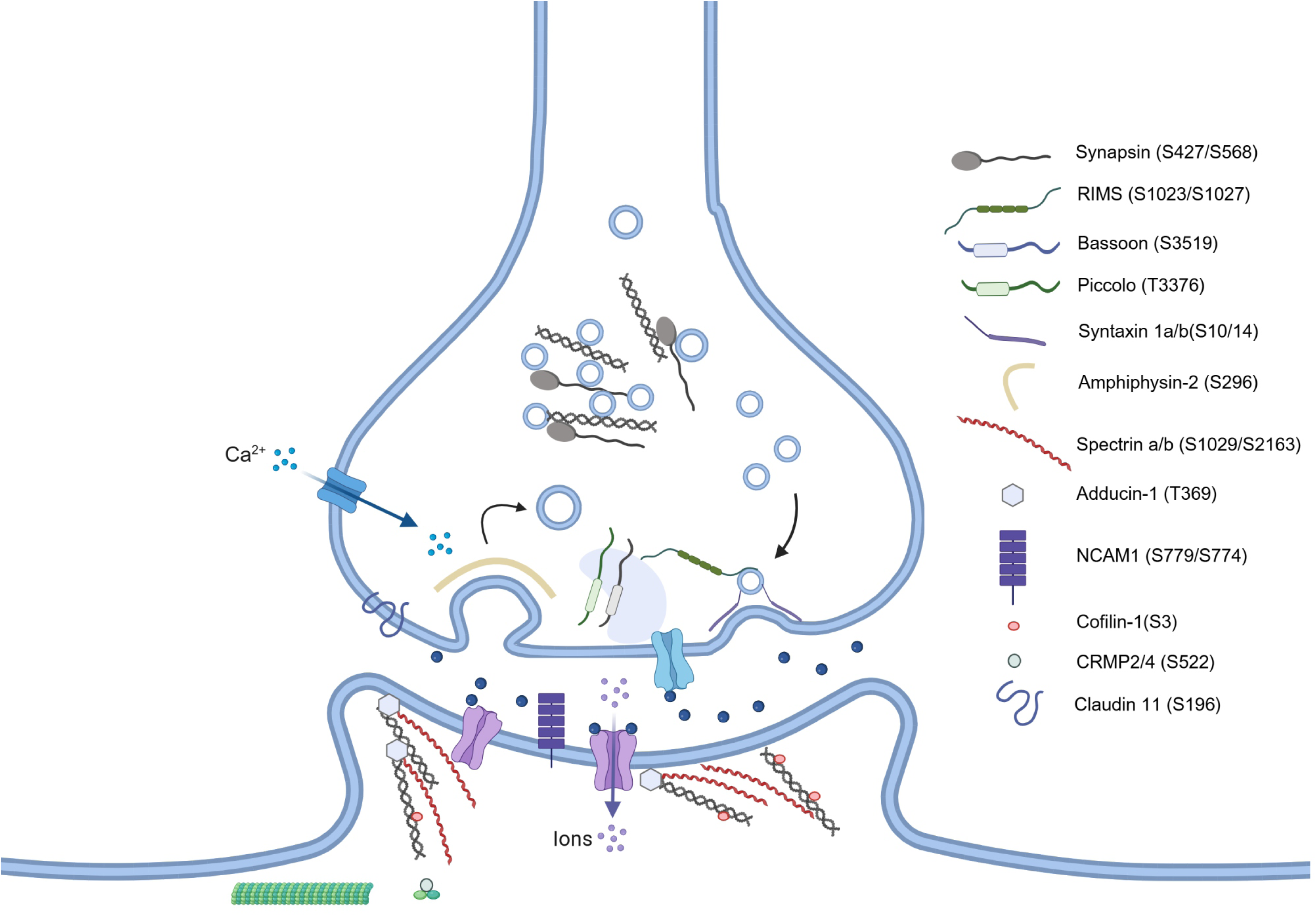
A scheme depicting the synaptic proteins whose phosphorylation is regulated by DJNKI-1 in this study. Quantitative data on phosphosite changes in hippocampus, nucleus accumbens and in responder and non-responder mouse hippocampus, can be found in Supplementary tables 1 and 2.

### DJNKI-1 treatment regulates insulin/glucose pathway regulation including pyruvate dehydrogenase inhibition

Figure 5L shows phosphosites and immunoblotting results for insulin/glucose pathway proteins regulated by DJNKI-1. The scheme includes IRS1 which JNK1 and 2 are known to phosphorylate and inhibit (Aguirre et al., 2002; Hirosumi et al., 2002). We show that in the hippocampus of DJNKI-1 treated mice, phosphorylation of PDPK1-S241 is augmented, signifying kinase activation (Levina et al., 2022). In line with this, DJNKI-1 induces phosphorylation-dependent activation of PRKCB isoform 2, a downstream target of PDPK1, also in hippocampus. Specifically, PRKCB-T641 phosphorylation increases indicating kinase activity increase, by analogy to PRKCB isoform 1 (Zhang et al., 1994). Finally, PDHA1 (pyruvate dehydrogenase E1 component subunit alpha) S293 phosphorylation increases, signifying enzymatic inhibition (Seifert et al., 2007). Inhibition of PDHA1 by DJNKI-1 is expected to promote lactate production.

## Discussion

Understanding the protein phosphorylation changes that contribute to neurological disorders is crucial for uncovering underlying mechanisms. Here we analyse phosphoproteomic changes in mice undergoing anxiety-inducing behavioural tests. We investigate how this phosphorylation is affected by chronic JNK inhibition, which reduces anxiety in animal models (Hollos et al., 2018; Li et al., 2024 2024; Mohammad et al., 2018; Morel et al., 2018; Stefanoska et al., 2018). Correlative analysis between phosphorylated protein intensities and anxiety features, alongside hub analysis, identifies phosphorylation changes that may contribute to the anxiolytic response. We report, large-fold phosphorylation changes upon treatment with DJNKI-1 on key microtubule stabilizing proteins, which are among the most abundant proteins expressed in brain. In neurons, microtubules are vital for axonal structure and transport in the nervous system, while also influencing synapse stability and synaptic vesicle priming (Chakraborti et al., 2016; Dent, 2020). In glial cells, microtubules have an additional role in driving the cytokine response (Adrian et al., 2023). These functions are under the control of microtubule associated proteins (MAPs) that bind and stabilize microtubules in response to phosphorylation control. We identify that multiple sites on MAP1A, MAP1B and MAP2 show decreased phosphorylation in hippocampus and nucleus accumbens, changing as much as -28-fold. One such site, MAP2-T1609 is a known JNK1 site that when phosphorylated increases microtubule binding and increases dendrite arborization in hippocampus CA3 pyramidal neurons (Björkblom et al., 2005; Komulainen et al., 2014). Also, S1352 and T1358 of MAP2 projection domain and MAP2-1783 (human S1782), are dephosphorylated in hippocampus and nucleus accumbens by DJNKI-1 treatment. MAP2-S1783 is similarly dephosphorylated in DJNKI-1 treated mice (Komulainen et al., 2014). Notably, phosphorylation of this site is elevated in schizophrenic brain, and is associated with regulation of basal dendrite complexity (Grubisha et al., 2022).

The first microtubule associated protein to be identified as a JNK1 target was MAP1B (Chang et al., 2003). Here, we validate this finding and identify that the JNK-phosphorylated site is MAP1B-S1260 (human T1265). This site is regulated in the hippocampus (-2-fold, p=0.008), and in the nucleus accumbens (-5-fold, p=0.003), in DJNKI-1 treated mice. Phosphorylation of this site controls axon branching and spacing (Scales et al., 2009; Ziak et al., 2024). Additional MAP1B phosphorylation sites (S11373, S1151, S1384, S1307, S1371, S1373 and S1781) also decrease in the nucleus accumbens upon DJNKI-1 treatment, with S1151 showing a -25-fold change. In addition to MAP1b, MAP1a sites are also regulated in hippocampus and nucleus accumbens. These results support the known role of JNK in controlling functions that are dependent on microtubule stability such as dendrite and axon stability and neuronal migration (Björkblom et al., 2005; Chang et al., 2003; Komulainen et al., 2014; Westerlund et al., 2011). Notably, MAP1B and MAP2 show functional redundancy for controlling dendrite arborization and anterior commissure and corpus callosum stability (Teng et al., 2001). Notably, JNK1 has also been shown to regulate the stability of these commissures and dendrite architecture (Björkblom et al., 2005; Chang et al., 2003). It is worth mentioning that MAP1A and B may also act at synapses, as the *Drosophila* MAP1 homologue Futsch regulates active zone density and neurotransmitter release (Lepicard et al., 2014). The data described here extends these earlier studies by identifying JNK1 specific phosphorylation sites on microtubule associated proteins. Moreover, our results indicate that JNK regulates these sites in adult brain.

Synaptic protein phosphorylation is highly regulated by DJNKI-1 in this study. Accordingly, synaptic proteins that interact with actin, and those controlling endocytosis undergo phosphorylation change. These include amphiphysin 2 (Bin1), adducin 1, piccolo, bassoon, CARMIL2 (Rltpr), AP180 (Snap91), cofilin-1 (dystrophin) and spectrin 1. Adducin 1 for example is dephosphorylated on several sites in DJNKI-1-treated hippocampus. Particularly in responder mice that exhibit low anxiety-like behavior following DJNKI-1, adducin 1-T639 and -T610 decrease by -18 and -22-fold. Adducin 1 confers stability to synapses, accordingly, knockout of adducin 1 increases turnover of filopodia (Bednarek and Caroni, 2011; Pielage et al., 2011). JNK inhibition, in turn has been shown to increase filopodia preponderance, and in *Jnk1-/-* mouse hippocampus, the number of thin spines increases (Komulainen et al., 2020). Thus adducin 1 is interesting as a potential mediator of these JNK-dependent effects on spine morphology.

Importantly, among the most significant changes in DJNKI-1 treated mouse hippocampus is cofilin-1 (DSTN/dystrophin) phosphorylation on S3, this is decreased by 5-fold. Phosphorylation on this site stabilizes actin filament growth and promotes formation of mature spines (Ben Zablah et al., 2020; Mizuno, 2013), thus, cofilin S3 phosphorylation, like adducin-1, may mediate the effects of JNK on spine morphology and AMPA receptor trafficking (Gu et al., 2010; Hollos et al., 2020; Komulainen et al., 2020; Thomas et al., 2008). Cofilin is also expressed pre-synaptically where it may regulate neurotransmitter release (Ben Zablah et al., 2020). Another phosphoprotein that is highly regulated by DJNKI-1 is claudin-11 S196. This protein is a tight junction protein component of myelin. Mice lacking the myelin Cldn-11 gene display a low anxiety phenotype (Maheras et al., 2018). While the phosphorylation of S196 is not functionally studied as of yet, it resides in the C-terminal adjacent to sites that regulate migration.

SNARE proteins syntaxin 1a and b are also regulated by DJNKI-1 in this study. These proteins mediate the fusion of synaptic vesicles with the plasma membrane, thereby facilitating exocytosis (Jahn et al., 2024). We find that in the hippocampus, DJNKI-1 decreases phosphorylation of syntaxin 1a at T10, and syntaxin 1b at S14. These N-terminal sites on syntaxins reside within the MUNC18 binding domain, which is required for fusion (Rickman and Duncan, 2010; Yang et al., 2023). Phospho-mimetic syntaxin 1 S14 (conserved in STX1a and b), prevents interaction with MUNC18 and impairs exocytosis (Rickman and Duncan, 2010; Shi et al., 2021). Consistent with our finding that DJNKI-1 alters syntaxin 1 phosphorylation, JNKs (2 and 3) have been shown to co-purify with syntaxins from nerve terminal preparations, while JNK2 is required for spontaneous neurotransmitter release (Abrahamsson et al., 2017; Biggi et al., 2017). Syntaxin 1 S14 phosphorylation has been shown to be required for non-vesicular dopamine release through reversal of the dopamine transporter (Shekar et al., 2023). By extrapolation from these studies, the decreased N-terminal phosphorylation of syntaxins 1a and b identified in DJNKI-1-treated mice could conceivably inhibit non-evoked transmitter release. Interestingly, in DJNKI-1-treated cortical slices, the frequency of spontaneous mEPSCs decreases (Biggi et al., 2017), demonstrating that JNK activity in the presynaptic zone can affect neurotransmitter release. Also, we see increased pyruvate dehydrogenase S293 phosphorylation, signifying reduced neural activity (Yang et al., 2024).

One of the aims of this study was to analyse the differential effects of DJNKI-1 in responder and non-responder mice. Interestingly, we only detect one phosphosite that is oppositely regulated in both groups. This is NCAM1-S774, a site which is required for NCAM1 interaction with CRMP2 (DPYSL2) (Karbe et al., 2013), whose phosphorylation is also regulated by DJNKI-1. Both proteins share common functions in the regulation of synaptic pruning and synapse stability (Duncan et al., 2021; Ziak et al., 2020), and may mediate the neuroplasticity effects of JNK. Interestingly, we find that CRMP2 S546 phosphorylation increases 7-9-fold in responder and non-mice, however S522, a functionally characterized site on CRMP2, increases 3.6-fold upon DJNKI-1 treatment, only in non-responder mice. Phosphorylation on this site stabilizes sodium channel NaV1.7 and calcium channel CaV2.2 (N-type voltage gated calcium channel) stability at the cell surface, thereby increasing presynaptic (Stratton et al., 2020). Thus, would could conceivably predict increased presynaptic firing in the non-responder mice.

We detect DJNKI-1dependent activation of the insulin signalling pathway in our experiments including downstream regulation of pyruvate metabolism. This supports the already characterised role of JNKs 1 and 2 in the negative regulation of IRS1 (Hirosumi et al., 2002; Solinas and Becattini, 2017). New in this study, we identify activation of PDPK1 and its downstream targets AKT and PRKCB upon JNK inhibition. Moreover, DJNKI-1 treatment increases the phosphorylation of PDHA1 (pyruvate dehydrogenase) on S293 which is known to inhibit its enzymatic function (Seifert et al., 2007), and this is specifically found in the 50% of mice that do not display a so-called anxiolytic response in the light/dark test. This adds to earlier findings which showed that JNK1 promotes glycolysis by activating phosphofructokinase-1 (Deng et al., 2008). Downstream on this pathway, inhibition of PDHA1 through S293 phosphorylation is expected to prevent pyruvate from entering the TCA cycle promoting lactate production rather than oxidative phosphorylation (Seifert et al., 2007). This is known as the neuronal Warburg effect (Bouzier-Sore et al., 2006; van Hall et al., 2009; Wyss et al., 2011). It may benefit neurons by avoiding production of excessive reactive oxygen species which would be generated should pyruvate be utilized for oxidative phosphorylation. Interestingly, lactate is reported to stimulate neurogenesis which has an antidepressant effect (Bas-Orth et al., 2017; Benarroch, 2024), and we have previously reported that JNK1 inhibition (with DJNKI-1) or *Jnk1* knockout, increases ventral hippocampal neurogenesis (Mohammad et al., 2018). Interestingly, glycolysis inhibition has also been shown to improve anxiety and depressive-like behaviours (Khatibi et al., 2023; Liu et al., 2024).

Consistent with these findings on glucose metabolic pathway regulation, we find that GSK3b activity is repressed in DJNKI-1 treated mice. Moreover, 28 putative GSK3b motifs show downregulated phosphorylation at both the priming site and the GSK3b site. These motifs are also found on three previously reported GSK3b target sites; Dpysl3-T514 (Cole et al., 2004), MAP1B-S1260 (Trivedi et al., 2005) and MAPT-S692 (Sayas and Ávila, 2021). Additionally, SPTBN1 (spectrin beta) is linked to GSK3 (Shinde et al., 2017), and other interesting phosphoproteins regulated by DJNKI-1 on GSK3 motifs are DSTN (dystrophin/cofilin) S3, MAP2 T1620 and T1623, and MAPT S692 (human S717), a site that is implicated in neurofibrillary tangles (Cruz et al., 2003).

The negative regulation of GSK3b by DJNKI-1 aligns with the known low depressive behaviours observed in *GSK3b-/-* mice (Ziak et al., 2024). Similarly, silencing of (Mohammad et al., 2018; Ziak et al., 2024)GSK3b in the nucleus accumbens was shown to lower anxiety in rats (Crofton et al., 2017). GSK3b inhibition by lithium, may contribute to its mood-stabilizing effects (Gould et al., 2004; Li and Jope, 2010; Stambolic et al., 1996), and to the action of anti-depressant drugs (Beaulieu, 2012).

We would like to point out aspects of this study that should be taken into account while interpreting the data. Firstly, the DJNKI-1 treatment was for six weeks, the time required to manifest an anxiolytic response (Mohammad et al., 2018). This period provides ample time for indirect and neuroplasticity changes to affect the protein phosphorylation profiles. Secondly, the responder, non-responder split analysis is done based on the light dark test behavior and therefore cannot be interpreted in the context of EPM behaviours. Finally, the correlations of phosphorylation and behaviours, do not taken into account signalling and circuit changes in other brain regions that may be important. Also, we are drawing parallels between phosphosite functions and possible contributions to anxiety behaviors, however these relationships have not been directly tested.

In summary, this study demonstrates that JNK inhibitor treatment regulates a considerable number of sites with known function. These findings uncover synaptic targets involved in neurotransmission regulation and insulin pathway enzymatic control, which appears to shift towards lactate production, suggesting a neuronal Warburg effect. Notably, many of the negatively regulated phosphosites are within a GSK3 motif, and consistent with this, we show that DJNKI-1 inhibits GSK3b activity. This data provides a valuable addition to the existing knowledge of JNK-regulated phosphosites, providing a systematic overview of JNK regulated phosphorylation in hippocampus and nucleus accumbens.

## Acknowledgements

This project was funded by grants from the Marie Slodowska Curie Actions r’BIRTH Initial Training Network grant #608346 and the Research Council of Finland grant #310583 to E.C..

