## Supplementary table 1 for "Identifying JNK-regulated phosphoproteome markers of anxiety-like behaviour in mouse hippocampus"

**Hippocampus**

| Protein | Protein.nai | Gene.nam | Amino.acic | Position | Sequence. | Fold Chan | ROTS_Pvalue |
|---|---|---|---|---|---|---|---|
| Q3UHJ0 | AP2-assoc | Aak1 | 604 | T | GQIQAPV | 1.94 | 0.049848 |
| E9Q1K3 | | Add1 | 629 | S | GSEENLD | -2.08 | 0.001508 |
| Q9QYC0-2 | Alpha-add | Add1 | 600 | S | GSEENLD | -2.08 | 0.001508 |
| Q9QYC0 | Alpha-add | Add1 | 600 | S | GSEENLD | -2.08 | 0.001508 |
| E9Q1K3;Q | Alpha-add | Add1 | 355 | S | NLVLLDP | -1.97 | 0.006469 |
| E9Q1K3;Q | Alpha-add | Add1 | 358 | T | LLDPGKY | -1.97 | 0.006469 |
| Q9QYC0 | Alpha-add | Add1 | 610 | T | EQKEKSP | -5.52 | 0.020551 |
| E9Q1K3 | | Add1 | 639 | T | EQKEKSP | -5.49 | 0.021113 |
| Q9QYC0-2 | Alpha-add | Add1 | 610 | T | EQKEKSP | -5.49 | 0.034749 |
| E9Q1K3;Q | Alpha-add | Add1 | 353 | S | PDNLVLL | 1.76 | 0.035431 |
| Q9QYB8 | Beta-addu | Add2 | 692 | S | DSYKDKT | -2.84 | 0.001049 |
| Q9QYB5 | Gamma-ad | Add3 | 681 | S | ERTEEVL | -1.83 | 0.000924 |
| Q9QYB5 | Gamma-ad | Add3 | 673 | S | IEITIKSPE | 1.68 | 0.004891 |
| Q9QYB5 | Gamma-ad | Add3 | 11 | T | _____MSS | 2.93 | 0.010644 |
| Q3UNS7 | | Adgrg4 | 15 | S | _MNLSMF | 20.95 | 0.038011 |
| Q9WTQ5 | A-kinase a | Akap12 | 584 | S | PSEAPQE | -2.19 | 0.025212 |
| Q8C8R3 | Ankyrin-2 | Ank2 | 3362 | S | ASSLDAK | 2.39 | 0.034375 |
| Q8C8R3 | Ankyrin-2 | Ank2 | 2364 | S | KTAEGTE | -1.6 | 0.038034 |
| Q8C8R3;Q | Ankyrin-2 | Ank2 | 3692 | S | DCLPKTE | -1.52 | 0.042098 |
| Q9EPJ9 | ADP-ribos | Arfgap1 | 135 | T | EGKEWSL | 2.01 | 0.003818 |
| A2A5R2 | Brefeldin A | Arfgef2 | 218 | S | QEARELE | -2.55 | 0.00859 |
| A2A5R2 | Brefeldin A | Arfgef2 | 227 | S | MQSKPQS | -2.58 | 0.009266 |
| Q5FWK3 | Rho GTPa | Arhgap1 | 47 | S | NWPSDEM | -2.71 | 0.000172 |
| Q5FWK3 | Rho GTPa | Arhgap1 | 51 | S | DEMPDFP | -2.84 | 0.000537 |
| Q811P8 | Rho GTPa | Arhgap32 | 952 | S | TSWDKSV | 1.59 | 0.0078 |
| P70704 | Phospholip | Atp8a1 | 29 | S | RAEGYEK | -3.93 | 0.002257 |
| P70704 | Phospholip | Atp8a1 | 25 | S | EIRSRAE | -4.09 | 0.002606 |
| O08539 | Myc box-d | Bin1 | 296 | S | VKAQPSD | -1.82 | 0.002898 |
| O08539 | Myc box-d | Bin1 | 296 | S | VKAQPSD | -2.32 | 0.042243 |
| O08539 | Myc box-d | Bin1 | 298 | S | AQPSDNA | -2.19 | 0.049339 |
| O88737 | Protein ba | Bsn | 3519 | S | RPPMRSQ | -1.82 | 0.000216 |
| O88737 | Protein ba | Bsn | 2595 | T | PDPSTVR | -2.1 | 0.003003 |
| Q80X80 | C2 domain | C2cd2l | 464 | S | VTTVQSR | -1.81 | 0.004514 |
| Q80X80 | C2 domain | C2cd2l | 376 | S | DAELLGQ | 4.04 | 0.006174 |
| A0A286YC | Calcium/ca | Camk2a | 234 | S | HKLYQQIK | 2.17 | 0.002239 |
| P11798 | Calcium/ca | Camk2a | 337 | T | KKNDGVK | -2.15 | 0.033502 |
| Q9CYT6 | Adenylyl cy | Cap2 | 308 | S | RAQGQIR | -1.76 | 0.008298 |
| Q9CYT6 | Adenylyl cy | Cap2 | 311 | S | GQIRSPTI | -1.76 | 0.008298 |
| Q9JKC6 | Cell cycle e | Cend1 | 94 | T | KPAPTVPA | -5.12 | 0.003927 |
| Q9JKC6 | Cell cycle e | Cend1 | 89 | S | NHSNLKP | 32.87 | 0.032768 |
| P18760 | Cofilin-1 | Cfl1 | 3 | S | _________ | -6.04 | 0.007371 |
| Q60771 | Claudin-11 | Cldn11 | 196 | S | DAQSFGE | -8.72 | 7.24E-05 |
| P97427 | Dihydropyr | Crmp1 | 509 | T | VSRGMYD | 1.73 | 0.027054 |
| Q61301;P2 | Catenin alp | Ctnna2 | 654 | S | DSDFEQE | -2.13 | 0.033674 |
| Q61301;P2 | Catenin alp | Ctnna2 | 657 | T | FEQEDYD | -1.78 | 0.038468 |
| Q02248 | Catenin be | Ctnnb1 | 552 | S | VQLLVRA | 1.94 | 0.018722 |
| O35927 | Catenin de | Ctnnd2 | 412 | S | SSQHGHL | 2.12 | 0.004208 |
| O35927 | Catenin de | Ctnnd2 | 264 | S | LYYSSSTL | -2.16 | 0.018973 |
| A0A0R4J1 | Drebrin | Dbn1 | 142 | S | GAIGQRL | 1.6 | 0.023288 |
| Q9D415 | Disks large | Dlgap1 | 437 | S | INRSLDSL | 2.32 | 0.016796 |
| B1AZP2 | Disks large | Dlgap4 | 405 | S | KTAARRQ | -1.56 | 0.011007 |
| P60904 | DnaJ hom | Dnajc5 | 10 | S | ______MA | 5.93 | 0.015816 |
| Q80TZ3 | Putative ty | Dnajc6 | 35 | S | AAGENRM | 3.54 | 0.010495 |
| Q80TZ3 | Putative ty | Dnajc6 | 36 | S | AGENRMK | 3.32 | 0.018733 |
| O08553 | Dihydropyr | Dpysl2 | 542 | S | QAPPVRN | 8.09 | 0.000623 |
| O08553 | Dihydropyr | Dpysl2 | 522 | S | SVTPKTV | 3.18 | 0.049748 |
| O08553 | Dihydropyr | Dpysl2 | 521 | T | VSVTPKT | 3.18 | 0.049748 |

| | | | | | | | |
|---|---|---|---|---|---|---|---|
| Q62188 | Dihydropyr | Dpysl3 | 509 | T | VPRGMYD | 2.11 | 0.003235 |
| Q62188 | Dihydropyr | Dpysl3 | 522 | S | TTTPKGG | -2.48 | 0.017931 |
| Q62188 | Dihydropyr | Dpysl3 | 508 | T | AVPRGMY | -1.82 | 0.019553 |
| Q62188 | Dihydropyr | Dpysl3 | 509 | T | VPRGMYD | -1.87 | 0.020894 |
| Q62188 | Dihydropyr | Dpysl3 | 514 | T | YDGPVFD | -1.86 | 0.049743 |
| Q9R0P5 | Destrin (C | Dstn | 3 | S | ________ | -5.15 | 5.90E-05 |
| Q8R1Q8 | Cytoplasm | Dync1li1 | 510 | S | VHAELDR | -1.91 | 0.001689 |
| Q8R1Q8 | Cytoplasm | Dync1li1 | 207 | S | RDFQEYV | 2.1 | 0.045166 |
| P57776 | Elongation | Eef1d | 133 | S | SSPTPRA | 1.72 | 0.000564 |
| Q8JZQ9 | Eukaryotic | Eif3b | 75 | S | VRAKPAA | -1.74 | 0.001782 |
| Q8JZQ9 | Eukaryotic | Eif3b | 79 | S | PAAQSEE | -2.24 | 0.04556 |
| A0A286YD | Band 4.1-li | Epb4.1l3 | 505 | S | ERTDTAA | -2.93 | 0.003176 |
| P70429 | Ena/VASP | Evl | 329 | S | PNSSEAG | 1.79 | 0.042568 |
| Q6NS60 | F-box only | Fbxo41 | 477 | T | RQAIQNW | 2.16 | 0.048587 |
| P09528 | Ferritin hea | Fth1 | 5 | S | ________ | 4.08 | 0.016994 |
| P07901 | Heat shock | Hsp90aa1 | 263 | S | EEKESDD | -1.71 | 0.028348 |
| P11499 | Heat shock | Hsp90ab1 | 255 | S | EDKEDEE | -1.77 | 0.011946 |
| Q61699 | Heat shock | Hsph1 | 810 | S | NVCEPVV | 1.87 | 0.029939 |
| Q5DU25 | IQ motif ar | Iqsec2 | 383 | S | RMSRRIIL | 1.64 | 0.000432 |
| Q91YS4 | | Klc2 | 151 | S | HLLFMSQ | -2.07 | 0.018056 |
| Q9QYR6 | Microtubule | Map1a | 1789 | S | PEMTGQP | -2.42 | 0.034541 |
| Q9QYR6 | Microtubule | Map1a | 1575 | T | KSSFLEDI | 4.9 | 0.038819 |
| P14873 | Microtubule | Map1b | 1260 | S | RLSPAKSI | -2.03 | 0.008747 |
| P20357 | Microtubule | Map2 | 1609 | T | TSTPTTP | -2.94 | 0.004763 |
| P20357 | Microtubule | Map2 | 1352 | S | EIEMAAE | -2.39 | 0.006332 |
| P20357 | Microtubule | Map2 | 1358 | T | EAQAEPK | -2.39 | 0.006332 |
| P20357 | Microtubule | Map2 | 1783 | S | HAKARVD | -1.76 | 0.021943 |
| P20357 | Microtubule | Map2 | 938 | S | AAGRVKD | -1.91 | 0.02232 |
| A2AJI0 | MAP7 dom | Map7d1 | 115 | S | PTATGPR | -1.79 | 0.003386 |
| P10637 | Microtubule | Mapt | 692 | S | KTDHGAE | -2.45 | 0.006651 |
| P10637 | Microtubule | Mapt | 695 | T | HGAEIVYK | 79.04 | 0.023792 |
| P04370-4 | Myelin bas | Mbp | 139 | S | PPSQGKG | 2.74 | 0.018247 |
| P04370-6; | Myelin bas | Mbp | 122 | T | ENPVVHF | 2.57 | 0.023476 |
| P14152 | Malate deh | Mdh1 | 241 | S | TVQQRGA | 1.67 | 0.032224 |
| Q9D2P8 | Myelin-ass | Mobp | 85 | S | RRATSPQ | 2.03 | 0.01221 |
| Q9D1H8 | 39S riboso | Mrpl53 | 38 | S | FEKNVES | 4.78 | 0.03674 |
| A0A494BB10 | | NA | 42 | S | LSSKIKEQ | 2.1 | 0.04824 |
| Q9QYG0 | Protein ND | Ndrg2 | 332 | S | MASSCMT | -1.77 | 0.017714 |
| Q9QYG0 | Protein ND | Ndrg2 | 338 | S | TRLSRSR | -1.77 | 0.017714 |
| P19246 | Neurofilam | Nefh | 834 | S | EGAKPPE | -2.63 | 0.011712 |
| P19246 | Neurofilam | Nefh | 839 | T | PEKAKPLI | -2.64 | 0.0164 |
| P19246 | Neurofilam | Nefh | 739 | S | AVKSPAE | -1.73 | 0.021587 |
| P19246 | Neurofilam | Nefh | 607 | S | EAKSPSE | -1.64 | 0.036205 |
| P19246 | Neurofilam | Nefh | 673 | S | EAKSPAE | 1.65 | 0.044123 |
| P19246 | Neurofilam | Nefh | 757 | S | EAKSPGE | -1.57 | 0.046389 |
| P19246 | Neurofilam | Nefh | 763 | S | EAKSPAE | -1.57 | 0.046389 |
| P08553 | Neurofilam | Nefm | 769 | S | GREEEKG | 1.61 | 0.038661 |
| Q9Z0P4 | Paralemmi | Palm | 124 | S | KENSAAP | -2.74 | 0.016647 |
| Q9QYX7 | Protein pic | Pclo | 3376 | T | DDPRNLK | -3.06 | 0.001433 |
| Q9QYX7 | Protein pic | Pclo | 1332 | S | IPSDEKDL | -1.57 | 0.008941 |
| Q9QYX7 | Protein pic | Pclo | 3610 | S | PPASPKT | -1.78 | 0.025435 |
| Q9QYX7 | Protein pic | Pclo | 3616 | S | TAKMMQP | -1.78 | 0.025435 |
| P35486 | Pyruvate d | Pdha1 | 293 | S | GPILMELQ | 4.58 | 0.017257 |
| Q9Z2A0 | 3-phospho | Pdpk1 | 244 | S | TAKVLSPE | 2.92 | 0.002855 |
| Q9Z1B3 | 1-phospha | Plcb1 | 978 | S | RAALEKS | 13.06 | 0.015848 |
| Q3UM45 | Protein pho | Ppp1r7 | 24 | S | QQQSQEM | -2.33 | 0.003249 |
| Q3UM45 | Protein pho | Ppp1r7 | 27 | S | SQEMMEV | -2.33 | 0.003249 |
| P68404-2 | Protein kin | Prkcb | 641 | T | AENFDRF | 1.77 | 0.001468 |
| P16054 | Protein kin | Prkce | 337 | S | KLAAGAE | -1.8 | 0.020873 |

| | | | | | | | |
|---|---|---|---|---|---|---|---|
| E9PUL5 | Proline-ricl | Prrt2 | 254 | S | GGHPGSI | 6.87 | 0.012723 |
| E9PUC5 | PH and SE | Psd3 | 339 | S | LTTDGNE | 2.83 | 0.023649 |
| Q99JF8 | PC4 and S | Psip1 | 171 | S | QVDTEEA | 2.36 | 0.032149 |
| Q3TXS7 | 26S protea | Psmd1 | 315 | S | EEKTASA | -1.57 | 0.023248 |
| Q3TXS7 | 26S protea | Psmd1 | 311 | T | PMETEEK | -1.57 | 0.023248 |
| Q9QVP9 | Protein-tyr | Ptk2b | 375 | S | GSLIMHAI | 4.2 | 0.040144 |
| Q99NE5 | Regulating | Rims1 | 1023 | S | SAPPSPLI | -2.37 | 0.004006 |
| Q99NE5 | Regulating | Rims1 | 1027 | S | SPLLTRTI | -2.55 | 0.006661 |
| Q3V3V9 | Leucine-ric | Rltpr | 1134 | S | SKRKQSK | -2.29 | 0.000565 |
| Q3UJU9 | Regulator | Rmdn3 | 46 | S | YSQRWKI | 1.88 | 0.02431 |
| P47708 | Rabphilin-3 | Rph3a | 679 | S | KKIERWH | -3.17 | 0.005735 |
| P47708 | Rabphilin-3 | Rph3a | 680 | S | KIERWHQ | -2.29 | 0.024405 |
| D3YXK2 | Scaffold at | Safb | 372 | S | LVRAPTA | 1.55 | 0.006004 |
| Q6ZPE2 | Myotubular | Sbf1 | 706 | S | QTHIRALY | 2.72 | 0.049996 |
| Q8BP27 | Swi5-depe | Sfr1 | 67 | S | PPTSPAVI | -1.72 | 0.002308 |
| Q8VD37 | SH3-conta | Sgip1 | 335 | T | PEHVTPE | 2.86 | 0.027338 |
| Q61548 | Clathrin co | Snap91 | 296 | S | HLNTLEG | -1.58 | 0.007987 |
| Q61548 | Clathrin co | Snap91 | 300 | S | LEGKKPG | -1.58 | 0.007987 |
| P16546 | Spectrin al | Sptan1 | 1029 | S | GFVPAAY | 6.27 | 0.012532 |
| Q62261 | Spectrin be | Sptbn1 | 2163 | S | EMVNGA | -2.69 | 0.009282 |
| Q62261 | Spectrin be | Sptbn1 | 2164 | S | MVNGAAE | -2.84 | 0.009306 |
| Q62261 | Spectrin be | Sptbn1 | 2168 | S | AAEQRTS | -2.76 | 0.010163 |
| Q62261 | Spectrin be | Sptbn1 | 2102 | S | VRRQQEE | 1.68 | 0.046578 |
| P54227 | Stathmin | Stmn1 | 38 | S | ILSPRSKE | 2.09 | 0.010422 |
| O88935 | Synapsin-1 | Syn1 | 427 | S | NKMTQAL | 2.75 | 0.032787 |
| Q64332 | Synapsin-2 | Syn2 | 426 | S | VISKMNQI | 1.71 | 0.041923 |
| Q9CPQ3 | Mitochondi | Tomm22 | 15 | S | _MAAAVA | 3.37 | 0.00279 |
| Q7TQD2 | Tubulin po | Tppp | 15 | T | _MADSKA | 2.36 | 0.007562 |
| Q9DBS2 | Tumor pro | Tprg1l | 34 | T | LAAGEDA | 2.89 | 0.000877 |
| Q02053 | Ubiquitin-li | Uba1 | 835 | S | SVDDSRL | 1.7 | 0.03292 |
| Q6ZPJ3 | E2/E3 hybi | Ube2o | 836 | S | LKNMTVE | 2.23 | 0.045379 |
| Q60932 | Voltage-de | Vdac1 | 117 | S | DQLARGL | 1.52 | 0.035925 |
| Q8C0E2 | Vacuolar p | Vps26b | 302 | S | QQEVVLW | 1.74 | 0.002076 |

**Nucleus accumbens**

| Protein | Protein.nai | Gene.nam | Amino.acic | Position | Sequence. | Fold chang | ROTS_Pvalue |
|---|---|---|---|---|---|---|---|
| P28661 | Septin-4 | Sept4 | 68 | S | KTRVARP | 2.43 | 0.045333 |
| Q8CHH9 | Septin-8 | Sept8 | 10 | S | _______MA | 3.21 | 0.025793 |
| Q8BL65 | Actin-bindi | Ablim2 | 475 | T | KKTTWLL | -5.35 | 0.017207 |
| Q8BL65 | Actin-bindi | Ablim2 | 294 | S | SSESIVSV | -23.18 | 0.019903 |
| E9Q1K3;Q | Alpha-add | Add1 | 11 | T | _____MNC | 7.47 | 0.006502 |
| E9Q1K3;Q | Alpha-add | Add1 | 353 | S | PDNLVLLI | 5.11 | 0.008156 |
| E9Q1K3;Q | Alpha-add | Add1 | 353 | S | 2PDNLVLL | -4.34 | 0.011707 |
| E9Q1K3 | | Add1 | 639 | T | EQKEKSP | 1.94 | 0.039284 |
| Q9QYC0-2 | Alpha-add | Add1 | 610 | T | EQKEKSP | 1.94 | 0.039284 |
| Q9QYC0 | Alpha-add | Add1 | 610 | T | EQKEKSP | 1.94 | 0.039284 |
| Q9QYB8 | Beta-addu | Add2 | 606 | S | 2KPGSPV | -5.62 | 0.008524 |
| Q9QYB8 | Beta-addu | Add2 | 602 | S | 2IATEKPG | -5.65 | 0.008666 |
| Q9QYB5 | Gamma-ac | Add3 | 677 | S | 2IKSPERT | -2.04 | 0.045532 |
| Q9QYB5 | Gamma-ac | Add3 | 681 | S | 2ERTEEVI | -2.04 | 0.045532 |
| O54774 | AP-3 comp | Ap3d1 | 784 | S | RVDIITEE | -2.25 | 0.04192 |
| B2RUJ5 | Amyloid be | Apba1 | 267 | S | SPEKEAE | -2.58 | 0.018954 |
| Q9EPJ9 | ADP-ribos | Arfgap1 | 342 | S | SFWETFG | 3.02 | 0.039508 |
| Q91YM2 | Rho GTPa | Arhgap35 | 1179 | S | RLGRFAS | -3.22 | 0.009911 |
| Q8VDN2 | Sodium/po | Atp1a1 | 228 | S | VDNSSLT | -4.04 | 0.009158 |
| P70704 | Phospholip | Atp8a1 | 29 | S | 2RAEGYE | -2.44 | 0.01752 |
| P70704 | Phospholip | Atp8a1 | 25 | S | 2EIRSRAE | -2.23 | 0.026747 |
| Q91XV3 | Brain acid | Basp1 | 31 | T | DEKAKDK | 56.28 | 0.028005 |
| O08539 | Myc box-d | Bin1 | 298 | S | AQPSDNA | -16.75 | 0.001086 |
| O08539 | Myc box-d | Bin1 | 298 | S | 2AQPSDN | -3.74 | 0.017744 |
| O08539 | Myc box-d | Bin1 | 296 | S | 2VKAQPS | -3.79 | 0.020657 |
| O88737 | Protein ba | Bsn | 3512 | S | SMAHGRA | -5.34 | 0.006581 |
| O88737 | Protein ba | Bsn | 3519 | S | 2RPPMRS | -4.26 | 0.008924 |
| O88737 | Protein ba | Bsn | 3512 | S | 2SMAHGF | -4.6 | 0.012685 |
| O88737 | Protein ba | Bsn | 2860 | S | 2QQTLPR | -2.65 | 0.036327 |
| O88737 | Protein ba | Bsn | 2866 | S | 2PMKTLQ | -2.65 | 0.036327 |
| O70589-3 | | Cask | 571 | S | SITFKIVPS | -2.48 | 0.035047 |
| O70589-3 | | Cask | 570 | S | GSITFKIVI | 7.42 | 0.043126 |
| Q61301;P2 | Catenin al | Ctnna2 | 654 | S | 2DSDFEQ | -2.99 | 0.006529 |
| Q61301;P2 | Catenin al | Ctnna2 | 657 | T | 2FEQEDY | -2.81 | 0.049056 |
| O35927 | Catenin de | Ctnnd2 | 273 | S | APPRGGS | 3.49 | 0.019121 |
| Q62188 | Dihydropyr | Dpysl3 | 509 | T | 2VPRGMY | -3.25 | 0.00716 |
| Q62188 | Dihydropyr | Dpysl3 | 522 | S | 2TTTPKG | -5.85 | 0.046359 |
| Q8R1Q8 | Cytoplasm | Dync1li1 | 510 | S | 2VHAELDI | -2.84 | 0.029988 |
| Q8JZQ9 | Eukaryotic | Eif3b | 75 | S | 2VRAKPA | -16.55 | 0.00092 |
| Q8JZQ9 | Eukaryotic | Eif3b | 79 | S | 2PAAQSE | -13.31 | 0.033139 |
| P09528 | Ferritin hea | Fth1 | 5 | S | __________ | 29.32 | 0.029568 |
| D3YXG0 | | Hmcn1 | 4087 | Y | 2LNVQVP | -6.06 | 0.025332 |
| Q8VEK3 | Heterogen | Hnrnpu | 4 | S | __________ | -3.87 | 0.013193 |
| P11499 | Heat shock | Hsp90ab1 | 255 | S | EDKEDEE | -1.99 | 0.024696 |
| Q8BNW9 | Kelch repe | Kbtbd11 | 316 | S | 2CLLAAAL | -6.35 | 0.035255 |
| P21619 | Lamin-B2 | Lmnb2 | 427 | S | GRGKRRF | 4.01 | 0.015325 |
| Q9QYR6 | Microtubul | Map1a | 981 | S | HPGEPAL | -4.32 | 0.009648 |
| Q9QYR6 | Microtubul | Map1a | 1606 | S | DGAVPEK | -4.49 | 0.011387 |
| Q9QYR6 | Microtubul | Map1a | 1772 | S | 2DKLTRSI | -6.29 | 0.031316 |
| P14873 | Microtubul | Map1b | 1373 | S | 2FEFSEAK | -26.1 | 0.000865 |
| P14873 | Microtubul | Map1b | 1151 | S | PRDVMSD | -28.48 | 0.001946 |
| P14873 | Microtubul | Map1b | 1384 | S | 2RASLSPI | -14.08 | 0.00226 |
| P14873 | Microtubul | Map1b | 1260 | S | 2RLSPAKS | -5.31 | 0.003468 |
| P14873 | Microtubul | Map1b | 1384 | S | RASLSPM | -10.13 | 0.00424 |
| P14873 | Microtubul | Map1b | 1371 | S | VSFEFSEA | -4.17 | 0.007015 |
| P14873 | Microtubul | Map1b | 1307 | S | VSPGVTQ | -5.9 | 0.007239 |
| P14873 | Microtubul | Map1b | 1373 | S | FEFSEAKI | -4.28 | 0.007703 |

| | | | | | | | |
|---|---|---|---|---|---|---|---|
| P14873 | Microtubul | Map1b | 1781 | S | VQSLEGE | -4.28 | 0.009824 |
| P14873 | Microtubul | Map1b | 614 | S | KPSVTEK | -3.79 | 0.014034 |
| P14873 | Microtubul | Map1b | 1260 | S | RLSPAKSI | -2.98 | 0.016869 |
| P14873 | Microtubul | Map1b | 1255 | S | 2DVSDER | -7.32 | 0.024414 |
| P14873 | Microtubul | Map1b | 1775 | S | SLASEKV( | -2.55 | 0.041388 |
| P20357 | Microtubul | Map2 | 1352 | S | 2EIEMAAE | -13.74 | 0.004577 |
| P20357 | Microtubul | Map2 | 1358 | T | 2EAQAEP | -13.97 | 0.00517 |
| P20357 | Microtubul | Map2 | 1609 | T | TSTPTTP( | -2.39 | 0.037851 |
| P10637 | Microtubul | Mapt | 692 | S | 2KTDHGA | -4.52 | 0.006328 |
| P10637 | Microtubul | Mapt | 494 | S | EPPKSGE | 3.8 | 0.044996 |
| P10637 | Microtubul | Mapt | 506 | S | SPGSPGT | 1.92 | 0.046621 |
| P26645 | Myristoylat | Marcks | 122 | S | EPAEPSS | 24.86 | 0.012515 |
| P04370-6 | Myelin bas | Mbp | 145 | S | GRGLSLS | 3.11 | 0.02821 |
| P13595 | Neural cell | Ncam1 | 770 | S | 2PGAKGK | -3.25 | 0.005144 |
| P13595 | Neural cell | Ncam1 | 774 | S | 2GKDMEE | -3.1 | 0.005455 |
| Q9QYG0 | Protein NE | Ndrg2 | 338 | S | 2TRLSRSI | -2.71 | 0.024835 |
| Q9QYG0 | Protein NE | Ndrg2 | 332 | S | 2MASSCM | -2.96 | 0.036015 |
| Q9QYG0 | Protein NE | Ndrg2 | 355 | S | DGSRSRS | 3.4 | 0.038051 |
| Q4KMM3 | Oxidation r | Oxr1 | 204 | S | STVSGIRI | -3.41 | 0.021344 |
| Q9Z0P4 | Paralemmi | Palm | 122 | S | ASKENSA | -13.09 | 0.009388 |
| Q9Z0P4 | Paralemmi | Palm | 124 | S | 2KENSAAI | -3.89 | 0.033751 |
| Q9Z0P4 | Paralemmi | Palm | 122 | S | 2ASKENSA | -3.79 | 0.043936 |
| Q9QYX7 | Protein pic | Pclo | 3376 | T | DDPRNLK | -4.5 | 0.006059 |
| Q8R3Q2 | Serine/thre | Ppp6r2 | 669 | S | TGSAVAR | -11.08 | 0.00744 |
| E9PUC5 | | Psd3 | 44 | S | IRRQSHR | 3.37 | 0.021953 |
| Q3TXS7 | 26S protea | Psmd1 | 315 | S | 2EEKTAS/ | -3.03 | 0.012915 |
| Q3TXS7 | 26S protea | Psmd1 | 311 | T | 2PMETEE | -3.18 | 0.018438 |
| Q99NE5 | Regulating | Rims1 | 1023 | S | SAPPSPLI | 3.12 | 0.03411 |
| Q8BP27 | Swi5-depe | Sfr1 | 71 | S | 2PAVPQT | -4.12 | 0.048129 |
| Q8VD37 | SH3-conta | Sgip1 | 259 | T | 2PTGTPPI | -25.58 | 0.000975 |
| Q8VD37 | SH3-conta | Sgip1 | 263 | T | 2PPPLPPI | -2.83 | 0.033432 |
| P53986 | Monocarbo | Slc16a1 | 491 | S | QSPQQHS | -3.35 | 0.00693 |
| Q8BJI1 | Sodium-de | Slc6a17 | 665 | S | TLSVSYKI | -2.47 | 0.025586 |
| Q61548 | Clathrin co | Snap91 | 321 | T | SPATTVTS | 2.35 | 0.03655 |
| O35526 | Syntaxin-1 | Stx1a | 10 | T | ______MK | -3.75 | 0.018241 |
| O08599 | Syntaxin-b | Stxbp1 | 594 | S | LDTLKKLI | -2.91 | 0.043615 |
| Q9JIS5 | Synaptic v | Sv2a | 127 | S | RMADGAI | -3.45 | 0.017434 |
| O88935 | Synapsin-1 | Syn1 | 568 | S | SPQRQAC | -4.16 | 0.039177 |
| A2A690 | Protein TA | Tanc2 | 404 | S | 2HGTRMF | -2.21 | 0.041336 |
| P70399 | Tumor sup | Tp53bp1 | 552 | S | EEDRENT | -7 | 0.003835 |
