## Supplementary table 2 for "Identifying JNK-regulated phosphoproteome markers of anxiety-like behaviour in mouse hippocampus"

| HIPPOCAMPUS | | | | | Responder | | | NonResponder | |
|---|---|---|---|---|---|---|---|---|---|
| Protein | Protein.names | Gene.nam | Amino.acid | AA | FC | ROTS p-va | ROTS FDR | FC | ROTS p-va |
| P42208 | Septin-2 | Septin4 | 218 | S | 1.3 | 0.997975 | 0.997984 | 2.7 | 0.014049 |
| P28661 | Septin-4 | Septin 4 | 325 | S | 1.1 | 0.96265 | 0.993884 | 4.1 | 0.032907 |
| P28661 | Septin-4 | Septin 4 | 68 | S | -1.3 | 0.854717 | 0.988571 | -1 | 0.883434 |
| Q9Z2Q6 | Septin-5 | Septin 5 | 225 | S | 2.9 | 0.566502 | 0.912975 | 7 | 0.066246 |
| Q9Z2Q6 | Septin-5 | Septin 5 | 327 | S | 1.1 | 0.782424 | 0.963975 | 1.5 | 0.12526 |
| Q8CHH9 | Septin-8 | Septin 8 | 10 | S | 1.9 | 0.21501 | 0.672078 | -1 | 0.453253 |
| Q8CHH9 | Septin-8 | Septin 8 | 4 | T | -3 | 0.371736 | 0.820796 | 2.9 | 0.170301 |
| E9Q3M9 | Capping protein inh | 2010300C | 612 | S | -1.4 | 0.264227 | 0.713499 | -1 | 0.207681 |
| E9Q3M9 | Capping protein inh | 2010300C | 507 | S | -1 | 0.730715 | 0.963975 | -1 | 0.059251 |
| E9Q3M9 | Capping protein inh | 2010300C | 189 | S | 1.1 | 0.682934 | 0.956522 | 1.5 | 0.189848 |
| E9Q3M9 | Capping protein inh | 2010300C | 961 | S | 1.5 | 0.833668 | 0.986874 | 2.5 | 0.127097 |
| Q3UHJ0 | AP2-associated pro | Aak1 | 618 | T | 1.5 | 0.276524 | 0.716535 | 1.8 | 0.15109 |
| Q3UHJ0 | AP2-associated pro | Aak1 | 604 | T | 2.2 | 0.113396 | 0.495238 | 1.7 | 0.216812 |
| Q3UHJ0 | AP2-associated pro | Aak1 | 621 | S | -35 | 0.102506 | 0.475 | 1.4 | 0.243357 |
| Q3UHJ0 | AP2-associated pro | Aak1 | 622 | S | -1.4 | 0.908204 | 0.992333 | -59 | 0.099394 |
| Q3UHJ0 | AP2-associated pro | Aak1 | 618 | T | -2.3 | 0.069644 | 0.407895 | -2 | 0.031759 |
| Q6P542 | ATP-binding casset | Abcf1 | 138 | S | -1 | 0.967108 | 0.993884 | 1 | 0.852593 |
| Q8CBW3 | Abl interactor 1 | Abi1 | 183 | S | 1.6 | 0.052488 | 0.366412 | 1.4 | 0.115243 |
| P62484 | Abl interactor 2 | Abi2 | 221 | S | 1.1 | 0.764844 | 0.963975 | -2 | 0.178077 |
| P62484 | Abl interactor 2 | Abi2 | 172 | T | 1.5 | 0.548658 | 0.902156 | 1.1 | 0.738852 |
| Q8K4G5 | Actin-binding LIM p | Ablim1 | 475 | S | 1.3 | 0.772898 | 0.963975 | 2.3 | 0.132433 |
| Q8BL65 | Actin-binding LIM p | Ablim2 | 477 | S | 1.3 | 0.481083 | 0.874773 | 1.2 | 0.384003 |
| Q8BL65 | Actin-binding LIM p | Ablim2 | 294 | S | -1.1 | 0.913512 | 0.992333 | 1.2 | 0.89555 |
| Q8BL65 | Actin-binding LIM p | Ablim2 | 475 | T | 1.3 | 0.481083 | 0.874773 | 1.2 | 0.384003 |
| Q99JY9 | Actin-related protei | Actr3 | 418 | S | 1 | 0.731669 | 0.963975 | -1 | 0.491663 |
| Q9R1V6-1 | Disintegrin and met | Adam22 | 884 | S | 1.2 | 0.892274 | 0.99118 | 2 | 0.024152 |
| Q9R1V6-1 | Disintegrin and met | Adam22 | 861 | S | 1.2 | 0.577813 | 0.912975 | -1 | 0.198016 |
| P51830 | Adenylate cyclase t | Adcy9 | 610 | S | 1 | 0.92904 | 0.993884 | -1 | 0.561915 |
| E9Q1K3;Q | Alpha-adducin | Add1 | 12 | S | 1.3 | 0.867176 | 0.988571 | 1.4 | 0.418019 |
| E9Q1K3;Q | Alpha-adducin | Add1 | 586 | S | 1.2 | 0.397695 | 0.836864 | -1 | 0.704403 |
| E9Q1K3;Q | Alpha-adducin | Add1 | 355 | S | 1.7 | 0.223017 | 0.674923 | 1.6 | 0.212078 |
| E9Q1K3;Q | Alpha-adducin | Add1 | 353 | S | 2 | 0.085227 | 0.436047 | 1.6 | 0.198732 |
| E9Q1K3;Q | Alpha-adducin | Add1 | 423 | S | 3.1 | 0.843055 | 0.988571 | -9 | 0.06754 |
| E9Q1K3;Q | Alpha-adducin | Add1 | 11 | T | -1.1 | 0.613317 | 0.926975 | 9.9 | 0.105506 |
| E9Q1K3 | | Add1 | 639 | T | 1.4 | 0.345966 | 0.789352 | 1.1 | 0.515269 |
| E9Q1K3;Q | Alpha-adducin | Add1 | 358 | T | 2 | 0.162035 | 0.590551 | 1.3 | 0.830454 |
| Q9QYC0-2 | Alpha-adducin | Add1 | 610 | T | 1.4 | 0.345966 | 0.789352 | 1.1 | 0.515269 |
| Q9QYC0 | Alpha-adducin | Add1 | 610 | T | 1.4 | 0.345966 | 0.789352 | 1.1 | 0.515269 |
| E9Q1K3 | | Add1 | 629 | S | -1.9 | 0.028434 | 0.306667 | -2 | 0.006417 |
| E9Q1K3;Q | Alpha-adducin | Add1 | 355 | S | -1.8 | 0.065194 | 0.381944 | -2 | 0.018751 |
| E9Q1K3;Q | Alpha-adducin | Add1 | 364 | S | 2.6 | 0.21389 | 0.67101 | -7 | 0.465557 |
| E9Q1K3;Q | Alpha-adducin | Add1 | 353 | S | -1.2 | 0.785296 | 0.963975 | -2 | 0.885576 |
| Q9QYC0-2 | Alpha-adducin | Add1 | 600 | S | -1.9 | 0.028434 | 0.306667 | -2 | 0.006417 |
| Q9QYC0 | Alpha-adducin | Add1 | 600 | S | -1.9 | 0.028434 | 0.306667 | -2 | 0.006417 |
| Q9QYC0 | Alpha-adducin | Add1 | 605 | S | -1.3 | 0.689651 | 0.956643 | -2 | 0.714098 |
| Q9QYC0 | Alpha-adducin | Add1 | 613 | S | 1.5 | 0.543188 | 0.902156 | 1.4 | 0.279831 |
| E9Q1K3 | | Add1 | 639 | T | -18 | 0.008868 | 0.195652 | -3 | 0.216655 |
| E9Q1K3;Q | Alpha-adducin | Add1 | 358 | T | -1.8 | 0.065194 | 0.381944 | -2 | 0.018751 |
| Q9QYC0-2 | Alpha-adducin | Add1 | 610 | T | -22 | 0.012245 | 0.195652 | -3 | 0.337769 |
| Q9QYC0 | Alpha-adducin | Add1 | 610 | T | -19 | 0.007892 | 0.195652 | -3 | 0.230601 |
| Q9QYC0 | Alpha-adducin | Add1 | 614 | T | 1.5 | 0.543188 | 0.902156 | 1.1 | 0.62182 |
| Q9QYC0 | Alpha-adducin | Add1 | 600 | S | -1.8 | 0.313133 | 0.753027 | -1 | 0.794974 |
| Q9QYC0 | Alpha-adducin | Add1 | 605 | S | -1.3 | 0.355539 | 0.809524 | -3 | 0.173624 |
| Q9QYC0 | Alpha-adducin | Add1 | 613 | S | -1.8 | 0.373243 | 0.8234 | -1 | 0.862055 |
| Q9QYC0 | Alpha-adducin | Add1 | 610 | T | -1.8 | 0.343366 | 0.787383 | -1 | 0.865667 |
| Q9QYC0 | Alpha-adducin | Add1 | 614 | T | -1.8 | 0.331326 | 0.77619 | -1 | 0.858118 |
| Q9QYB8 | Beta-adducin | Add2 | 614 | S | 1.1 | 0.616419 | 0.926975 | -6 | 0.041013 |

| | | | | | | | | | |
|---|---|---|---|---|---|---|---|---|---|
| Q9QYB8 | Beta-adducin | Add2 | 618 | S | 1.5 | 0.475125 | 0.874773 | 1.4 | 0.226497 |
| Q9QYB8 | Beta-adducin | Add2 | 594 | S | -1.2 | 0.589939 | 0.92272 | 1.1 | 0.808073 |
| Q9QYB8 | Beta-adducin | Add2 | 11 | S | 1.6 | 0.053914 | 0.366412 | -1 | 0.523589 |
| Q9QYB8 | Beta-adducin | Add2 | 602 | S | 1.3 | 0.436587 | 0.848425 | -1 | 0.502107 |
| Q9QYB8 | Beta-adducin | Add2 | 692 | S | -1 | 0.942582 | 0.993884 | -2 | 0.061952 |
| Q9QYB8 | Beta-adducin | Add2 | 702 | S | 1 | 0.843539 | 0.988571 | -1 | 0.158905 |
| Q9QYB8 | Beta-adducin | Add2 | 614 | S | 1.4 | 0.372015 | 0.820796 | -1 | 0.726632 |
| Q9QYB8 | Beta-adducin | Add2 | 618 | S | -1.1 | 0.660625 | 0.940678 | -1 | 0.181535 |
| Q9QYB8 | Beta-adducin | Add2 | 620 | S | -6.2 | 0.202433 | 0.65894 | 1.6 | 0.802811 |
| Q9QYB8 | Beta-adducin | Add2 | 602 | S | -1.2 | 0.780431 | 0.963975 | -2 | 0.884295 |
| Q9QYB8 | Beta-adducin | Add2 | 606 | S | -1.2 | 0.716444 | 0.963975 | -2 | 0.985331 |
| Q9QYB8 | Beta-adducin | Add2 | 692 | S | -2.5 | 0.007242 | 0.195652 | -3 | 0.002199 |
| Q9QYB8 | Beta-adducin | Add2 | 696 | S | -1.9 | 0.494017 | 0.879061 | -2 | 0.855844 |
| Q9QYB8 | Beta-adducin | Add2 | 700 | S | -1.8 | 0.59408 | 0.92272 | -2 | 0.799118 |
| Q9QYB8 | Beta-adducin | Add2 | 612 | T | 4.1 | 0.185925 | 0.637681 | -15 | 0.288371 |
| Q9QYB5 | Gamma-adducin | Add3 | 673 | S | 1.7 | 0.065271 | 0.381944 | 1.7 | 0.00281 |
| Q9QYB5 | Gamma-adducin | Add3 | 677 | S | 1.2 | 0.54856 | 0.902156 | -1 | 0.623511 |
| Q9QYB5 | Gamma-adducin | Add3 | 681 | S | 1.5 | 0.033015 | 0.311111 | -3 | 0.02495 |
| Q9QYB5 | Gamma-adducin | Add3 | 683 | S | 2 | 0.13089 | 0.530973 | 1.1 | 0.988062 |
| Q9QYB5 | Gamma-adducin | Add3 | 11 | T | 4 | 0.002436 | 0.105263 | 1.9 | 0.161614 |
| Q9QYB5 | Gamma-adducin | Add3 | 677 | S | 1.1 | 0.438875 | 0.848425 | -1 | 0.593867 |
| Q9QYB5 | Gamma-adducin | Add3 | 681 | S | -1.7 | 0.028932 | 0.311111 | -2 | 0.003399 |
| Q3UNS7 | | Adgrg4 | 15 | S | 32 | 0.013072 | 0.211538 | 9.8 | 0.132902 |
| Q68FL4 | Putative adenosylhc | Ahcyl2 | 109 | S | 1.3 | 0.279234 | 0.716535 | -1 | 0.279746 |
| Q9WTQ5 | A-kinase anchor prc | Akap12 | 584 | S | -2 | 0.135971 | 0.536797 | -2 | 0.047336 |
| D3YVF0 | A-kinase anchor prc | Akap5 | 251 | S | -1 | 0.857709 | 0.988571 | 1.1 | 0.726469 |
| D3YVF0 | A-kinase anchor prc | Akap5 | 22 | S | 1 | 0.784539 | 0.963975 | 1.3 | 0.323703 |
| D3YVF0 | A-kinase anchor prc | Akap5 | 220 | T | 1.2 | 0.765188 | 0.963975 | 1.2 | 0.281153 |
| P05064 | Fructose-bisphosph | Aldoa | 36 | S | 1.5 | 0.246636 | 0.699708 | 1.3 | 0.415625 |
| P05064 | Fructose-bisphosph | Aldoa | 39 | S | 1.4 | 0.379904 | 0.828326 | -1 | 0.855197 |
| Q7TQF7 | Amphiphysin | Amph | 500 | S | -1.1 | 0.798304 | 0.967352 | -2 | 0.103785 |
| Q7TQF7 | Amphiphysin | Amph | 250 | S | -1.8 | 0.732708 | 0.963975 | 1.5 | 0.953811 |
| Q7TQF7 | Amphiphysin | Amph | 252 | S | -1.1 | 0.732839 | 0.963975 | 1.1 | 0.975927 |
| Q8C8R3 | Ankyrin-2 | Ank2 | 3362 | S | 1.3 | 0.433101 | 0.848425 | 3.4 | 0.004156 |
| Q8C8R3 | Ankyrin-2 | Ank2 | 1426 | S | 1.1 | 0.726367 | 0.963975 | -1 | 0.84741 |
| Q8C8R3 | Ankyrin-2 | Ank2 | 1428 | S | 1 | 0.748127 | 0.963975 | -1 | 0.938353 |
| Q8C8R3;Q | Ankyrin-2 | Ank2 | 3774 | S | 1.1 | 0.627335 | 0.927536 | -1 | 0.651821 |
| Q8C8R3;Q | Ankyrin-2 | Ank2 | 3781 | S | -1.6 | 0.759504 | 0.963975 | -4 | 0.328377 |
| Q8C8R3 | Ankyrin-2 | Ank2 | 2364 | S | -1.6 | 0.075363 | 0.414013 | -2 | 0.013663 |
| Q8C8R3;Q | Ankyrin-2 | Ank2 | 3692 | S | 1.3 | 0.288123 | 0.718346 | -1 | 0.658505 |
| Q8C8R3-3 | Ankyrin-2 | Ank2 | 588 | S | 1.1 | 0.646557 | 0.929088 | -1 | 0.936728 |
| Q8C8R3-3 | Ankyrin-2 | Ank2 | 590 | S | 1 | 0.758231 | 0.963975 | -1 | 0.916946 |
| Q8C8R3;Q | Ankyrin-2 | Ank2 | 3785 | T | 4 | 0.013566 | 0.211538 | 2.7 | 0.235971 |
| Q8C8R3 | Ankyrin-2 | Ank2 | 3050 | T | -2.1 | 0.056334 | 0.366412 | -1 | 0.84817 |
| Q8C8R3 | Ankyrin-2 | Ank2 | 2232 | T | 1.1 | 0.660469 | 0.940678 | -1 | 0.4731 |
| Q8C8R3 | Ankyrin-2 | Ank2 | 1699 | S | -5.2 | 0.035599 | 0.329787 | 1.5 | 0.207147 |
| Q8C8R3 | Ankyrin-2 | Ank2 | 1700 | S | -4.8 | 0.054711 | 0.366412 | 1.5 | 0.209675 |
| Q8C8R3;Q | Ankyrin-2 | Ank2 | 3692 | S | -1.4 | 0.167689 | 0.6 | -2 | 0.025448 |
| Q6PD24 | Ankyrin repeat dom | Ankrd13d | 552 | S | -1.3 | 0.282236 | 0.718016 | 1.1 | 0.464955 |
| Q6PD24 | Ankyrin repeat dom | Ankrd13d | 556 | T | -1.2 | 0.860404 | 0.988571 | 1.3 | 0.430921 |
| B2RW11 | | Ankrd34a | 460 | S | 1.1 | 0.688739 | 0.956643 | 1.6 | 0.031858 |
| P10107 | Annexin A1 | Anxa1 | 5 | S | -6.8 | 0.180372 | 0.632353 | -2 | 0.567185 |
| Q9JME5 | AP-3 complex subu | Ap3b2 | 272 | S | 1.7 | 0.805781 | 0.967352 | 2 | 0.837072 |
| O54774 | AP-3 complex subu | Ap3d1 | 760 | S | 1.4 | 0.303841 | 0.748744 | 1.2 | 0.908855 |
| O54774 | AP-3 complex subu | Ap3d1 | 825 | S | 3.2 | 0.126861 | 0.52968 | 4.2 | 0.109582 |
| O54774 | AP-3 complex subu | Ap3d1 | 784 | S | 1.9 | 0.615668 | 0.926975 | 2.8 | 0.060704 |
| O54774 | AP-3 complex subu | Ap3d1 | 755 | S | -1.9 | 0.603542 | 0.926154 | -1 | 0.981592 |
| O54774 | AP-3 complex subu | Ap3d1 | 760 | S | -2.4 | 0.221284 | 0.674923 | -2 | 0.4984 |
| B2RUJ5 | Amyloid beta A4 pre | Apba1 | 267 | S | 1.2 | 0.57486 | 0.912975 | 1.3 | 0.30134 |

| | | | | | | | | | |
|---|---|---|---|---|---|---|---|---|---|
| E9Q414 | Apolipoprotein B-10 | Apob | 3477 | S | -1.2 | 0.965613 | 0.993884 | -1 | 0.818303 |
| E9Q414 | Apolipoprotein B-10 | Apob | 3483 | S | -1.2 | 0.980705 | 0.993884 | -1 | 0.829277 |
| E9Q414 | Apolipoprotein B-10 | Apob | 3486 | S | -1.2 | 0.867242 | 0.988571 | -1 | 0.751063 |
| Q9EPJ9 | ADP-ribosylation fa | Arfgap1 | 342 | S | -1.1 | 0.914024 | 0.992333 | 1.2 | 0.462138 |
| Q9EPJ9 | ADP-ribosylation fa | Arfgap1 | 135 | T | 1.9 | 0.040719 | 0.356436 | 2.1 | 0.023653 |
| A2A5R2 | Brefeldin A-inhibitec | Arfgef2 | 218 | S | -1.8 | 0.076621 | 0.421384 | -4 | 0.003683 |
| A2A5R2 | Brefeldin A-inhibitec | Arfgef2 | 227 | S | -1.9 | 0.062495 | 0.381944 | -4 | 0.004927 |
| Q5FWK3 | Rho GTPase-activa | Arhgap1 | 47 | S | -3 | 0.001249 | 0.090909 | -2 | 0.002115 |
| Q5FWK3 | Rho GTPase-activa | Arhgap1 | 51 | S | -3 | 0.001275 | 0.090909 | -3 | 0.006026 |
| Q8C0D4 | Rho GTPase-activa | Arhgap12 | 238 | S | -1.1 | 0.889444 | 0.99118 | -1 | 0.719564 |
| Q8C0D4 | Rho GTPase-activa | Arhgap12 | 228 | T | -1.1 | 0.889265 | 0.99118 | -1 | 0.719487 |
| Q8C0D4 | Rho GTPase-activa | Arhgap12 | 229 | T | -1.1 | 0.968315 | 0.993884 | -1 | 0.763022 |
| Q6DFV3 | Rho GTPase-activa | Arhgap21 | 41 | S | 1.3 | 0.448327 | 0.859073 | 1.4 | 0.115771 |
| Q69ZH9 | Rho GTPase-activa | Arhgap23 | 361 | S | -1.1 | 0.790821 | 0.966952 | -1 | 0.989314 |
| Q811P8 | Rho GTPase-activa | Arhgap32 | 952 | S | 1.5 | 0.041762 | 0.356436 | 1.7 | 0.026422 |
| Q811P8 | Rho GTPase-activa | Arhgap32 | 706 | S | 1.4 | 0.356815 | 0.809524 | 2.1 | 0.83471 |
| Q91YM2 | Rho GTPase-activa | Arhgap35 | 1179 | S | 1.2 | 0.632964 | 0.927536 | 1.4 | 0.279133 |
| P59281 | Rho GTPase-activa | Arhgap39 | 597 | S | 1.2 | 0.360788 | 0.810811 | -1 | 0.9552 |
| Q60875 | Rho guanine nuclec | Arhgef2 | 931 | S | -1 | 0.799405 | 0.967352 | -2 | 0.010167 |
| Q9ES28 | Rho guanine nuclec | Arhgef7 | 228 | S | 1.3 | 0.26622 | 0.713499 | 1.7 | 0.0008 |
| Q9ES28 | Rho guanine nuclec | Arhgef7 | 497 | S | 1.4 | 0.073468 | 0.410256 | -2 | 0.078613 |
| Q9ES28 | Rho guanine nuclec | Arhgef7 | 155 | S | 2 | 0.095351 | 0.47027 | 1.3 | 0.354385 |
| Q9D0L7 | Armadillo repeat-co | Armc10 | 43 | S | 1.1 | 0.668675 | 0.940678 | 2.9 | 0.221138 |
| Q7SIG6 | Arf-GAP with SH3 c | Asap2 | 704 | S | -1.1 | 0.985618 | 0.994914 | 1.5 | 0.27798 |
| Q8BHE3 | Caytaxin | Atcay | 44 | S | -1 | 0.369283 | 0.819599 | 1.6 | 0.041532 |
| Q8BHE3 | Caytaxin | Atcay | 46 | S | -2.1 | 0.972016 | 0.993884 | 5.3 | 0.2095 |
| Q8VDN2 | Sodium/potassium-' | Atp1a1 | 228 | S | -1.2 | 0.407097 | 0.84696 | 1.1 | 0.574515 |
| Q6PIC6;Q: | Sodium/potassium-' | Atp1a3 | 217 | S | -1.1 | 0.729671 | 0.963975 | -2 | 0.431308 |
| A2ALL9 | Calcium-transportin | Atp2b3 | 1124 | S | 2.4 | 0.182054 | 0.6337 | -1 | 0.812369 |
| P70704 | Phospholipid-transp | Atp8a1 | 25 | S | -3.8 | 0.009315 | 0.195652 | -4 | 0.006803 |
| P70704 | Phospholipid-transp | Atp8a1 | 29 | S | -3.6 | 0.010595 | 0.195652 | -4 | 0.00633 |
| Q7TQH0 | Ataxin-2-like proteir | Atxn2l | 597 | S | 1.3 | 0.292926 | 0.728205 | 1.6 | 0.117161 |
| Q3UHD1 | Brain-specific angio | Bai1 | 1467 | S | -1.6 | 0.327924 | 0.768496 | -1 | 0.974296 |
| Q8BKX1 | Brain-specific angio | Baiap2 | 326 | S | 1.1 | 0.898933 | 0.99118 | -1 | 0.097142 |
| Q80TT2 | BAI1-associated pro | Baiap3 | 78 | S | 1.8 | 0.732076 | 0.963975 | -1 | 0.778103 |
| Q91XV3 | Brain acid soluble p | Basp1 | 92 | S | 1.3 | 0.54398 | 0.902156 | 2.2 | 0.222719 |
| Q91XV3 | Brain acid soluble p | Basp1 | 31 | T | -13 | 0.5662 | 0.912975 | 2.2 | 0.763358 |
| Q91XV3 | Brain acid soluble p | Basp1 | 36 | T | -1.3 | 0.485935 | 0.874773 | 2.1 | 0.928397 |
| Q91XV3 | Brain acid soluble p | Basp1 | 40 | S | -1.2 | 0.622404 | 0.927083 | -2 | 0.523564 |
| Q91XV3 | Brain acid soluble p | Basp1 | 31 | T | -2.1 | 0.035736 | 0.329787 | -1 | 0.568808 |
| Q91XV3 | Brain acid soluble p | Basp1 | 36 | T | -3.3 | 0.177733 | 0.62963 | -1 | 0.840556 |
| A2AUY4 | | Baz2b | 2093 | T | 1.5 | 0.856345 | 0.988571 | -2 | 0.414226 |
| Q8K019 | Bcl-2-associated tra | Bclaf1 | 494 | S | 1.2 | 0.280758 | 0.717277 | -1 | 0.23064 |
| Q8K019 | Bcl-2-associated tra | Bclaf1 | 656 | S | 1.1 | 0.543308 | 0.902156 | 1.5 | 0.061795 |
| Q68EF6 | Brain-enriched guar | Begain | 229 | S | -1 | 0.967524 | 0.993884 | 1.2 | 0.795252 |
| Q68EF6 | Brain-enriched guar | Begain | 246 | S | 1 | 0.948581 | 0.993884 | 1.4 | 0.257309 |
| O08539 | Myc box-dependent | Bin1 | 296 | S | 1.5 | 0.528877 | 0.894378 | -2 | 0.665549 |
| O08539 | Myc box-dependent | Bin1 | 298 | S | -1 | 0.990585 | 0.995947 | 1.4 | 0.358585 |
| O08539 | Myc box-dependent | Bin1 | 296 | S | -2.6 | 0.00096 | 0.090909 | -1 | 0.021412 |
| O08539 | Myc box-dependent | Bin1 | 298 | S | -2.2 | 0.765614 | 0.963975 | -1 | 0.783911 |
| O08539 | Myc box-dependent | Bin1 | 296 | S | -4.6 | 0.007218 | 0.195652 | -2 | 0.502897 |
| O08539 | Myc box-dependent | Bin1 | 298 | S | -3.7 | 0.015725 | 0.224138 | -2 | 0.502897 |
| O08539 | Myc box-dependent | Bin1 | 304 | S | -3.8 | 0.016992 | 0.225806 | -2 | 0.502897 |
| P28028 | Serine/threonine-pr | Braf | 431 | S | 1.3 | 0.608584 | 0.926718 | 2.2 | 0.109054 |
| P28028 | Serine/threonine-pr | Braf | 432 | S | 1.2 | 0.831553 | 0.986874 | 2.1 | 0.110994 |
| P28028 | Serine/threonine-pr | Braf | 714 | S | 1.2 | 0.252983 | 0.704871 | -1 | 0.060011 |
| G3UWV4 | Serine/threonine-pr | Brsk2 | 512 | S | 1.2 | 0.60888 | 0.926718 | -1 | 0.457898 |
| G3UWV4 | | Brsk2 | 417 | S | 1.4 | 0.911913 | 0.992333 | -1 | 0.791158 |
| O88737 | Protein bassoon | Bsn | 1482 | S | -2.1 | 0.278479 | 0.716535 | 1.8 | 0.462949 |

| | | | | | | | | | |
|---|---|---|---|---|---|---|---|---|---|
| O88737 | Protein bassoon | Bsn | 105 | S | 1.2 | 0.665754 | 0.940678 | 1.2 | 0.459281 |
| O88737 | Protein bassoon | Bsn | 1114 | S | 1 | 0.769591 | 0.963975 | 1.3 | 0.260751 |
| O88737 | Protein bassoon | Bsn | 2858 | S | -1 | 0.755691 | 0.963975 | 1 | 0.356132 |
| O88737 | Protein bassoon | Bsn | 2860 | S | 1.1 | 0.666034 | 0.940678 | 1 | 0.793067 |
| O88737 | Protein bassoon | Bsn | 3512 | S | -1.7 | 0.501149 | 0.880143 | -1 | 0.901492 |
| O88737 | Protein bassoon | Bsn | 1236 | S | -4.8 | 0.047443 | 0.366412 | -2 | 0.305332 |
| O88737 | Protein bassoon | Bsn | 2595 | T | -1.8 | 0.023573 | 0.279412 | -3 | 0.003017 |
| O88737 | Protein bassoon | Bsn | 1491 | S | -1.2 | 0.730498 | 0.963975 | 1.1 | 0.461806 |
| O88737 | Protein bassoon | Bsn | 1108 | S | -2.5 | 0.091786 | 0.466667 | -2 | 0.593196 |
| O88737 | Protein bassoon | Bsn | 1114 | S | -2.5 | 0.079456 | 0.427711 | -2 | 0.593196 |
| O88737 | Protein bassoon | Bsn | 2860 | S | 1.2 | 0.288785 | 0.719072 | -1 | 0.809004 |
| O88737 | Protein bassoon | Bsn | 2866 | S | -1.5 | 0.994113 | 0.997978 | -2 | 0.340283 |
| O88737 | Protein bassoon | Bsn | 3512 | S | -1.4 | 0.738714 | 0.963975 | -1 | 0.623877 |
| O88737 | Protein bassoon | Bsn | 3519 | S | -2 | 0.003244 | 0.105263 | -2 | 0.002021 |
| O88737 | Protein bassoon | Bsn | 1108 | S | -2.7 | 0.436072 | 0.848425 | -1 | 0.806364 |
| O88737 | Protein bassoon | Bsn | 1114 | S | -2.8 | 0.40885 | 0.84728 | -2 | 0.690327 |
| O88737 | Protein bassoon | Bsn | 1102 | T | -2.6 | 0.44039 | 0.848425 | -2 | 0.861309 |
| Q80X80 | C2 domain-containi | C2cd2l | 376 | S | 2.7 | 0.086982 | 0.44382 | 5.4 | 0.001295 |
| Q80X80 | C2 domain-containi | C2cd2l | 464 | S | -2 | 0.01767 | 0.225806 | -2 | 0.058996 |
| Q80X80 | C2 domain-containi | C2cd2l | 470 | S | -1.6 | 0.793155 | 0.966952 | -1 | 0.992539 |
| Q7TPS5 | C2 domain-containi | C2cd5 | 262 | S | 1.3 | 0.504079 | 0.883186 | 1.2 | 0.442441 |
| Q8R464 | Cell adhesion mole | Cadm4 | 361 | S | 1.6 | 0.561759 | 0.912975 | 2.4 | 0.080947 |
| A0A286YC | Calcium/calmodulin | Camk2a | 234 | S | 1.9 | 0.03121 | 0.311111 | 2.4 | 0.003806 |
| P11798 | Calcium/calmodulin | Camk2a | 330 | S | 1.1 | 0.88865 | 0.99118 | -2 | 0.574816 |
| P11798 | Calcium/calmodulin | Camk2a | 331 | S | 2.5 | 0.049515 | 0.366412 | -1 | 0.695463 |
| P11798 | Calcium/calmodulin | Camk2a | 337 | T | 1.8 | 0.222818 | 0.674923 | 1.2 | 0.539022 |
| P11798 | Calcium/calmodulin | Camk2a | 286 | T | -1.2 | 0.935536 | 0.993884 | -1 | 0.874493 |
| P11798 | Calcium/calmodulin | Camk2a | 331 | S | -1.8 | 0.493228 | 0.879061 | -1 | 0.844845 |
| P11798 | Calcium/calmodulin | Camk2a | 333 | S | -1.7 | 0.553757 | 0.910744 | -1 | 0.916994 |
| P11798 | Calcium/calmodulin | Camk2a | 336 | T | 1.7 | 0.350016 | 0.799539 | 1.6 | 0.601612 |
| P11798 | Calcium/calmodulin | Camk2a | 337 | T | -3.9 | 0.003793 | 0.136364 | -1 | 0.00914 |
| P28652 | Calcium/calmodulin | Camk2b | 395 | S | 1.4 | 0.896877 | 0.99118 | -2 | 0.476864 |
| P28652 | Calcium/calmodulin | Camk2b | 397 | S | 1.5 | 0.669679 | 0.940678 | 1.1 | 0.823372 |
| Q8VBY2 | Calcium/calmodulin | Camkk1 | 458 | S | 1.1 | 0.716262 | 0.963975 | 1.1 | 0.584831 |
| Q3UHL1 | CaM kinase-like ve | Camkv | 446 | T | 1.3 | 0.524865 | 0.894378 | -2 | 0.178651 |
| P35564 | Calnexin | Canx | 582 | S | -1.3 | 0.38345 | 0.828326 | 1.4 | 0.1915 |
| P35564 | Calnexin | Canx | 553 | S | -1.2 | 0.564555 | 0.912975 | 1.1 | 0.54695 |
| P35564 | Calnexin | Canx | 563 | S | 1.2 | 0.59551 | 0.92272 | 2.5 | 0.067782 |
| P35564 | Calnexin | Canx | 553 | S | -3.1 | 0.030075 | 0.311111 | -2 | 0.215503 |
| P35564 | Calnexin | Canx | 563 | S | -3.1 | 0.030075 | 0.311111 | -2 | 0.215503 |
| Q9CYT6 | Adenylyl cyclase-as | Cap2 | 308 | S | -1.6 | 0.061949 | 0.381944 | -2 | 0.009734 |
| Q9CYT6 | Adenylyl cyclase-as | Cap2 | 311 | S | -1.6 | 0.061949 | 0.381944 | -2 | 0.009734 |
| O70589-3 | | Cask | 570 | S | 2 | 0.073753 | 0.410256 | -6 | 0.317775 |
| O70589-3 | | Cask | 571 | S | -6.2 | 0.244122 | 0.699115 | -1 | 0.818852 |
| Q6P9K8 | Caskin-1 | Caskin1 | 1363 | S | -1.3 | 0.33477 | 0.786223 | 1.3 | 0.280928 |
| Q6P9K8 | Caskin-1 | Caskin1 | 1259 | S | 1.4 | 0.528797 | 0.894378 | 1.5 | 0.001536 |
| Q6P9K8 | Caskin-1 | Caskin1 | 365 | S | -1.1 | 0.95876 | 0.993884 | 1 | 0.737592 |
| Q3TVA9 | Coiled-coil domain- | Ccdc136 | 50 | S | -1.3 | 0.575251 | 0.912975 | 1 | 0.815891 |
| Q8BGU5 | Cyclin-Y | Ccny | 326 | S | 1.7 | 0.227275 | 0.674923 | -1 | 0.988289 |
| Q99L43 | Phosphatidate cytid | Cds2 | 32 | S | -2.2 | 0.364213 | 0.81573 | -5 | 0.072778 |
| Q99L43 | Phosphatidate cytid | Cds2 | 30 | T | 1.5 | 0.757541 | 0.963975 | -2 | 0.275871 |
| Q9JKC6 | Cell cycle exit and r | Cend1 | 89 | S | 21 | 0.068124 | 0.401361 | 45 | 0.023429 |
| Q9JKC6 | Cell cycle exit and r | Cend1 | 90 | S | -2.3 | 0.490064 | 0.874773 | 2.9 | 0.987745 |
| Q9JKC6 | Cell cycle exit and r | Cend1 | 94 | T | -3.4 | 0.049564 | 0.366412 | -11 | 0.003118 |
| P18760 | Cofilin-1 | Cfl1 | 3 | S | -5.9 | 0.04217 | 0.356436 | -6 | 0.049406 |
| Q60771 | Claudin-11 | Cldn11 | 196 | S | 14 | 0.046049 | 0.366412 | 8.5 | 0.458059 |
| Q60771 | Claudin-11 | Cldn11 | 198 | S | -9.5 | 0.371576 | 0.820796 | 1.5 | 0.852769 |
| Q60771 | Claudin-11 | Cldn11 | 196 | S | -9.1 | 0.001161 | 0.090909 | -8 | 0.001686 |
| Q80YA9 | Connector enhance | Cnksr2 | 390 | S | 1.2 | 0.842835 | 0.988571 | 1.4 | 0.40905 |

| | | | | | | | | | |
|---|---|---|---|---|---|---|---|---|---|
| Q80YA9 | Connector enhance | Cnksr2 | 906 | S | -1 | 0.729537 | 0.963975 | 1.8 | 0.047503 |
| O54991 | Contactin-associate | Cntnap1 | 1382 | S | 1.3 | 0.716188 | 0.963975 | -4 | 0.28284 |
| O54991 | Contactin-associate | Cntnap1 | 1384 | S | -1.3 | 0.284199 | 0.718346 | 1.1 | 0.614413 |
| O88587 | Catechol O-methylt | Comt | 261 | S | 1.3 | 0.723952 | 0.963975 | 1.1 | 0.555696 |
| Q9Z140 | Copine-6 | Cpne6 | 556 | S | -1.1 | 0.932567 | 0.993884 | -2 | 0.054169 |
| Q9Z140 | Copine-6 | Cpne6 | 68 | S | -1.3 | 0.597652 | 0.92272 | -1 | 0.374422 |
| Q9Z140 | Copine-6 | Cpne6 | 330 | S | -1.2 | 0.730258 | 0.963975 | -1 | 0.907058 |
| P97427 | Dihydropyrimidinase | Crmp1 | 524 | S | -19 | 0.096975 | 0.470588 | 2.1 | 0.610054 |
| P97427 | Dihydropyrimidinase | Crmp1 | 542 | S | -1.2 | 0.874645 | 0.99118 | -2 | 0.474765 |
| P97427 | Dihydropyrimidinase | Crmp1 | 8 | S | 1.1 | 0.804261 | 0.967352 | 1.2 | 0.39449 |
| P97427 | Dihydropyrimidinase | Crmp1 | 509 | T | 2.2 | 0.017111 | 0.225806 | 1.3 | 0.234605 |
| P97427 | Dihydropyrimidinase | Crmp1 | 521 | S | -1.8 | 0.082375 | 0.429412 | -2 | 0.216589 |
| P97427 | Dihydropyrimidinase | Crmp1 | 522 | S | -1.8 | 0.082375 | 0.429412 | -2 | 0.216589 |
| P97427 | Dihydropyrimidinase | Crmp1 | 524 | S | -1.8 | 0.082375 | 0.429412 | -2 | 0.216589 |
| P97315 | Cysteine and glycin | Csrp1 | 192 | S | -1 | 0.785195 | 0.963975 | 2.1 | 0.018222 |
| P26231 | Catenin alpha-1 | Ctnna1 | 641 | S | -1.3 | 0.900203 | 0.99118 | 2 | 0.146726 |
| Q61301 | Catenin alpha-2 | Ctnna2 | 262 | S | 1.1 | 0.663024 | 0.940678 | 1.1 | 0.337368 |
| Q61301 | Catenin alpha-2 | Ctnna2 | 640 | S | -2.6 | 0.032031 | 0.311111 | 1.4 | 0.188586 |
| Q61301;P2 | Catenin alpha-2;Ca | Ctnna2 | 654 | S | -2.2 | 0.032005 | 0.311111 | -2 | 0.080291 |
| Q61301;P2 | Catenin alpha-2;Ca | Ctnna2 | 657 | T | -2.3 | 0.031271 | 0.311111 | -1 | 0.224143 |
| Q02248 | Catenin beta-1 | Ctnnb1 | 552 | S | 2 | 0.078169 | 0.427711 | 1.9 | 0.104533 |
| O35927 | Catenin delta-2 | Ctnnd2 | 412 | S | 1.6 | 0.090641 | 0.463687 | 2.7 | 0.004339 |
| O35927 | Catenin delta-2 | Ctnnd2 | 264 | S | 1.1 | 0.917067 | 0.993884 | 1 | 0.753988 |
| O35927 | Catenin delta-2 | Ctnnd2 | 273 | S | 1.1 | 0.662043 | 0.940678 | -1 | 0.204544 |
| O35927 | Catenin delta-2 | Ctnnd2 | 458 | S | -2.5 | 0.054657 | 0.366412 | -1 | 0.447582 |
| O35927 | Catenin delta-2 | Ctnnd2 | 264 | S | -2.9 | 0.009352 | 0.195652 | -2 | 0.031392 |
| O35927 | Catenin delta-2 | Ctnnd2 | 268 | T | -2.3 | 0.101285 | 0.471795 | -1 | 0.633875 |
| Q60598 | Src substrate cortac | Cttn | 407 | S | 1.1 | 0.836455 | 0.988571 | -1 | 0.590099 |
| Q60598 | Src substrate cortac | Cttn | 401 | T | 3.8 | 0.124056 | 0.520737 | 4.2 | 0.159806 |
| Q60598 | Src substrate cortac | Cttn | 401 | T | 1.4 | 0.794328 | 0.966952 | 1.5 | 0.319591 |
| Q60598 | Src substrate cortac | Cttn | 421 | Y | 1.1 | 0.92438 | 0.993884 | 3.4 | 0.298869 |
| D3YZ21 | | D430041D | 899 | S | -1.4 | 0.253418 | 0.704871 | 1.3 | 0.453292 |
| A0A0R4J1 | Drebrin | Dbn1 | 142 | S | 1.5 | 0.233968 | 0.690909 | 1.7 | 0.010603 |
| Q9QXS6 | Drebrin | Dbn1 | 344 | S | 1.5 | 0.343587 | 0.787383 | -1 | 0.815046 |
| Q9QXS6 | Drebrin | Dbn1 | 656 | S | -4.6 | 0.099285 | 0.471795 | 1.5 | 0.368221 |
| Q8N7N5 | DDB1- and CUL4-a | Dcaf8 | 100 | S | 1.3 | 0.301161 | 0.743655 | 1.4 | 0.091754 |
| Q9JLM8 | Serine/threonine-pr | Dclk1 | 352 | S | 1.6 | 0.419152 | 0.848425 | 1.9 | 0.4456 |
| Q9JLM8 | Serine/threonine-pr | Dclk1 | 392 | S | 1.3 | 0.644806 | 0.927536 | 2.7 | 0.016735 |
| Q9JLM8 | Serine/threonine-pr | Dclk1 | 337 | S | -1.3 | 0.805128 | 0.967352 | 1.1 | 0.661912 |
| Q9CYC6 | m7GpppN-mRNA h | Dcp2 | 247 | S | 1.1 | 0.796178 | 0.966952 | -3 | 0.384291 |
| O70133-2 | | Dhx9 | 137 | S | 1.7 | 0.028722 | 0.306667 | -1 | 0.725839 |
| Q8CI75 | DIS3-like exonuclea | Dis3l2 | 864 | S | -1.6 | 0.071563 | 0.407895 | -1 | 0.192788 |
| D3Z3B8 | | Dlg1 | 40 | S | 1.1 | 0.643993 | 0.927536 | -1 | 0.855532 |
| Q91XM9 | Disks large homolo | Dlg2 | 414 | S | 1 | 0.890627 | 0.99118 | 1.4 | 0.060501 |
| Q9D415 | Disks large-associa | Dlgap1 | 509 | S | -2.4 | 0.180565 | 0.632353 | -1 | 0.799105 |
| Q9D415 | Disks large-associa | Dlgap1 | 973 | S | 3.4 | 0.045478 | 0.366412 | 2.9 | 0.175126 |
| Q9D415 | Disks large-associa | Dlgap1 | 437 | S | 1.7 | 0.238872 | 0.696429 | 3 | 0.003474 |
| Q8BJ42 | Disks large-associa | Dlgap2 | 1012 | S | -1.1 | 0.598349 | 0.92272 | 1.7 | 0.967319 |
| Q8BJ42 | Disks large-associa | Dlgap2 | 745 | S | -1.9 | 0.048443 | 0.366412 | -1 | 0.739638 |
| Q6PFD5 | Disks large-associa | Dlgap3 | 712 | S | -1 | 0.887292 | 0.99118 | 1.1 | 0.713391 |
| Q6PFD5 | Disks large-associa | Dlgap3 | 362 | T | -1.1 | 0.597941 | 0.92272 | -1 | 0.902654 |
| Q6PFD5 | Disks large-associa | Dlgap3 | 363 | Y | -1 | 0.780741 | 0.963975 | -1 | 0.902654 |
| Q6PFD5 | Disks large-associa | Dlgap3 | 365 | Y | -1.1 | 0.532019 | 0.896435 | -1 | 0.902654 |
| Q6PFD5 | Disks large-associa | Dlgap3 | 362 | T | -2.6 | 0.246941 | 0.699708 | -1 | 0.581576 |
| Q6PFD5 | Disks large-associa | Dlgap3 | 363 | Y | -2.3 | 0.576727 | 0.912975 | 1 | 0.465544 |
| Q6PFD5 | Disks large-associa | Dlgap3 | 365 | Y | -2.6 | 0.325368 | 0.766265 | 1 | 0.492031 |
| B1AZP2 | Disks large-associa | Dlgap4 | 405 | S | -1.4 | 0.128648 | 0.530973 | -2 | 0.00777 |
| Q9WV69 | Dematin | Dmtn | 226 | S | -1.8 | 0.04482 | 0.366412 | -1 | 0.078675 |
| Q9WV69 | Dematin | Dmtn | 333 | S | -1.4 | 0.229835 | 0.68 | -1 | 0.82176 |

| | | | | | | | | | |
|---|---|---|---|---|---|---|---|---|---|
| Q9WV69 | Dematin | Dmtn | 96 | S | -1.7 | 0.206866 | 0.65894 | 1.2 | 0.538254 |
| Q8BPN8 | DmX-like protein 2 | Dmxl2 | 1288 | S | 1.5 | 0.616669 | 0.926975 | 1.3 | 0.289408 |
| Q8BPN8 | DmX-like protein 2 | Dmxl2 | 1856 | S | 2.9 | 0.452339 | 0.859073 | 1.2 | 0.410467 |
| Q8BPN8 | DmX-like protein 2 | Dmxl2 | 423 | S | -1.6 | 0.161033 | 0.590551 | -1 | 0.245483 |
| P60904 | DnaJ homolog subf | Dnajc5 | 10 | S | 3.1 | 0.137265 | 0.547414 | 8.8 | 0.011494 |
| Q80TZ3 | Putative tyrosine-pr | Dnajc6 | 35 | S | 3 | 0.133859 | 0.532751 | 4.1 | 0.00121 |
| Q80TZ3 | Putative tyrosine-pr | Dnajc6 | 36 | S | 2.8 | 0.205037 | 0.65894 | 3.9 | 0.001458 |
| P39053-4 | Dynamin-1 | Dnm1 | 851 | S | -1 | 0.887073 | 0.99118 | 1.3 | 0.693063 |
| Q8K1M6 | Dynamin-1-like prot | Dnm1l | 622 | S | 1.4 | 0.153837 | 0.571429 | 1.1 | 0.712757 |
| Q8BZ98 | Dynamin-3 | Dnm3 | 763 | S | 1.4 | 0.299508 | 0.742347 | 1.3 | 0.252608 |
| O08553 | Dihydropyrimidinase | Dpysl2 | 540 | S | 2.8 | 0.072189 | 0.407895 | 1.9 | 0.355403 |
| O08553 | Dihydropyrimidinase | Dpysl2 | 542 | S | 9.5 | 0.001187 | 0.090909 | 6.7 | 0.006138 |
| O08553 | Dihydropyrimidinase | Dpysl2 | 522 | S | 2.7 | 0.337952 | 0.787234 | 3.7 | 0.014559 |
| O08553 | Dihydropyrimidinase | Dpysl2 | 509 | T | 1.3 | 0.569741 | 0.912975 | 1.9 | 0.02531 |
| O08553 | Dihydropyrimidinase | Dpysl2 | 521 | T | 2.7 | 0.337952 | 0.787234 | 3.7 | 0.014559 |
| O08553 | Dihydropyrimidinase | Dpysl2 | 522 | S | -1.4 | 0.800283 | 0.967352 | -2 | 0.646032 |
| O08553 | Dihydropyrimidinase | Dpysl2 | 518 | S | -1.4 | 0.163899 | 0.592157 | -2 | 0.017042 |
| O08553 | Dihydropyrimidinase | Dpysl2 | 521 | T | -1.4 | 0.63898 | 0.927536 | -2 | 0.479234 |
| O08553 | Dihydropyrimidinase | Dpysl2 | 522 | S | -1.2 | 0.727053 | 0.963975 | -2 | 0.954432 |
| O08553 | Dihydropyrimidinase | Dpysl2 | 517 | S | 2 | 0.854386 | 0.988571 | 1.7 | 0.199604 |
| O08553 | Dihydropyrimidinase | Dpysl2 | 518 | S | 1.6 | 0.583402 | 0.916404 | 1.4 | 0.200532 |
| O08553 | Dihydropyrimidinase | Dpysl2 | 521 | T | -1.3 | 0.946709 | 0.993884 | -2 | 0.929927 |
| O08553 | Dihydropyrimidinase | Dpysl2 | 514 | T | -2 | 0.086531 | 0.44382 | -3 | 0.039494 |
| Q62188 | Dihydropyrimidinase | Dpysl3 | 522 | S | -2.1 | 0.028138 | 0.306667 | -1 | 0.892767 |
| Q62188 | Dihydropyrimidinase | Dpysl3 | 509 | T | 1.8 | 0.049129 | 0.366412 | 2.4 | 0.007387 |
| Q62188 | Dihydropyrimidinase | Dpysl3 | 522 | S | -2 | 0.055806 | 0.366412 | -3 | 0.012496 |
| Q62188 | Dihydropyrimidinase | Dpysl3 | 514 | T | -2 | 0.053481 | 0.366412 | -2 | 0.038709 |
| Q62188 | Dihydropyrimidinase | Dpysl3 | 508 | T | -1.5 | 0.116346 | 0.5 | -2 | 0.006214 |
| Q62188 | Dihydropyrimidinase | Dpysl3 | 509 | T | -1.5 | 0.110856 | 0.495238 | -2 | 0.006272 |
| Q9R0P5 | Destrin | Dstn | 3 | S | -4.7 | 0.001007 | 0.090909 | -6 | 0.001148 |
| O88485 | Cytoplasmic dynein | Dync1i1 | 618 | S | 1.5 | 0.701866 | 0.963975 | 2.6 | 0.078551 |
| O88485 | Cytoplasmic dynein | Dync1i1 | 162 | S | -1.5 | 0.482894 | 0.874773 | -9 | 0.116259 |
| O88485 | Cytoplasmic dynein | Dync1i1 | 159 | T | 1.2 | 0.529883 | 0.894378 | 2.6 | 0.26226 |
| Q8R1Q8 | Cytoplasmic dynein | Dync1li1 | 207 | S | 1.4 | 0.319252 | 0.753027 | 2.8 | 0.050382 |
| Q8R1Q8 | Cytoplasmic dynein | Dync1li1 | 516 | S | -2.5 | 0.445006 | 0.853516 | 1.3 | 0.306888 |
| Q8R1Q8 | Cytoplasmic dynein | Dync1li1 | 510 | S | -1.7 | 0.036062 | 0.329787 | -2 | 0.002341 |
| Q8R1Q8 | Cytoplasmic dynein | Dync1li1 | 516 | S | -1.1 | 0.621783 | 0.926975 | -4 | 0.361869 |
| Q6PDL0 | Cytoplasmic dynein | Dync1li2 | 194 | S | -1.1 | 0.480308 | 0.874773 | -1 | 0.535489 |
| P57776 | Elongation factor 1- | Eef1d | 133 | S | 1.8 | 0.008409 | 0.195652 | 1.6 | 0.003447 |
| P57776 | Elongation factor 1- | Eef1d | 162 | S | -2 | 0.150167 | 0.571429 | 2 | 0.0855 |
| Q8CHW4 | Translation initiatior | Eif2b5 | 540 | S | -1 | 0.859826 | 0.988571 | -1 | 0.918937 |
| Q99L45 | Eukaryotic translati | Eif2s2 | 2 | S | -1 | 0.896537 | 0.99118 | -1 | 0.265804 |
| Q8JZQ9 | Eukaryotic translati | Eif3b | 75 | S | 1.1 | 0.980106 | 0.993884 | -1 | 0.654941 |
| Q8JZQ9 | Eukaryotic translati | Eif3b | 75 | S | -1.5 | 0.044273 | 0.366412 | -2 | 0.001064 |
| Q8JZQ9 | Eukaryotic translati | Eif3b | 79 | S | -1.9 | 0.080456 | 0.427711 | -3 | 0.036605 |
| Q9Z1D1 | Eukaryotic translati | Eif3g | 42 | S | 1.2 | 0.693101 | 0.960894 | -2 | 0.280358 |
| Q8BGD9 | Eukaryotic translati | Eif4b | 498 | S | 1.6 | 0.279756 | 0.716535 | 1.1 | 0.527185 |
| Q80XI3 | Eukaryotic translati | Eif4g3 | 267 | S | 1.3 | 0.055365 | 0.366412 | -1 | 0.206934 |
| Q05D44 | Eukaryotic translati | Eif5b | 215 | S | -1 | 0.972723 | 0.993884 | -1 | 0.200079 |
| Q61701 | ELAV-like protein 4 | Elavl4 | 38 | S | 1.2 | 0.587107 | 0.919937 | -1 | 0.790252 |
| P17182 | Alpha-enolase | Eno1 | 263 | S | -1 | 0.786494 | 0.965261 | 1.2 | 0.070694 |
| A2AUK5 | | Epb4.1l1 | 578 | S | -1.4 | 0.274273 | 0.716535 | -2 | 0.05482 |
| A2AUK5 | | Epb4.1l1 | 648 | S | 1.1 | 0.964384 | 0.993884 | -1 | 0.843493 |
| A2AUK5 | Band 4.1-like protei | Epb4.1l1 | 677 | S | -1.4 | 0.383411 | 0.828326 | -2 | 0.054844 |
| A2AUK5 | Band 4.1-like protei | Epb4.1l1 | 868 | S | -1.1 | 0.989117 | 0.995939 | -1 | 0.857609 |
| A2AUK5 | Band 4.1-like protei | Epb4.1l1 | 546 | S | 1.2 | 0.969128 | 0.993884 | 1.3 | 0.483756 |
| A2AUK5 | Band 4.1-like protei | Epb4.1l1 | 546 | S | -1 | 0.735679 | 0.963975 | -2 | 0.022696 |
| A2AUK5 | Band 4.1-like protei | Epb4.1l1 | 550 | T | -1 | 0.735679 | 0.963975 | -2 | 0.022696 |
| A0A286YD | Band 4.1-like protei | Epb4.1l3 | 268 | S | 1.7 | 0.06371 | 0.381944 | -1 | 0.159844 |

| | | | | | | | | | |
|---|---|---|---|---|---|---|---|---|---|
| A0A286YD | Band 4.1-like protei | Epb4.1l3 | 251 | S | 1.2 | 0.663649 | 0.940678 | 1.1 | 0.767877 |
| A0A286YD | Band 4.1-like protei | Epb4.1l3 | 762 | S | 1.4 | 0.518024 | 0.887153 | -1 | 0.409548 |
| A0A286YD | Band 4.1-like protei | Epb4.1l3 | 505 | S | -4 | 0.003193 | 0.105263 | -2 | 0.043368 |
| Q5EBJ4 | Ermin | Ermn | 211 | S | -1.5 | 0.365432 | 0.818386 | 2 | 0.08867 |
| P70429 | Ena/VASP-like prot | Evl | 327 | S | -1.1 | 0.977228 | 0.993884 | 1.1 | 0.722541 |
| P70429 | Ena/VASP-like prot | Evl | 329 | S | 1.4 | 0.279781 | 0.716535 | 2.2 | 0.068339 |
| Q3TY60 | Protein FAM131B | Fam131b | 322 | S | -1.4 | 0.04892 | 0.366412 | -2 | 0.122404 |
| Q8QZR8 | Cyclin-related prote | Fam58b | 107 | S | 1.1 | 0.955185 | 0.993884 | -1 | 0.845593 |
| Q8QZR8 | Cyclin-related prote | Fam58b | 114 | S | -1.2 | 0.532239 | 0.896435 | -1 | 0.548978 |
| Q6NS60 | F-box only protein 4 | Fbxo41 | 477 | T | 1.5 | 0.424299 | 0.848425 | 2.9 | 0.006039 |
| P09528 | Ferritin heavy chain | Fth1 | 5 | S | 5.2 | 0.010301 | 0.195652 | 3 | 0.165365 |
| Q9Z239 | Phospholemman | Fxyd1 | 82 | S | 2 | 0.072307 | 0.407895 | 2.3 | 0.543161 |
| Q9Z239 | Phospholemman | Fxyd1 | 83 | S | 2 | 0.087572 | 0.44382 | 2.4 | 0.385931 |
| P97855 | Ras GTPase-activa | G3bp1 | 231 | S | 1.4 | 0.948006 | 0.993884 | -2 | 0.235836 |
| P97379 | Ras GTPase-activa | G3bp2 | 225 | S | 1.6 | 0.768849 | 0.963975 | 1.6 | 0.710539 |
| P97379 | Ras GTPase-activa | G3bp2 | 227 | T | 1.3 | 0.929517 | 0.993884 | 1.3 | 0.791364 |
| P06837 | Neuromodulin | Gap43 | 142 | S | 2 | 0.028111 | 0.306667 | -1 | 0.743234 |
| P06837 | Neuromodulin | Gap43 | 96 | S | 1.4 | 0.493268 | 0.879061 | -1 | 0.739984 |
| P06837 | Neuromodulin | Gap43 | 95 | T | 31 | 0.079556 | 0.427711 | 31 | 0.28766 |
| P06837 | Neuromodulin | Gap43 | 171 | T | 1 | 0.927673 | 0.993884 | -1 | 0.574803 |
| P06837 | Neuromodulin | Gap43 | 172 | T | 1.5 | 0.301527 | 0.743655 | 1.6 | 0.037044 |
| Q68FF6 | ARF GTPase-activa | Git1 | 601 | S | 1.6 | 0.625252 | 0.927536 | 1.9 | 0.360655 |
| Q68FF6 | ARF GTPase-activa | Git1 | 370 | S | 3.6 | 0.171324 | 0.612167 | 3.8 | 0.167876 |
| Q68FF6 | ARF GTPase-activa | Git1 | 371 | S | -49 | 0.039928 | 0.350515 | -2 | 0.533153 |
| Q68FF6 | ARF GTPase-activa | Git1 | 397 | S | -1.1 | 0.619927 | 0.926975 | 1.8 | 0.237653 |
| P23242 | Gap junction alpha- | Gja1 | 325 | S | 3.6 | 0.166236 | 0.6 | 2.9 | 0.122979 |
| A2AJA9 | Uncharacterized pro | Gm996 | 109 | S | 1.6 | 0.270991 | 0.716535 | 1.6 | 0.280249 |
| Q8BUV3 | Gephyrin;Molybdop | Gphn | 270 | S | -1.3 | 0.200573 | 0.658621 | -2 | 0.019951 |
| Q8BUV3 | Gephyrin;Molybdop | Gphn | 188 | S | 1.1 | 0.514183 | 0.884817 | 1.7 | 0.270475 |
| Q8BUV3 | Gephyrin;Molybdop | Gphn | 194 | S | 1 | 0.586593 | 0.919937 | 1.6 | 0.306739 |
| P35802 | Neuronal membran | Gpm6a | 256 | S | 1.7 | 0.214229 | 0.67101 | 6.3 | 0.039162 |
| P35802 | Neuronal membran | Gpm6a | 267 | S | -1.3 | 0.912104 | 0.992333 | 1.9 | 0.49009 |
| P35802 | Neuronal membran | Gpm6a | 268 | T | 2.4 | 0.246567 | 0.699708 | 5.7 | 0.185911 |
| P35803 | Neuronal membran | Gpm6b | 318 | S | 1 | 0.868348 | 0.988571 | 1.1 | 0.766437 |
| P35803 | Neuronal membran | Gpm6b | 320 | S | -1.2 | 0.69804 | 0.963975 | -3 | 0.055714 |
| Q3UNH4 | G protein-regulated | Gprin1 | 693 | S | 1.7 | 0.307175 | 0.749373 | 2.1 | 0.185811 |
| Q3UNH4 | G protein-regulated | Gprin1 | 219 | S | 1.2 | 0.482612 | 0.874773 | 1.1 | 0.686922 |
| Q3UNH4 | G protein-regulated | Gprin1 | 575 | T | 1.1 | 0.901263 | 0.99118 | 1.1 | 0.22376 |
| Q8BWS5 | G protein-regulated | Gprin3 | 325 | S | -1.3 | 0.239492 | 0.696429 | -1 | 0.087235 |
| Q8BWS5 | G protein-regulated | Gprin3 | 327 | S | -1.2 | 0.35904 | 0.809524 | -1 | 0.129232 |
| Q6IR34 | G-protein-signaling | Gpsm1 | 490 | S | 1.1 | 0.940902 | 0.993884 | 1.3 | 0.832645 |
| Q6IR34 | G-protein-signaling | Gpsm1 | 653 | S | 1 | 0.794708 | 0.966952 | 1.2 | 0.559756 |
| Q8VD04 | GRIP1-associated p | Gripap1 | 657 | S | -1 | 0.933657 | 0.993884 | 14 | 0.014776 |
| Q9WV60 | Glycogen synthase | Gsk3b | 215 | S | 1 | 0.765432 | 0.963975 | -10 | 0.225873 |
| Q9WV60 | Glycogen synthase | Gsk3b | 389 | S | -1.1 | 0.735963 | 0.963975 | -1 | 0.456659 |
| Q9WV60 | Glycogen synthase | Gsk3b | 216 | Y | -1.2 | 0.910273 | 0.992333 | 1.7 | 0.247587 |
| O08582 | GTP-binding protein | Gtpbp1 | 24 | S | -1.4 | 0.437931 | 0.848425 | -6 | 0.009772 |
| O88703 | Potassium/sodium h | Hcn2 | 726 | S | 1.7 | 0.052151 | 0.366412 | 1.3 | 0.247738 |
| P51859 | Hepatoma-derived g | Hdgf | 165 | S | 1.4 | 0.552106 | 0.907285 | 2.2 | 0.001329 |
| Q3UMU9 | Hepatoma-derived g | Hdgfrp2 | 366 | S | -1.8 | 0.236889 | 0.691843 | -2 | 0.112672 |
| Q3UMU9 | Hepatoma-derived g | Hdgfrp2 | 367 | S | -1.5 | 0.501563 | 0.880357 | -1 | 0.551105 |
| Q640R3 | Hepatocyte cell adh | Hepacam | 280 | S | 1.1 | 0.647864 | 0.929293 | -1 | 0.689367 |
| Q640R3 | Hepatocyte cell adh | Hepacam | 379 | S | 1.1 | 0.708536 | 0.963975 | -3 | 0.553848 |
| Q640R3 | Hepatocyte cell adh | Hepacam | 385 | S | 1.4 | 0.244373 | 0.699115 | -1 | 0.89152 |
| Q640R3 | Hepatocyte cell adh | Hepacam | 386 | S | -3.7 | 0.302044 | 0.746835 | -2 | 0.836876 |
| Q640R3 | Hepatocyte cell adh | Hepacam | 378 | T | -1.2 | 0.32795 | 0.768496 | -1 | 0.458096 |
| Q6PAV2 | Probable E3 ubiquit | Herc4 | 318 | S | 3.7 | 0.523886 | 0.894378 | 3.8 | 0.65466 |
| Q6PAV2 | Probable E3 ubiquit | Herc4 | 333 | S | 2.1 | 0.240872 | 0.696429 | -3 | 0.085032 |
| Q6PAV2 | Probable E3 ubiquit | Herc4 | 332 | T | 2.1 | 0.240872 | 0.696429 | -3 | 0.083669 |

| | | | | | | | | | |
|---|---|---|---|---|---|---|---|---|---|
| Q6PAV2 | Probable E3 ubiquit | Herc4 | 317 | Y | -2.1 | 0.512184 | 0.884413 | 1.2 | 0.535651 |
| D3YXG0 | | Hmcn1 | 4090 | T | -1.1 | 0.79198 | 0.966952 | -2 | 0.904461 |
| D3YXG0 | | Hmcn1 | 4087 | Y | -1.1 | 0.860152 | 0.988571 | -2 | 0.863479 |
| P49312 | Heterogeneous nuc | Hnrnpa1 | 4 | S | 2.3 | 0.102728 | 0.475 | -9 | 0.098 |
| P49312 | Heterogeneous nuc | Hnrnpa1 | 6 | S | 1.2 | 0.893849 | 0.99118 | -1 | 0.820586 |
| P49312 | Heterogeneous nuc | Hnrnpa1 | 4 | S | 1.3 | 0.473331 | 0.874773 | 1.5 | 0.194574 |
| P49312 | Heterogeneous nuc | Hnrnpa1 | 6 | S | -1.9 | 0.234965 | 0.690909 | -2 | 0.713365 |
| P49312 | Heterogeneous nuc | Hnrnpa1 | 2 | S | -3.8 | 0.3153 | 0.753027 | -7 | 0.110254 |
| Q9Z204 | Heterogeneous nuc | Hnrnpc | 241 | S | 1.2 | 0.502819 | 0.883186 | 1 | 0.813357 |
| Q60668 | Heterogeneous nuc | Hnrnpd | 83 | S | -6.8 | 0.471963 | 0.874773 | -12 | 0.197395 |
| P70333 | Heterogeneous nuc | Hnrnph2 | 104 | S | 1.1 | 0.619079 | 0.926975 | 1.1 | 0.519848 |
| P61979 | Heterogeneous nuc | Hnrnpk | 284 | S | 1.1 | 0.953134 | 0.993884 | -2 | 0.039648 |
| Q8VEK3 | Heterogeneous nuc | Hnrnpu | 4 | S | 1.9 | 0.604932 | 0.92638 | 1.9 | 0.464816 |
| Q00PI9 | Heterogeneous nuc | Hnrnpul2 | 159 | S | -1.1 | 0.783877 | 0.963975 | -2 | 0.081698 |
| P07901 | Heat shock protein | Hsp90aa1 | 263 | S | -2.3 | 0.013992 | 0.211538 | -1 | 0.080279 |
| P11499 | Heat shock protein | Hsp90ab1 | 255 | S | -1.4 | 0.211865 | 0.664474 | -3 | 0.001445 |
| P11499 | Heat shock protein | Hsp90ab1 | 255 | S | -1.5 | 0.855427 | 0.988571 | -2 | 0.857027 |
| P11499 | Heat shock protein | Hsp90ab1 | 261 | S | -1.5 | 0.9551 | 0.993884 | -2 | 0.855739 |
| Q8K0U4 | Heat shock 70 kDa | Hspa12a | 24 | S | 1.3 | 0.973283 | 0.993884 | -1 | 0.730229 |
| P48722 | Heat shock 70 kDa | Hspa4l | 74 | S | -1.4 | 0.097001 | 0.470588 | -1 | 0.102065 |
| Q61699 | Heat shock protein | Hsph1 | 810 | S | 1.9 | 0.103595 | 0.477612 | 1.8 | 0.143257 |
| Q8R0S2 | IQ motif and SEC7 | Iqsec1 | 179 | S | -1.1 | 0.773397 | 0.963975 | -1 | 0.071627 |
| Q8R0S2 | IQ motif and SEC7 | Iqsec1 | 513 | S | 1.1 | 0.971079 | 0.993884 | 1.2 | 0.268042 |
| Q5DU25 | IQ motif and SEC7 | Iqsec2 | 383 | S | 1.6 | 0.020041 | 0.25 | 1.7 | 0.002969 |
| Q8BNW9 | Kelch repeat and B | Kbtbd11 | 70 | S | 1.1 | 0.647504 | 0.929293 | -1 | 0.192638 |
| Q8BNW9 | Kelch repeat and B | Kbtbd11 | 320 | S | -1.2 | 0.361194 | 0.810811 | -1 | 0.384219 |
| Q8BNW9 | Kelch repeat and B | Kbtbd11 | 316 | S | 1.1 | 0.787505 | 0.966543 | -3 | 0.238139 |
| Q8BNW9 | Kelch repeat and B | Kbtbd11 | 320 | S | 1.5 | 0.273888 | 0.716535 | -3 | 0.253082 |
| P63141 | Potassium voltage-( | Kcna2 | 434 | S | 1.6 | 0.202875 | 0.65894 | 1.8 | 0.51072 |
| Q6WVG3 | BTB/POZ domain-c | Kctd12 | 187 | S | 1.4 | 0.385599 | 0.828326 | 1.1 | 0.598436 |
| Q6WVG3 | BTB/POZ domain-c | Kctd12 | 189 | S | 1.8 | 0.20379 | 0.65894 | 1.4 | 0.347317 |
| Q3U0V1 | Far upstream eleme | Khsrp | 182 | S | 1.3 | 0.465346 | 0.872137 | 2 | 0.02107 |
| Q8R0A7 | Uncharacterized pro | Kiaa0513 | 275 | S | 1.1 | 0.636026 | 0.927536 | -1 | 0.877552 |
| Q148V7 | LisH domain and HI | Kiaa1468 | 180 | S | -1.2 | 0.776051 | 0.963975 | 1.3 | 0.314817 |
| Q8BRV5 | Uncharacterized pro | Kiaa1671 | 205 | S | -1.5 | 0.099445 | 0.471795 | 1.3 | 0.14655 |
| P33173 | Kinesin-like protein | Kif1a | 937 | S | -2 | 0.021194 | 0.257576 | -1 | 0.399866 |
| Q9QXL2 | Kinesin-like protein | Kif21a | 855 | S | 1.1 | 0.788614 | 0.966952 | 2.8 | 0.061268 |
| Q9QXL2 | Kinesin-like protein | Kif21a | 257 | S | 1.4 | 0.504568 | 0.883186 | 1.8 | 0.073844 |
| Q91YS4 | | Klc2 | 151 | S | -2.6 | 0.008993 | 0.195652 | -2 | 0.107073 |
| O35344 | Importin subunit alp | Kpna3 | 56 | S | -1.3 | 0.193036 | 0.644366 | 1.1 | 0.552752 |
| Q61097 | Kinase suppressor ( | Ksr1 | 392 | S | 1.9 | 0.082848 | 0.429412 | 1.6 | 0.50181 |
| P11627 | Neural cell adhesio | L1cam | 1184 | S | -1.5 | 0.386113 | 0.828326 | -1 | 0.454884 |
| Q8VED9 | Galectin-related pro | Lgalsl | 4 | S | -1.8 | 0.148787 | 0.571429 | -2 | 0.082926 |
| Q8VED9 | Galectin-related pro | Lgalsl | 25 | S | 3.3 | 0.032965 | 0.311111 | 2.2 | 0.351835 |
| Q9D1T0 | Leucine-rich repeat | Lingo1 | 596 | S | 1 | 0.853707 | 0.988571 | -1 | 0.869331 |
| P21619 | Lamin-B2 | Lmnb2 | 427 | S | 1.6 | 0.343245 | 0.787383 | 1.5 | 0.362103 |
| Q7TQ95 | Protein lunapark | Lnp | 411 | S | 1.2 | 0.078479 | 0.427711 | 1.5 | 0.003585 |
| Q7TME0 | Lipid phosphate pho | Lppr4 | 549 | S | 1.3 | 0.746732 | 0.963975 | 1.2 | 0.392036 |
| P0C192 | Leucine-rich repeat- | Lrrc4b | 698 | S | 1 | 0.756422 | 0.963975 | -1 | 0.641345 |
| P0C192 | Leucine-rich repeat- | Lrrc4b | 700 | S | 1 | 0.756422 | 0.963975 | -1 | 0.641345 |
| Q80TE7 | Leucine-rich repeat- | Lrrc7 | 1392 | S | -1.1 | 0.556818 | 0.912975 | 1 | 0.817454 |
| Q80TE7 | Leucine-rich repeat- | Lrrc7 | 949 | S | -1.4 | 0.195916 | 0.648084 | -2 | 0.083825 |
| Q80U28 | MAP kinase-activati | Madd | 1058 | S | 1.4 | 0.135308 | 0.534783 | 1.3 | 0.163382 |
| Q9QYR6 | Microtubule-associa | Map1a | 667 | S | 2.1 | 0.479861 | 0.874773 | 5.2 | 0.13613 |
| Q9QYR6 | Microtubule-associa | Map1a | 1606 | S | -2.7 | 0.128101 | 0.530973 | -3 | 0.154671 |
| Q9QYR6 | Microtubule-associa | Map1a | 981 | S | -1.5 | 0.266027 | 0.713499 | -1 | 0.384126 |
| Q9QYR6 | Microtubule-associa | Map1a | 499 | S | 1.3 | 0.61976 | 0.926975 | 1.2 | 0.95884 |
| Q9QYR6 | Microtubule-associa | Map1a | 1580 | S | -1.1 | 0.804544 | 0.967352 | 1.7 | 0.049643 |
| Q9QYR6 | Microtubule-associa | Map1a | 1634 | S | -2 | 0.40122 | 0.839662 | -1 | 0.635313 |

| | | | | | | | | | |
|---|---|---|---|---|---|---|---|---|---|
| Q9QYR6 | Microtubule-associa | Map1a | 1648 | S | 1.4 | 0.722981 | 0.963975 | 1.1 | 0.686532 |
| Q9QYR6 | Microtubule-associa | Map1a | 991 | S | 1.4 | 0.538531 | 0.900673 | 1.9 | 0.191658 |
| Q9QYR6 | Microtubule-associa | Map1a | 2230 | S | 2.4 | 0.413605 | 0.848425 | 2 | 0.263613 |
| Q9QYR6 | Microtubule-associa | Map1a | 1205 | S | 1.3 | 0.651347 | 0.935252 | 2 | 0.699253 |
| Q9QYR6 | Microtubule-associa | Map1a | 2082 | S | 3.2 | 0.056024 | 0.366412 | 1.9 | 0.376626 |
| Q9QYR6 | Microtubule-associa | Map1a | 1768 | S | 1.2 | 0.482723 | 0.874773 | 1.3 | 0.279359 |
| Q9QYR6 | Microtubule-associa | Map1a | 1062 | S | 5.3 | 0.25768 | 0.70904 | 3.1 | 0.239584 |
| Q9QYR6 | Microtubule-associa | Map1a | 2603 | S | 1.8 | 0.166249 | 0.6 | -1 | 0.348655 |
| Q9QYR6 | Microtubule-associa | Map1a | 873 | S | 1.3 | 0.322554 | 0.756039 | 1.1 | 0.904202 |
| Q9QYR6 | Microtubule-associa | Map1a | 114 | S | -1.7 | 0.251928 | 0.704871 | -1 | 0.624171 |
| Q9QYR6 | Microtubule-associa | Map1a | 1796 | S | -1.3 | 0.665198 | 0.940678 | 1.3 | 0.518738 |
| Q9QYR6 | Microtubule-associa | Map1a | 1575 | T | 7.2 | 0.020004 | 0.25 | 2.6 | 0.147987 |
| Q9QYR6 | Microtubule-associa | Map1a | 319 | S | -2 | 0.699341 | 0.963975 | -3 | 0.26707 |
| Q9QYR6 | Microtubule-associa | Map1a | 322 | S | -2 | 0.729316 | 0.963975 | -3 | 0.354481 |
| Q9QYR6 | Microtubule-associa | Map1a | 1648 | S | -1.3 | 0.963742 | 0.993884 | -2 | 0.437157 |
| Q9QYR6 | Microtubule-associa | Map1a | 991 | S | -1.6 | 0.179103 | 0.62963 | -2 | 0.097363 |
| Q9QYR6 | Microtubule-associa | Map1a | 1008 | S | -1.2 | 0.960097 | 0.993884 | -1 | 0.595233 |
| Q9QYR6 | Microtubule-associa | Map1a | 2234 | S | -1.1 | 0.60577 | 0.926493 | 1.1 | 0.394443 |
| Q9QYR6 | Microtubule-associa | Map1a | 1768 | S | -1.5 | 0.151679 | 0.571429 | -1 | 0.725923 |
| Q9QYR6 | Microtubule-associa | Map1a | 1772 | S | -1.5 | 0.20495 | 0.65894 | -1 | 0.725923 |
| Q9QYR6 | Microtubule-associa | Map1a | 895 | S | 1.3 | 0.438723 | 0.848425 | 2.6 | 0.067416 |
| Q9QYR6 | Microtubule-associa | Map1a | 1789 | S | -2 | 0.012769 | 0.208333 | -3 | 0.034219 |
| P14873 | Microtubule-associa | Map1b | 1371 | S | 1.1 | 0.971247 | 0.993884 | -2 | 0.13199 |
| P14873 | Microtubule-associa | Map1b | 1373 | S | 1.1 | 0.971247 | 0.993884 | -2 | 0.13199 |
| P14873 | Microtubule-associa | Map1b | 1384 | S | -2 | 0.186765 | 0.638989 | -2 | 0.223676 |
| P14873 | Microtubule-associa | Map1b | 1151 | S | -1.4 | 0.845433 | 0.988571 | -3 | 0.282196 |
| P14873 | Microtubule-associa | Map1b | 614 | S | -1.2 | 0.596259 | 0.92272 | -2 | 0.044025 |
| P14873 | Microtubule-associa | Map1b | 1813 | S | 7.7 | 0.221565 | 0.674923 | 14 | 0.208672 |
| P14873 | Microtubule-associa | Map1b | 561 | S | 1 | 0.948528 | 0.993884 | -1 | 0.886281 |
| P14873 | Microtubule-associa | Map1b | 1497 | S | 2.9 | 0.261583 | 0.70904 | 2.1 | 0.156933 |
| P14873 | Microtubule-associa | Map1b | 1438 | S | 2 | 0.397747 | 0.836864 | 2.5 | 0.062501 |
| P14873 | Microtubule-associa | Map1b | 1781 | S | -1.2 | 0.771178 | 0.963975 | -1 | 0.338643 |
| P14873 | Microtubule-associa | Map1b | 2030 | S | -1.2 | 0.750554 | 0.963975 | -1 | 0.817467 |
| P14873 | Microtubule-associa | Map1b | 1395 | S | 1.5 | 0.486883 | 0.874773 | -1 | 0.559192 |
| P14873 | Microtubule-associa | Map1b | 1260 | S | -1.1 | 0.577355 | 0.912975 | -1 | 0.187398 |
| P14873 | Microtubule-associa | Map1b | 1307 | S | -1.6 | 0.159626 | 0.589641 | -1 | 0.902271 |
| P14873 | Microtubule-associa | Map1b | 1317 | S | -1.5 | 0.405581 | 0.844538 | 1.7 | 0.209808 |
| P14873 | Microtubule-associa | Map1b | 1877 | S | -1.3 | 0.525771 | 0.894378 | -1 | 0.22054 |
| P14873 | Microtubule-associa | Map1b | 1775 | S | -1.2 | 0.436874 | 0.848425 | -1 | 0.770779 |
| P14873 | Microtubule-associa | Map1b | 1373 | S | -2 | 0.374137 | 0.823789 | -2 | 0.709088 |
| P14873 | Microtubule-associa | Map1b | 1384 | S | -2.1 | 0.398603 | 0.836864 | -2 | 0.714014 |
| P14873 | Microtubule-associa | Map1b | 1789 | S | -1.4 | 0.44326 | 0.853516 | -1 | 0.829576 |
| P14873 | Microtubule-associa | Map1b | 1793 | S | -2 | 0.076473 | 0.421384 | -2 | 0.333573 |
| P14873 | Microtubule-associa | Map1b | 1255 | S | -1.4 | 0.850565 | 0.988571 | -11 | 0.013604 |
| P14873 | Microtubule-associa | Map1b | 1260 | S | -1.8 | 0.073619 | 0.410256 | -2 | 0.008548 |
| P14873 | Microtubule-associa | Map1b | 1319 | S | -4.2 | 0.481957 | 0.874773 | -2 | 0.837494 |
| P14873 | Microtubule-associa | Map1b | 1325 | S | -4 | 0.538768 | 0.900673 | -2 | 0.853674 |
| P20357 | Microtubule-associa | Map2 | 1783 | S | -1.5 | 0.149061 | 0.571429 | -2 | 0.012564 |
| P20357 | Microtubule-associa | Map2 | 1352 | S | -1.2 | 0.479568 | 0.874773 | -2 | 0.185671 |
| P20357 | Microtubule-associa | Map2 | 1537 | S | 1 | 0.71572 | 0.963975 | -1 | 0.614283 |
| P20357 | Microtubule-associa | Map2 | 1539 | S | -1.9 | 0.298929 | 0.742347 | 1.5 | 0.284317 |
| P20357 | Microtubule-associa | Map2 | 938 | S | -1.4 | 0.228897 | 0.68 | -3 | 0.00569 |
| P20357 | Microtubule-associa | Map2 | 1013 | S | -1.1 | 0.958922 | 0.993884 | 2.6 | 0.036751 |
| P20357 | Microtubule-associa | Map2 | 1161 | S | -2.4 | 0.50562 | 0.883392 | -1 | 0.770308 |
| P20357 | Microtubule-associa | Map2 | 739 | S | 2.6 | 0.736012 | 0.963975 | -3 | 0.518998 |
| P20357 | Microtubule-associa | Map2 | 608 | S | -1.1 | 0.570561 | 0.912975 | -1 | 0.605425 |
| P20357 | Microtubule-associa | Map2 | 1485 | S | 1.5 | 0.47426 | 0.874773 | 1.1 | 0.945746 |
| P20357 | Microtubule-associa | Map2 | 1609 | T | -1.2 | 0.461092 | 0.868069 | -5 | 0.009669 |
| P20357 | Microtubule-associa | Map2 | 626 | S | 4 | 0.383018 | 0.828326 | 1.1 | 0.955009 |

| | | | | | | | | | |
|---|---|---|---|---|---|---|---|---|---|
| P20357 | Microtubule-associa | Map2 | 654 | S | 4.2 | 0.426158 | 0.848425 | 1.1 | 0.868733 |
| P20357 | Microtubule-associa | Map2 | 1352 | S | -3.6 | 0.002422 | 0.105263 | -2 | 0.020249 |
| P20357 | Microtubule-associa | Map2 | 1358 | T | -3.6 | 0.002422 | 0.105263 | -2 | 0.020249 |
| P20357 | Microtubule-associa | Map2 | 1609 | T | -3.7 | 0.010151 | 0.195652 | -2 | 0.050562 |
| P27546 | Microtubule-associa | Map4 | 667 | S | 1.2 | 0.496584 | 0.880143 | 1.8 | 0.006615 |
| P27546 | Microtubule-associa | Map4 | 667 | S | 1.1 | 0.946537 | 0.993884 | -1 | 0.113469 |
| P27546 | Microtubule-associa | Map4 | 658 | T | 1.5 | 0.321632 | 0.753027 | -1 | 0.809482 |
| Q7TSJ2 | Microtubule-associa | Map6 | 687 | S | 1.8 | 0.123205 | 0.520737 | 2.5 | 0.154721 |
| Q7TSJ2 | Microtubule-associa | Map6 | 632 | S | 1.2 | 0.641951 | 0.927536 | 1 | 0.853465 |
| Q7TSJ2 | Microtubule-associa | Map6 | 905 | S | 1.5 | 0.825671 | 0.980792 | 4.6 | 0.437688 |
| A2AJI0 | MAP7 domain-conta | Map7d1 | 115 | S | -1.7 | 0.041402 | 0.356436 | -2 | 0.008813 |
| A2AJI0 | MAP7 domain-conta | Map7d1 | 118 | S | -1.2 | 0.940098 | 0.993884 | -2 | 0.534018 |
| Q91Y86 | Mitogen-activated p | Mapk8 | 185 | Y | -1.1 | 0.831747 | 0.986874 | 1.1 | 0.653203 |
| P10637 | Microtubule-associa | Mapt | 696 | S | -1.1 | 0.207567 | 0.65894 | -3 | 0.019312 |
| P10637 | Microtubule-associa | Mapt | 648 | S | -1.7 | 0.074022 | 0.410256 | -1 | 0.592681 |
| P10637 | Microtubule-associa | Mapt | 494 | S | -1.6 | 0.111204 | 0.495238 | 1.1 | 0.766825 |
| P10637 | Microtubule-associa | Mapt | 506 | S | 1.6 | 0.386306 | 0.828326 | 1.3 | 0.293413 |
| P10637 | Microtubule-associa | Mapt | 695 | T | 32 | 0.110396 | 0.495238 | 126 | 0.004734 |
| P10637 | Microtubule-associa | Mapt | 473 | T | 1.9 | 0.095507 | 0.47027 | 1.9 | 0.931715 |
| P10637 | Microtubule-associa | Mapt | 688 | S | -1.8 | 0.038118 | 0.34375 | 2 | 0.022097 |
| P10637 | Microtubule-associa | Mapt | 692 | S | -3.4 | 0.004867 | 0.173913 | -2 | 0.003069 |
| P10637 | Microtubule-associa | Mapt | 696 | S | -2.4 | 0.997181 | 0.997982 | -4 | 0.370656 |
| P10637 | Microtubule-associa | Mapt | 695 | T | -1.8 | 0.723611 | 0.963975 | -2 | 0.693779 |
| P10637 | Microtubule-associa | Mapt | 688 | S | -2.6 | 0.256393 | 0.70904 | 1.7 | 0.177932 |
| P10637 | Microtubule-associa | Mapt | 692 | S | -2.8 | 0.273346 | 0.716535 | 1.8 | 0.186225 |
| P10637 | Microtubule-associa | Mapt | 696 | S | -3.1 | 0.18991 | 0.64311 | 1.7 | 0.227807 |
| P10637 | Microtubule-associa | Mapt | 695 | T | -2.4 | 0.395131 | 0.836864 | 1.8 | 0.166528 |
| P26645 | Myristoylated alanin | Marcks | 112 | S | -1.1 | 0.86792 | 0.988571 | -2 | 0.60754 |
| P26645 | Myristoylated alanin | Marcks | 113 | S | -1.1 | 0.978494 | 0.993884 | -2 | 0.604084 |
| P26645 | Myristoylated alanin | Marcks | 122 | S | -2.1 | 0.93619 | 0.993884 | -3 | 0.504233 |
| P26645 | Myristoylated alanin | Marcks | 27 | S | 1 | 0.945321 | 0.993884 | 1.2 | 0.544936 |
| P26645 | Myristoylated alanin | Marcks | 163 | S | 1.9 | 0.141909 | 0.559829 | 1.6 | 0.16645 |
| P26645 | Myristoylated alanin | Marcks | 143 | T | 4.7 | 0.115094 | 0.49763 | 4.1 | 0.193673 |
| P28667 | MARCKS-related pr | Marcksl1 | 22 | S | 1.1 | 0.467992 | 0.874773 | 1 | 0.913526 |
| Q8K310 | Matrin-3 | Matr3 | 195 | S | 1 | 0.93586 | 0.993884 | 3.7 | 0.138274 |
| Q8K310 | Matrin-3 | Matr3 | 598 | S | 1.7 | 0.521988 | 0.894281 | 1.7 | 0.036223 |
| Q8K310 | Matrin-3 | Matr3 | 598 | S | -1.3 | 0.574109 | 0.912975 | -1 | 0.709798 |
| Q8K310 | Matrin-3 | Matr3 | 604 | S | -1.9 | 0.100759 | 0.471795 | -2 | 0.06841 |
| P04370-6 | Myelin basic protein | Mbp | 145 | S | -1.5 | 0.121781 | 0.520737 | -2 | 0.040472 |
| P04370-6;I | Myelin basic protein | Mbp | 66 | S | -1.6 | 0.16915 | 0.605364 | -1 | 0.276252 |
| P04370-6;I | Myelin basic protein | Mbp | 94 | S | 1.8 | 0.534046 | 0.89661 | -1 | 0.834481 |
| P04370-6;I | Myelin basic protein | Mbp | 99 | S | 1.9 | 0.975492 | 0.993884 | -1 | 0.875625 |
| P04370-4 | Myelin basic protein | Mbp | 139 | S | 2.3 | 0.150755 | 0.571429 | 3.2 | 0.009379 |
| P04370-6;I | Myelin basic protein | Mbp | 122 | T | 2.4 | 0.224453 | 0.674923 | 2.8 | 0.001699 |
| P14152 | Malate dehydrogena | Mdh1 | 241 | S | 1.7 | 0.162236 | 0.590551 | 1.6 | 0.01671 |
| Q9Z2D6 | Methyl-CpG-binding | Mecp2 | 80 | S | -1.2 | 0.620563 | 0.926975 | 1.3 | 0.420314 |
| Q6PCP5 | Mitochondrial fissio | Mff | 151 | S | 1.4 | 0.505323 | 0.883186 | 1.1 | 0.609174 |
| Q8CJ19 | Protein-methionine | Mical3 | 1154 | S | 1.1 | 0.710919 | 0.963975 | -1 | 0.594635 |
| Q9JM52 | Misshapen-like kina | Mink1 | 745 | S | 3.1 | 0.469333 | 0.874773 | 1.7 | 0.761973 |
| Q9JM52 | Misshapen-like kina | Mink1 | 746 | S | -1.2 | 0.254053 | 0.704871 | -2 | 0.062109 |
| Q9JM52 | Misshapen-like kina | Mink1 | 729 | S | 1.4 | 0.879925 | 0.99118 | 2.4 | 0.08455 |
| Q9JM52 | Misshapen-like kina | Mink1 | 603 | S | 1.5 | 0.16711 | 0.6 | -2 | 0.214911 |
| Q99KX1 | Myeloid leukemia fa | Mlf2 | 237 | S | -1.2 | 0.25353 | 0.704871 | -1 | 0.823251 |
| Q9D2P8 | Myelin-associated c | Mobp | 85 | S | 2.2 | 0.058155 | 0.381944 | 1.8 | 0.005224 |
| Q9WV34 | MAGUK p55 subfar | Mpp2 | 121 | S | -1.1 | 0.792107 | 0.966952 | 1.2 | 0.875311 |
| Q9D1H8 | 39S ribosomal prote | Mrpl53 | 38 | S | 5.2 | 0.016639 | 0.224138 | 4.4 | 0.157283 |
| Q9D1H8 | 39S ribosomal prote | Mrpl53 | 39 | S | 4.9 | 0.014526 | 0.224138 | 4 | 0.286956 |
| Q9D1H8 | 39S ribosomal prote | Mrpl53 | 32 | T | 4.9 | 0.034788 | 0.32967 | 3.9 | 0.588721 |
| Q9D1H8 | 39S ribosomal prote | Mrpl53 | 36 | T | 4.2 | 0.070042 | 0.407895 | 3.6 | 0.288384 |

| | | | | | | | | | |
|---|---|---|---|---|---|---|---|---|---|
| Q9CWE0 | Mitochondrial fissio | Mtfr1l | 100 | S | -1.1 | 0.853623 | 0.988571 | -1 | 0.531663 |
| Q9Z2C4 | Myotubularin-relate | Mtmr1 | 657 | S | 5.7 | 0.128485 | 0.530973 | 8.8 | 0.048953 |
| D3YTP3 | | Mtx3 | 311 | S | -1.1 | 0.481207 | 0.874773 | -1 | 0.707128 |
| A0A087W! | Mucin-2 | Muc2 | 440 | T | 2.1 | 0.342446 | 0.787383 | 3.9 | 0.137191 |
| A0A087W! | Mucin-2 | Muc2 | 444 | T | 2 | 0.594964 | 0.92272 | 3.8 | 0.237056 |
| Q8VDD5 | Myosin-9 | Myh9 | 1943 | S | -2.3 | 0.053839 | 0.366412 | -1 | 0.223947 |
| Q99104 | Unconventional my | Myo5a | 600 | S | 1 | 0.816945 | 0.974729 | -1 | 0.489516 |
| A0A0R4J1E3 | | NA | 613 | S | 1.6 | 0.471123 | 0.874773 | -2 | 0.164765 |
| A0A286YCB8 | | NA | 126 | S | 2.6 | 0.319094 | 0.753027 | 2 | 0.330611 |
| A0A286YCB8 | | NA | 127 | S | 1.1 | 0.876283 | 0.99118 | -1 | 0.880942 |
| A0A286YCB8 | | NA | 131 | S | -1 | 0.743111 | 0.963975 | -1 | 0.511058 |
| A0A494BB10 | | NA | 42 | S | 2 | 0.244275 | 0.699115 | 2.2 | 0.02728 |
| Q9CWZ7 | Gamma-soluble NS | Napg | 284 | S | 1.8 | 0.340487 | 0.787383 | 2.2 | 0.254808 |
| Q9EPN1 | Neurobeachin | Nbea | 1519 | S | 1.1 | 0.639617 | 0.927536 | -1 | 0.524614 |
| P13595 | Neural cell adhesio | Ncam1 | 774 | S | -1 | 0.946806 | 0.993884 | 2.7 | 0.000462 |
| P13595 | Neural cell adhesio | Ncam1 | 887 | S | 1.3 | 0.259496 | 0.70904 | 1 | 0.892349 |
| P13595 | Neural cell adhesio | Ncam1 | 1005 | S | 1.2 | 0.599636 | 0.924383 | -1 | 0.639822 |
| P13595 | Neural cell adhesio | Ncam1 | 770 | S | -3.8 | 0.009428 | 0.195652 | 1 | 0.772446 |
| P13595 | Neural cell adhesio | Ncam1 | 774 | S | -3.8 | 0.009207 | 0.195652 | 1 | 0.772446 |
| Q62433 | Protein NDRG1 | Ndrg1 | 330 | S | -1.1 | 0.665044 | 0.940678 | -2 | 0.401023 |
| Q9QYG0 | Protein NDRG2 | Ndrg2 | 332 | S | 1.6 | 0.995781 | 0.99798 | 1.7 | 0.329452 |
| Q9QYG0 | Protein NDRG2 | Ndrg2 | 338 | S | 1.2 | 0.572945 | 0.912975 | 1.1 | 0.817819 |
| Q9QYG0 | Protein NDRG2 | Ndrg2 | 352 | S | 2.4 | 0.264015 | 0.713499 | 1.5 | 0.947332 |
| Q9QYG0 | Protein NDRG2 | Ndrg2 | 355 | S | -1.6 | 0.485552 | 0.874773 | -2 | 0.321267 |
| Q9QYG0 | Protein NDRG2 | Ndrg2 | 348 | T | -1.1 | 0.796471 | 0.966952 | 1 | 0.673758 |
| Q9QYG0 | Protein NDRG2 | Ndrg2 | 328 | S | -2.4 | 0.726598 | 0.963975 | -1 | 0.85404 |
| Q9QYG0 | Protein NDRG2 | Ndrg2 | 332 | S | -1.9 | 0.056114 | 0.366412 | -2 | 0.049966 |
| Q9QYG0 | Protein NDRG2 | Ndrg2 | 335 | S | -4.3 | 0.5661 | 0.912975 | -6 | 0.449146 |
| Q9QYG0 | Protein NDRG2 | Ndrg2 | 338 | S | -1.9 | 0.056114 | 0.366412 | -2 | 0.049966 |
| Q9QYG0 | Protein NDRG2 | Ndrg2 | 353 | S | -1.3 | 0.958619 | 0.993884 | -4 | 0.407937 |
| Q9QYG0 | Protein NDRG2 | Ndrg2 | 330 | T | -1.2 | 0.826056 | 0.980815 | 1.2 | 0.572387 |
| Q9QYG0 | Protein NDRG2 | Ndrg2 | 334 | T | -1.5 | 0.74112 | 0.963975 | -1 | 0.811648 |
| Q9QYG0 | Protein NDRG2 | Ndrg2 | 348 | T | -2.3 | 0.210859 | 0.663366 | -2 | 0.639618 |
| Q9QYF9 | Protein NDRG3 | Ndrg3 | 361 | S | -1.2 | 0.69638 | 0.963975 | 1 | 0.859503 |
| Q8BTG7 | Protein NDRG4 | Ndrg4 | 298 | S | 1.2 | 0.57285 | 0.912975 | 1.2 | 0.938547 |
| P19246 | Neurofilament heav | Nefh | 834 | S | -1.2 | 0.326727 | 0.768496 | 1.4 | 0.243719 |
| P19246 | Neurofilament heav | Nefh | 727 | S | 1.4 | 0.288185 | 0.718346 | 1.4 | 0.087588 |
| P19246 | Neurofilament heav | Nefh | 739 | S | 1.3 | 0.558202 | 0.912975 | 1.2 | 0.246299 |
| P19246 | Neurofilament heav | Nefh | 763 | S | 1.4 | 0.379091 | 0.828326 | 1.6 | 0.177586 |
| P19246 | Neurofilament heav | Nefh | 673 | S | 1.6 | 0.198217 | 0.65625 | 1.7 | 0.035863 |
| P19246 | Neurofilament heav | Nefh | 571 | S | 1.3 | 0.578166 | 0.912975 | 1.3 | 0.232203 |
| P19246 | Neurofilament heav | Nefh | 679 | S | 1.3 | 0.431817 | 0.848425 | 1.4 | 0.121808 |
| P19246 | Neurofilament heav | Nefh | 834 | S | -2.3 | 0.004129 | 0.136364 | -3 | 0.020407 |
| P19246 | Neurofilament heav | Nefh | 783 | S | 1.1 | 0.756679 | 0.963975 | -3 | 0.071338 |
| P19246 | Neurofilament heav | Nefh | 789 | S | 1.1 | 0.729443 | 0.963975 | -2 | 0.096249 |
| P19246 | Neurofilament heav | Nefh | 727 | S | -1.4 | 0.836601 | 0.988571 | -5 | 0.283174 |
| P19246 | Neurofilament heav | Nefh | 739 | S | -1.8 | 0.054574 | 0.366412 | -2 | 0.062734 |
| P19246 | Neurofilament heav | Nefh | 601 | S | -1.2 | 0.866954 | 0.988571 | -2 | 0.3314 |
| P19246 | Neurofilament heav | Nefh | 607 | S | -1.6 | 0.114027 | 0.495238 | -2 | 0.029634 |
| P19246 | Neurofilament heav | Nefh | 613 | S | -1.4 | 0.305514 | 0.748744 | -2 | 0.056764 |
| P19246 | Neurofilament heav | Nefh | 757 | S | -1.6 | 0.102714 | 0.475 | -2 | 0.105636 |
| P19246 | Neurofilament heav | Nefh | 763 | S | -1.6 | 0.102714 | 0.475 | -2 | 0.105636 |
| P19246 | Neurofilament heav | Nefh | 541 | S | -1.2 | 0.475842 | 0.874773 | -2 | 0.095238 |
| P19246 | Neurofilament heav | Nefh | 547 | S | 1.2 | 0.86991 | 0.988571 | -1 | 0.670092 |
| P19246 | Neurofilament heav | Nefh | 685 | S | -1.4 | 0.505403 | 0.883186 | -2 | 0.376365 |
| P19246 | Neurofilament heav | Nefh | 691 | S | -1.4 | 0.583925 | 0.916535 | -2 | 0.471434 |
| P19246 | Neurofilament heav | Nefh | 839 | T | -2.3 | 0.004129 | 0.136364 | -3 | 0.024405 |
| P08551 | Neurofilament light | Nefl | 473 | S | 1.1 | 0.610126 | 0.926829 | -2 | 0.169678 |
| P08553 | Neurofilament medi | Nefm | 715 | S | 1.2 | 0.713151 | 0.963975 | -1 | 0.919682 |

| | | | | | | | | | |
|---|---|---|---|---|---|---|---|---|---|
| P08553 | Neurofilament medi | Nefm | 769 | S | 1.6 | 0.124169 | 0.520737 | 1.6 | 0.145142 |
| P08553 | Neurofilament medi | Nefm | 610 | S | 1.1 | 0.893389 | 0.99118 | -1 | 0.067615 |
| P08553 | Neurofilament medi | Nefm | 30 | S | 1.4 | 0.575664 | 0.912975 | -1 | 0.867495 |
| P08553 | Neurofilament medi | Nefm | 715 | S | -1 | 0.85822 | 0.988571 | -1 | 0.614896 |
| O70310 | Glycylpeptide N-tetr | Nmt1 | 47 | S | -1.1 | 0.453748 | 0.859073 | -1 | 0.414708 |
| Q61937 | Nucleophosmin | Npm1 | 125 | S | 1.9 | 0.477296 | 0.874773 | -2 | 0.992367 |
| Q9CZ44 | NSFL1 cofactor p47 | Nsfl1c | 114 | S | -1.3 | 0.837621 | 0.988571 | 2.9 | 0.033248 |
| Q5F2E7 | Nuclear fragile X m | Nufip2 | 649 | S | -1.1 | 0.843973 | 0.988571 | 1.2 | 0.416343 |
| Q8VGL2 | | Olfr531 | 271 | S | 2.6 | 0.423335 | 0.848425 | 4.5 | 0.006125 |
| Q8VGL2 | | Olfr531 | 287 | T | 1.5 | 0.9026 | 0.99118 | 2.5 | 0.047265 |
| Q8VGL2 | | Olfr531 | 286 | Y | 1.4 | 0.927524 | 0.993884 | 2.5 | 0.047265 |
| Q8CI95 | Oxysterol-binding p | Osbpl11 | 186 | S | 1.2 | 0.189171 | 0.642857 | -1 | 0.140444 |
| Q4KMM3 | Oxidation resistanc | Oxr1 | 204 | S | -1 | 0.74751 | 0.963975 | 1.3 | 0.468869 |
| P50580 | Proliferation-associ | Pa2g4 | 2 | S | 1.1 | 0.891684 | 0.99118 | 1.3 | 0.223783 |
| Q8K212 | Phosphofurin acidic | Pacs1 | 28 | S | 1 | 0.865108 | 0.988571 | 1.3 | 0.389774 |
| Q3V3Q7 | Phosphofurin acidic | Pacs2 | 361 | S | 1.4 | 0.152761 | 0.571429 | 1.5 | 0.051074 |
| G5E884 | Non-specific serine/ | Pak1 | 222 | S | 1.8 | 0.21348 | 0.67101 | 1.2 | 0.969757 |
| G5E884 | Non-specific serine/ | Pak1 | 174 | S | -1.1 | 0.972342 | 0.993884 | 2.1 | 0.057261 |
| G5E884 | Non-specific serine/ | Pak1 | 224 | T | 1.8 | 0.18495 | 0.637681 | -1 | 0.875415 |
| Q9Z0P4 | Paralemmin-1 | Palm | 122 | S | -2.9 | 0.132148 | 0.530973 | -2 | 0.288491 |
| Q9Z0P4 | Paralemmin-1 | Palm | 141 | T | 1.1 | 0.738947 | 0.963975 | -1 | 0.246347 |
| Q9Z0P4 | Paralemmin-1 | Palm | 145 | T | 1.5 | 0.133394 | 0.532751 | 1.2 | 0.166916 |
| Q9Z0P4 | Paralemmin-1 | Palm | 122 | S | -2.2 | 0.34496 | 0.789352 | -2 | 0.329035 |
| Q9Z0P4 | Paralemmin-1 | Palm | 124 | S | -2.7 | 0.021157 | 0.257576 | -3 | 0.031896 |
| Q9Z0P4 | Paralemmin-1 | Palm | 157 | S | 1.2 | 0.746298 | 0.963975 | -3 | 0.476299 |
| Q9Z0P4 | Paralemmin-1 | Palm | 161 | S | -1.2 | 0.758551 | 0.963975 | -1 | 0.900671 |
| Q9Z0P4 | Paralemmin-1 | Palm | 141 | T | -1.3 | 0.761583 | 0.963975 | -1 | 0.717422 |
| Q9Z0P4 | Paralemmin-1 | Palm | 145 | T | -1.3 | 0.77213 | 0.963975 | -1 | 0.727365 |
| P60335 | Poly(rC)-binding pro | Pcbp1 | 173 | S | 1.2 | 0.733656 | 0.963975 | 1.2 | 0.542169 |
| F7BJK1 | | Pcdh1 | 1012 | S | 1.2 | 0.357682 | 0.809524 | -1 | 0.276349 |
| F7BJK1 | | Pcdh1 | 823 | S | 1.3 | 0.440339 | 0.848425 | -1 | 0.616192 |
| Q9QYX7 | Protein piccolo | Pclo | 1341 | S | -4.4 | 0.002304 | 0.090909 | -1 | 0.608138 |
| Q9QYX7 | Protein piccolo | Pclo | 1439 | S | -1 | 0.772517 | 0.963975 | -1 | 0.045003 |
| Q9QYX7 | Protein piccolo | Pclo | 1829 | S | 1.6 | 0.423671 | 0.848425 | -1 | 0.746046 |
| Q9QYX7 | Protein piccolo | Pclo | 212 | S | 1.2 | 0.799089 | 0.967352 | -2 | 0.05121 |
| Q9QYX7 | Protein piccolo | Pclo | 1772 | S | -1.3 | 0.473553 | 0.874773 | 1.3 | 0.260751 |
| Q9QYX7 | Protein piccolo | Pclo | 3376 | T | -2.1 | 0.012898 | 0.208333 | -6 | 0.000774 |
| Q9QYX7 | Protein piccolo | Pclo | 4323 | S | -1.8 | 0.17484 | 0.625 | -1 | 0.860496 |
| Q9QYX7 | Protein piccolo | Pclo | 4325 | S | -1.8 | 0.129983 | 0.530973 | -1 | 0.989946 |
| Q9QYX7 | Protein piccolo | Pclo | 1332 | S | -1.6 | 0.032599 | 0.311111 | -2 | 0.04667 |
| Q9QYX7 | Protein piccolo | Pclo | 1338 | S | -1.7 | 0.032142 | 0.311111 | -1 | 0.987553 |
| Q9QYX7 | Protein piccolo | Pclo | 3608 | S | -1.7 | 0.88519 | 0.99118 | 1.2 | 0.788758 |
| Q9QYX7 | Protein piccolo | Pclo | 3610 | S | -1.6 | 0.145532 | 0.567797 | -2 | 0.02637 |
| Q9QYX7 | Protein piccolo | Pclo | 3616 | S | -1.6 | 0.145532 | 0.567797 | -2 | 0.02637 |
| Q9QYX7 | Protein piccolo | Pclo | 1766 | S | -2.4 | 0.101368 | 0.47449 | -2 | 0.593196 |
| Q9QYX7 | Protein piccolo | Pclo | 1772 | S | -2.5 | 0.08545 | 0.439306 | -2 | 0.593196 |
| Q01065 | Calcium/calmodulin | Pde1b | 7 | S | 1.1 | 0.723271 | 0.963975 | 1.1 | 0.695576 |
| P35486 | Pyruvate dehydroge | Pdha1 | 293 | S | 3.1 | 0.24504 | 0.699708 | 6.1 | 0.000627 |
| Q9Z2A0 | 3-phosphoinositide- | Pdpk1 | 244 | S | 2.8 | 0.025999 | 0.304348 | 3 | 0.015543 |
| Q62048 | Astrocytic phospho | Pea15 | 116 | S | 1.4 | 0.440002 | 0.848425 | 2.2 | 0.066635 |
| Q8C437 | PEX5-related protei | Pex5l | 202 | S | -1.1 | 0.602242 | 0.92604 | 1.4 | 0.041985 |
| P12382 | ATP-dependent 6-p | Pfkl | 775 | S | 1.3 | 0.04437 | 0.366412 | 1.1 | 0.451996 |
| D3YZI9-2 | | Pgbd5 | 405 | S | -4.1 | 0.031895 | 0.311111 | -1 | 0.347198 |
| D3YZI9-2 | | Pgbd5 | 406 | S | 1.1 | 0.863058 | 0.988571 | -1 | 0.761206 |
| Q8CAA7 | Glucose 1,6-bispho | Pgm2l1 | 175 | S | 2.8 | 0.067002 | 0.4 | 1.8 | 0.282768 |
| Q8CAA7 | Glucose 1,6-bispho | Pgm2l1 | 173 | T | 1.9 | 0.200075 | 0.658621 | -4 | 0.009366 |
| O55022 | Membrane-associat | Pgrmc1 | 181 | S | -1 | 0.870279 | 0.988571 | -1 | 0.264671 |
| E9Q3L2 | | Pi4ka | 268 | S | 1.5 | 0.247797 | 0.700581 | -3 | 0.123654 |
| O70161 | Phosphatidylinositol | Pip5k1c | 554 | S | -1.9 | 0.616382 | 0.926975 | -2 | 0.744369 |

| | | | | | | | | | |
|---|---|---|---|---|---|---|---|---|---|
| O70161 | Phosphatidylinositol | Pip5k1c | 552 | T | 2.3 | 0.546395 | 0.902156 | 3 | 0.185076 |
| Q68FH0 | Plakophilin-4 | Pkp4 | 313 | S | 1.3 | 0.100232 | 0.471795 | 1.1 | 0.529738 |
| Q9Z1B3 | 1-phosphatidylinosit | Plcb1 | 978 | S | 8.5 | 0.050711 | 0.366412 | 18 | 0.011134 |
| Q9Z1B3 | 1-phosphatidylinosit | Plcb1 | 982 | S | 1 | 0.906631 | 0.99118 | 4.3 | 0.281383 |
| A2AP18 | 1-phosphatidylinosit | Plch2 | 676 | S | 1.1 | 0.644614 | 0.927536 | 1.1 | 0.648789 |
| Q9QXS1 | Plectin | Plec | 1443 | S | -1.3 | 0.369462 | 0.819599 | -2 | 0.156547 |
| Q9QZC2 | Plexin-C1 | Plxnc1 | 528 | S | 1.1 | 0.805067 | 0.967352 | 3.2 | 0.562211 |
| Q9CR73 | Proline-rich nuclear | Pnrc2 | 14 | S | 1.4 | 0.60446 | 0.926267 | -1 | 0.682765 |
| B2RXC6 | DNA-directed RNA | Polr3a | 1208 | S | -1 | 0.930682 | 0.993884 | 1.5 | 0.503929 |
| B2RXC6 | DNA-directed RNA | Polr3a | 1218 | S | 1 | 0.976737 | 0.993884 | 1.5 | 0.339487 |
| P60469 | Liprin-alpha-3 | Ppfia3 | 142 | S | 1.3 | 0.80705 | 0.967391 | -2 | 0.525181 |
| P60469 | Liprin-alpha-3 | Ppfia3 | 714 | T | 1.8 | 0.179465 | 0.62963 | 1.5 | 0.962034 |
| Q80TL0 | Protein phosphatase | Ppm1e | 532 | S | 1.3 | 0.620764 | 0.926975 | 2.2 | 0.068162 |
| Q60829 | Protein phosphatase | Ppp1r1b | 192 | S | 1.2 | 0.577688 | 0.912975 | -1 | 0.998854 |
| Q3UM45 | Protein phosphatase | Ppp1r7 | 24 | S | -1 | 0.93807 | 0.993884 | -1 | 0.411308 |
| Q3UM45 | Protein phosphatase | Ppp1r7 | 45 | S | -3.4 | 0.456004 | 0.861538 | 1.8 | 0.129579 |
| Q3UM45 | Protein phosphatase | Ppp1r7 | 48 | S | -3.4 | 0.440364 | 0.848425 | 1.8 | 0.142438 |
| Q3UM45 | Protein phosphatase | Ppp1r7 | 24 | S | -2.7 | 0.011171 | 0.195652 | -2 | 0.029424 |
| Q3UM45 | Protein phosphatase | Ppp1r7 | 27 | S | -2.7 | 0.011171 | 0.195652 | -2 | 0.029424 |
| Q7TN74 | | Ppp1r9a | 372 | S | -1.4 | 0.108874 | 0.495238 | 1 | 0.845505 |
| Q6R891 | Neurabin-2 | Ppp1r9b | 99 | S | 1.4 | 0.094994 | 0.47027 | 1.5 | 0.038599 |
| Q6R891 | Neurabin-2 | Ppp1r9b | 100 | S | 1.4 | 0.094994 | 0.47027 | 1.5 | 0.038599 |
| Q8R3Q2 | Serine/threonine-pro | Ppp6r2 | 669 | S | 2.5 | 0.063938 | 0.381944 | 1 | 0.566504 |
| Q922D4 | Serine/threonine-pro | Ppp6r3 | 588 | S | 1 | 0.921492 | 0.993884 | -1 | 0.860989 |
| P12367 | cAMP-dependent pr | Prkar2a | 96 | S | -1 | 0.992475 | 0.996964 | 2.1 | 0.011894 |
| P31324 | cAMP-dependent pr | Prkar2b | 112 | S | -1.5 | 0.32791 | 0.768496 | 4.3 | 0.057683 |
| Q4VA93 | Protein kinase C | Prkca | 226 | S | 2.3 | 0.067843 | 0.401361 | 2.6 | 0.19252 |
| Q4VA93 | Protein kinase C;Pr | Prkca | 319 | S | 1.3 | 0.23664 | 0.690909 | -1 | 0.328145 |
| P68404-2 | Protein kinase C be | Prkcb | 660 | S | 1.8 | 0.053234 | 0.366412 | 4.5 | 0.386663 |
| P68404-2 | Protein kinase C be | Prkcb | 641 | T | 1.7 | 0.03283 | 0.311111 | 1.9 | 0.008681 |
| P16054 | Protein kinase C ep | Prkce | 329 | S | 1.1 | 0.595767 | 0.92272 | -2 | 0.057051 |
| P16054 | Protein kinase C ep | Prkce | 334 | S | -3.7 | 0.401684 | 0.839662 | -3 | 0.569954 |
| P16054 | Protein kinase C ep | Prkce | 337 | S | -1.4 | 0.265007 | 0.713499 | -3 | 0.007096 |
| P16054 | Protein kinase C ep | Prkce | 346 | S | -2.3 | 0.273113 | 0.716535 | -2 | 0.322471 |
| P16054 | Protein kinase C ep | Prkce | 350 | S | -2.2 | 0.422972 | 0.848425 | -3 | 0.353915 |
| P63318 | Protein kinase C ga | Prkcg | 330 | S | -2.3 | 0.018166 | 0.225806 | 1.8 | 0.002956 |
| P63318 | Protein kinase C ga | Prkcg | 687 | S | -4.3 | 0.316888 | 0.753027 | -2 | 0.626777 |
| P63318 | Protein kinase C ga | Prkcg | 655 | T | 1.2 | 0.309768 | 0.753027 | 1.2 | 0.400479 |
| P63318;P6 | Protein kinase C ga | Prkcg | 514 | T | 9.8 | 0.529514 | 0.894378 | 7.4 | 0.079053 |
| Q8C5R2 | Proline and serine-r | Proser2 | 43 | S | 1.2 | 0.51227 | 0.884413 | 1.5 | 0.368673 |
| Q8C5R2 | Proline and serine-r | Proser2 | 50 | S | 1.2 | 0.511253 | 0.884413 | 1.5 | 0.37168 |
| Q8C5R2 | Proline and serine-r | Proser2 | 45 | T | 3.5 | 0.08023 | 0.427711 | 1.9 | 0.918807 |
| E9PUL5 | Proline-rich transme | Prrt2 | 98 | S | 1.2 | 0.497262 | 0.880143 | -1 | 0.963644 |
| E9PUL5 | Proline-rich transme | Prrt2 | 254 | S | 2.7 | 0.187789 | 0.641577 | 11 | 0.001259 |
| E9PUL5 | Proline-rich transme | Prrt2 | 244 | S | -1.5 | 0.225446 | 0.674923 | -2 | 0.16683 |
| E9PUL5 | Proline-rich transme | Prrt2 | 248 | S | -1.4 | 0.937295 | 0.993884 | -2 | 0.691465 |
| E9PUL5 | Proline-rich transme | Prrt2 | 250 | S | -1.2 | 0.907169 | 0.99118 | -2 | 0.244724 |
| Q6PE13 | Proline-rich transme | Prrt3 | 799 | S | 1.7 | 0.222571 | 0.674923 | 2.1 | 0.993502 |
| Q6PE13 | Proline-rich transme | Prrt3 | 845 | S | 1.1 | 0.720535 | 0.963975 | 1.1 | 0.750852 |
| E9PUC5 | | Psd3 | 44 | S | 1.5 | 0.514622 | 0.885217 | 2.4 | 0.084769 |
| E9PUC5 | PH and SEC7 doma | Psd3 | 339 | S | 2 | 0.141884 | 0.559829 | 3.7 | 0.061157 |
| E9PUC5 | PH and SEC7 doma | Psd3 | 348 | S | -1.3 | 0.273902 | 0.716535 | -1 | 0.545448 |
| E9PUC5 | PH and SEC7 doma | Psd3 | 337 | S | -3.6 | 0.001301 | 0.090909 | 1.1 | 0.976745 |
| E9PUC5 | PH and SEC7 doma | Psd3 | 339 | S | -1.7 | 0.536484 | 0.900673 | 1.3 | 0.918911 |
| E9PUC5 | PH and SEC7 doma | Psd3 | 340 | S | -1.8 | 0.56755 | 0.912975 | -1 | 0.769031 |
| E9PUC5 | PH and SEC7 doma | Psd3 | 342 | S | -3.7 | 0.001336 | 0.090909 | -1 | 0.460223 |
| E9PUC5 | PH and SEC7 doma | Psd3 | 348 | S | -1.6 | 0.009147 | 0.195652 | 1 | 0.959782 |
| E9PUC5 | PH and SEC7 doma | Psd3 | 339 | S | -4.3 | 0.002362 | 0.105263 | -1 | 0.69474 |
| E9PUC5 | PH and SEC7 doma | Psd3 | 340 | S | -4.2 | 0.002617 | 0.105263 | -1 | 0.69474 |

| | | | | | | | | | |
|---|---|---|---|---|---|---|---|---|---|
| E9PUC5 | PH and SEC7 doma | Psd3 | 348 | S | -4.6 | 0.002032 | 0.090909 | -1 | 0.69474 |
| Q99JF8 | PC4 and SFRS1-int | Psip1 | 106 | S | 1.1 | 0.748981 | 0.963975 | -1 | 0.693099 |
| Q99JF8 | PC4 and SFRS1-int | Psip1 | 171 | S | 2.6 | 0.015146 | 0.224138 | 2.1 | 0.218313 |
| Q99JF8 | PC4 and SFRS1-int | Psip1 | 176 | S | 1.1 | 0.451254 | 0.859073 | 2.1 | 0.00412 |
| O70435 | Proteasome subuni | Psma3 | 250 | S | 1.5 | 0.359221 | 0.809524 | -2 | 0.991432 |
| Q3TXS7 | 26S proteasome no | Psmd1 | 315 | S | -1.4 | 0.178728 | 0.62963 | -2 | 0.011605 |
| Q3TXS7 | 26S proteasome no | Psmd1 | 311 | T | -1.4 | 0.178728 | 0.62963 | -2 | 0.011605 |
| P34152 | Focal adhesion kina | Ptk2 | 910 | S | 1.2 | 0.38628 | 0.828326 | 1.1 | 0.694042 |
| Q9QVP9 | Protein-tyrosine kin | Ptk2b | 375 | S | 4.8 | 0.099234 | 0.471795 | 3.6 | 0.063798 |
| B9EKR1 | Receptor-type tyros | Ptprz1 | 576 | S | 1.7 | 0.576029 | 0.912975 | 2.7 | 0.270302 |
| P42669 | Transcriptional activ | Pura | 182 | T | 1.8 | 0.644372 | 0.927536 | 2.5 | 0.357098 |
| O35295 | Transcriptional activ | Purb | 6 | S | -2.1 | 0.219615 | 0.674923 | 1.1 | 0.754467 |
| O35295 | Transcriptional activ | Purb | 8 | S | -2.1 | 0.193822 | 0.645614 | 1.1 | 0.890837 |
| Q80TM6 | R3H domain-contai | R3hdm2 | 381 | S | 1.3 | 0.160033 | 0.589641 | 2.3 | 0.072612 |
| Q8BMG7 | Rab3 GTPase-activ | Rab3gap2 | 448 | S | 1.5 | 0.277281 | 0.716535 | -1 | 0.845215 |
| Q8CHG7 | Rap guanine nuclec | Rapgef2 | 1115 | S | 1.1 | 0.853077 | 0.988571 | -1 | 0.764705 |
| Q8CHG7 | Rap guanine nuclec | Rapgef2 | 930 | S | 1.5 | 0.267568 | 0.713499 | 1.7 | 0.145276 |
| F2Z3U3 | | Raph1 | 899 | S | 1.2 | 0.733537 | 0.963975 | 1.1 | 0.828643 |
| Q8C2Q3 | RNA-binding protei | Rbm14 | 206 | T | 1.4 | 0.190737 | 0.64311 | 1.6 | 0.054264 |
| Q8VH51 | RNA-binding protei | Rbm39 | 136 | S | 1.1 | 0.904172 | 0.99118 | 1.9 | 0.001673 |
| Q8VEL9 | GTP-binding protei | Rem2 | 27 | S | 1.4 | 0.207395 | 0.65894 | 1.2 | 0.245448 |
| P97492 | Regulator of G-prot | Rgs14 | 203 | S | 1 | 0.865465 | 0.988571 | 1.4 | 0.250336 |
| P97492 | Regulator of G-prot | Rgs14 | 289 | S | -1.2 | 0.570149 | 0.912975 | -1 | 0.245804 |
| Q80U40 | RIMS-binding prote | Rimbp2 | 852 | S | -1.1 | 0.807857 | 0.967431 | 2.4 | 0.052203 |
| Q99NE5 | Regulating synaptic | Rims1 | 1023 | S | -1.3 | 0.268257 | 0.713499 | -1 | 0.682518 |
| Q99NE5 | Regulating synaptic | Rims1 | 1023 | S | -2 | 0.030304 | 0.311111 | -3 | 0.003785 |
| Q99NE5 | Regulating synaptic | Rims1 | 1027 | S | -2.1 | 0.037616 | 0.336842 | -3 | 0.004888 |
| Q3V3V9 | Leucine-rich repeat- | Rltpr | 1134 | S | -2.4 | 0.003401 | 0.105263 | -2 | 0.007172 |
| Q3UJU9 | Regulator of microtu | Rmdn3 | 46 | S | 1.6 | 0.170748 | 0.610687 | 2.2 | 0.059408 |
| Q9D0L8 | mRNA cap guanine | Rnmt | 15 | S | 1.2 | 0.426982 | 0.848425 | -1 | 0.725163 |
| Q9D0L8 | mRNA cap guanine | Rnmt | 64 | S | 1.7 | 0.312996 | 0.753027 | -2 | 0.374697 |
| P47708 | Rabphilin-3A | Rph3a | 679 | S | -2.2 | 0.047107 | 0.366412 | -5 | 0.002778 |
| P47708 | Rabphilin-3A | Rph3a | 680 | S | -1.5 | 0.268306 | 0.713499 | -5 | 0.002281 |
| Q8BLK9 | Ribosomal protein S | Rps6kc1 | 778 | S | -1.2 | 0.736238 | 0.963975 | 1.4 | 0.220578 |
| Q8BLK9 | Ribosomal protein S | Rps6kc1 | 779 | S | 1.5 | 0.314514 | 0.753027 | 1.6 | 0.135741 |
| Q8K4Q0 | Regulatory-associat | Rptor | 863 | S | 1.5 | 0.26487 | 0.713499 | -1 | 0.98721 |
| Q8K0T0 | Reticulon-1 | Rtn1 | 16 | S | 1.4 | 0.291985 | 0.724936 | 1.6 | 0.35743 |
| Q8K0T0 | Reticulon-1 | Rtn1 | 350 | S | -1.1 | 0.732165 | 0.963975 | -1 | 0.553355 |
| Q8K0T0 | Reticulon-1 | Rtn1 | 352 | S | 1 | 0.962201 | 0.993884 | 1.4 | 0.056825 |
| Q8K0T0 | Reticulon-1 | Rtn1 | 92 | S | 1.6 | 0.476938 | 0.874773 | 1.9 | 0.370742 |
| Q9ES97 | Reticulon-3 | Rtn3 | 31 | S | -1.1 | 0.63141 | 0.927536 | 1.2 | 0.522267 |
| Q9ES97 | Reticulon-3 | Rtn3 | 673 | S | 1.8 | 0.000862 | 0.090909 | -1 | 0.423461 |
| Q9ES97 | Reticulon-3 | Rtn3 | 524 | T | -1.2 | 0.627362 | 0.927536 | -1 | 0.146883 |
| Q99P72 | Reticulon-4 | Rtn4 | 489 | S | -1 | 0.437229 | 0.848425 | 1.8 | 0.013271 |
| Q99P72 | Reticulon-4 | Rtn4 | 857 | S | -1.1 | 0.937616 | 0.993884 | -1 | 0.523136 |
| Q99P72 | Reticulon-4 | Rtn4 | 953 | S | 1.1 | 0.598953 | 0.92272 | 1.3 | 0.027533 |
| Q99P72 | Reticulon-4 | Rtn4 | 165 | S | 2.4 | 0.403688 | 0.84 | 3.2 | 0.327047 |
| Q99P72 | Reticulon-4 | Rtn4 | 16 | S | -1.1 | 0.453584 | 0.859073 | 1.1 | 0.548852 |
| Q99P72 | Reticulon-4 | Rtn4 | 488 | T | 1.4 | 0.514255 | 0.885017 | 1.7 | 0.067279 |
| Q99P72 | Reticulon-4 | Rtn4 | 862 | T | 1.4 | 0.31886 | 0.753027 | -1 | 0.871465 |
| Q99P72 | Reticulon-4 | Rtn4 | 171 | T | 3 | 0.083426 | 0.432749 | 1.4 | 0.828712 |
| Q99P72 | Reticulon-4 | Rtn4 | 165 | S | -1.3 | 0.914513 | 0.992341 | 1.4 | 0.659811 |
| Q99P72 | Reticulon-4 | Rtn4 | 167 | S | -1.6 | 0.413811 | 0.848425 | 1.2 | 0.889559 |
| Q99P72 | Reticulon-4 | Rtn4 | 171 | T | -1.5 | 0.638285 | 0.927536 | 1.2 | 0.863056 |
| D3YXK2 | Scaffold attachmen | Safb | 366 | S | 1.3 | 0.437118 | 0.848425 | 1.8 | 0.07421 |
| D3YXK2 | Scaffold attachmen | Safb | 372 | S | 1.5 | 0.09891 | 0.471795 | 1.7 | 0.010239 |
| Q6ZPE2 | Myotubularin-relate | Sbf1 | 706 | S | 3 | 0.184766 | 0.637681 | 2.5 | 0.051136 |
| O35609 | Secretory carrier-as | Scamp3 | 78 | S | 1.3 | 0.112644 | 0.495238 | 1.7 | 0.01183 |
| O08547 | Vesicle-trafficking p | Sec22b | 137 | S | 1.1 | 0.820193 | 0.975962 | -1 | 0.697573 |

| | | | | | | | | | |
|---|---|---|---|---|---|---|---|---|---|
| Q9CQS8 | Protein transport pr | Sec61b | 17 | S | 1.6 | 0.063393 | 0.381944 | 1.2 | 0.313229 |
| Q64213 | Splicing factor 1 | Sf1 | 80 | S | -2.5 | 0.22542 | 0.674923 | 1.2 | 0.395615 |
| Q8BP27 | Swi5-dependent rec | Sfr1 | 67 | S | -2 | 0.006344 | 0.195652 | -2 | 0.028575 |
| Q8BP27 | Swi5-dependent rec | Sfr1 | 71 | S | -3.3 | 0.441565 | 0.850688 | 1.4 | 0.545211 |
| Q8BP27 | Swi5-dependent rec | Sfr1 | 70 | T | -1.7 | 0.569912 | 0.912975 | -2 | 0.326761 |
| Q8VD37 | SH3-containing GR | Sgip1 | 169 | S | 1.4 | 0.10675 | 0.490099 | 1.3 | 0.331352 |
| Q8VD37 | SH3-containing GR | Sgip1 | 265 | S | -1.1 | 0.983455 | 0.993884 | 1.3 | 0.861959 |
| Q8VD37 | SH3-containing GR | Sgip1 | 335 | T | 3.1 | 0.155842 | 0.578947 | 2.6 | 0.00303 |
| Q8VD37 | SH3-containing GR | Sgip1 | 263 | T | -1.8 | 0.717838 | 0.963975 | -2 | 0.848298 |
| Q8VD37 | SH3-containing GR | Sgip1 | 263 | T | 1.1 | 0.982996 | 0.993884 | -1 | 0.993667 |
| Q8VD37 | SH3-containing GR | Sgip1 | 259 | T | 1.1 | 0.94203 | 0.993884 | 1.1 | 0.448601 |
| Q8R3V5 | Endophilin-B2 | Sh3glb2 | 400 | S | -1 | 0.985303 | 0.994908 | 1.7 | 0.10107 |
| D3YZU1 | SH3 and multiple ar | Shank1 | 413 | S | -1 | 0.919922 | 0.993884 | -1 | 0.320454 |
| Q4ACU6 | SH3 and multiple ar | Shank3 | 781 | S | 1.2 | 0.510126 | 0.884413 | 3.1 | 0.020446 |
| Q3UH99 | Protein shisa-6 hom | Shisa6 | 247 | S | 1.7 | 0.667206 | 0.940678 | 1.7 | 0.419218 |
| Q3UH99 | Protein shisa-6 hom | Shisa6 | 246 | T | 2.1 | 0.087755 | 0.44382 | -2 | 0.933153 |
| Q3UH99 | Protein shisa-6 hom | Shisa6 | 252 | T | -1.3 | 0.990447 | 0.995947 | -2 | 0.893445 |
| Q8C0T5 | Signal-induced proli | Sipa1l1 | 1528 | S | 1.2 | 0.548103 | 0.902156 | 1.1 | 0.535393 |
| Q8C0T5 | Signal-induced proli | Sipa1l1 | 1564 | S | -1.5 | 0.894673 | 0.99118 | -5 | 0.421794 |
| Q8VDQ8 | NAD-dependent pro | Sirt2 | 372 | S | 4.8 | 0.454479 | 0.859345 | 7.4 | 0.092842 |
| P53986 | Monocarboxylate tra | Slc16a1 | 491 | S | -1 | 0.713008 | 0.963975 | -1 | 0.557463 |
| P53986 | Monocarboxylate tra | Slc16a1 | 210 | S | -1.5 | 0.278568 | 0.716535 | -2 | 0.201613 |
| P53986 | Monocarboxylate tra | Slc16a1 | 213 | S | -1.5 | 0.260351 | 0.70904 | -2 | 0.179989 |
| P56564 | Excitatory amino ac | Slc1a3 | 512 | S | 1.3 | 0.565025 | 0.912975 | 1.1 | 0.647969 |
| Q8BUN9 | | Slc24a2 | 312 | S | 1.1 | 0.833374 | 0.986874 | -2 | 0.80137 |
| Q8BUN9 | | Slc24a2 | 334 | S | -1.5 | 0.232948 | 0.690909 | -1 | 0.276156 |
| Q8BUN9 | | Slc24a2 | 337 | S | -1.5 | 0.232948 | 0.690909 | -1 | 0.276156 |
| O35633 | Vesicular inhibitory | Slc32a1 | 97 | S | 3.6 | 0.087474 | 0.44382 | 1.1 | 0.728177 |
| Q5DTL9 | Sodium-driven chlor | Slc4a10 | 91 | S | -2.2 | 0.508651 | 0.884413 | -2 | 0.409949 |
| Q8BJI1 | Sodium-dependent | Slc6a17 | 665 | S | -1 | 0.612876 | 0.926975 | -2 | 0.008438 |
| Q8BXR1 | Probable cationic ar | Slc7a14 | 769 | S | -2.5 | 0.310615 | 0.753027 | 1.1 | 0.876046 |
| P70441 | Na(+)/H(+) exchang | Slc9a3r1 | 275 | S | 1.3 | 0.682177 | 0.956522 | 1.9 | 0.0019 |
| A1L3P4 | Sodium/hydrogen e | Slc9a6 | 700 | S | 3.2 | 0.462468 | 0.868069 | 2.2 | 0.111191 |
| Q9R0P4 | Small acidic protein | Smap | 15 | S | -1.2 | 0.810616 | 0.968675 | -1 | 0.556924 |
| Q9R0P4 | Small acidic protein | Smap | 17 | S | -1.2 | 0.967218 | 0.993884 | -2 | 0.406346 |
| Q6PDG5 | SWI/SNF complex s | Smarcc2 | 302 | S | -1.2 | 0.806689 | 0.967352 | -2 | 0.104495 |
| Q6PDG5 | SWI/SNF complex s | Smarcc2 | 304 | S | -1.2 | 0.784358 | 0.963975 | -2 | 0.314406 |
| Q6PDG5 | SWI/SNF complex s | Smarcc2 | 306 | S | -1.1 | 0.898046 | 0.99118 | -3 | 0.020531 |
| Q61548 | Clathrin coat assem | Snap91 | 300 | S | 1.3 | 0.187712 | 0.641577 | -1 | 0.533461 |
| Q61548 | Clathrin coat assem | Snap91 | 313 | S | 2.1 | 0.113753 | 0.495238 | 1.3 | 0.672515 |
| Q61548 | Clathrin coat assem | Snap91 | 321 | T | 5.1 | 0.57137 | 0.912975 | 5.1 | 0.185898 |
| Q61548 | Clathrin coat assem | Snap91 | 296 | S | -1.5 | 0.065221 | 0.381944 | -2 | 0.019793 |
| Q61548 | Clathrin coat assem | Snap91 | 300 | S | -1.5 | 0.065221 | 0.381944 | -2 | 0.019793 |
| Q61234 | Alpha-1-syntrophin | Snta1 | 183 | S | 1.5 | 0.023525 | 0.279412 | 1.5 | 0.024688 |
| Q8BVL3 | Sorting nexin-17 | Snx17 | 331 | S | 1.4 | 0.377762 | 0.828326 | 1.2 | 0.464992 |
| Q3UTJ2 | Sorbin and SH3 dor | Sorbs2 | 829 | S | -1 | 0.73651 | 0.963975 | 2.2 | 0.000713 |
| Q3UTJ2 | Sorbin and SH3 dor | Sorbs2 | 382 | S | -5.1 | 0.160175 | 0.589641 | -5 | 0.225131 |
| Q3UTJ2 | Sorbin and SH3 dor | Sorbs2 | 377 | S | -3.6 | 0.080993 | 0.427711 | -1 | 0.767176 |
| A0A5H1ZF | Protein spire homol | Spire1 | 663 | S | 1.2 | 0.649048 | 0.932277 | 2.3 | 0.007984 |
| P16546 | Spectrin alpha chair | Sptan1 | 1029 | S | 4.3 | 0.013222 | 0.211538 | 8.2 | 0.075405 |
| P16546 | Spectrin alpha chair | Sptan1 | 1031 | S | 1.5 | 0.500439 | 0.880143 | 1.9 | 0.768404 |
| Q62261 | Spectrin beta chain, | Sptbn1 | 2340 | S | 1.5 | 0.484081 | 0.874773 | 1.2 | 0.998893 |
| Q62261 | Spectrin beta chain, | Sptbn1 | 2137 | S | -1.2 | 0.869599 | 0.988571 | -1 | 0.547154 |
| Q62261 | Spectrin beta chain, | Sptbn1 | 2102 | S | 2 | 0.033828 | 0.311111 | 1.3 | 0.320628 |
| Q62261 | Spectrin beta chain, | Sptbn1 | 2168 | S | -1.4 | 0.954497 | 0.993884 | -1 | 0.853024 |
| Q62261 | Spectrin beta chain, | Sptbn1 | 2163 | S | -3.5 | 0.01233 | 0.195652 | -2 | 0.084734 |
| Q62261 | Spectrin beta chain, | Sptbn1 | 2164 | S | -3.6 | 0.011476 | 0.195652 | -2 | 0.070271 |
| Q62261 | Spectrin beta chain, | Sptbn1 | 2168 | S | -3.8 | 0.010383 | 0.195652 | -2 | 0.085487 |
| P05480 | Neuronal proto-onco | Src | 74 | S | 1.2 | 0.685588 | 0.956522 | -1 | 0.102227 |

| | | | | | | | | | |
|---|---|---|---|---|---|---|---|---|---|
| Q9QWI6 | SRC kinase signalir | Srcin1 | 588 | S | -1.2 | 0.31788 | 0.753027 | -1 | 0.220332 |
| Q9QWI6 | SRC kinase signalir | Srcin1 | 233 | S | 1.2 | 0.3056 | 0.748744 | 1.2 | 0.326162 |
| Q9QWI6 | SRC kinase signalir | Srcin1 | 1110 | S | -1.3 | 0.164865 | 0.6 | 1.5 | 0.006378 |
| Q9QWI6 | SRC kinase signalir | Srcin1 | 1054 | S | -1.1 | 0.629848 | 0.927536 | 1.2 | 0.233585 |
| Q52KI8 | Serine/arginine repe | Srrm1 | 260 | S | 1.2 | 0.43991 | 0.848425 | 1.5 | 0.029459 |
| Q52KI8 | Serine/arginine repe | Srrm1 | 401 | S | 1.1 | 0.625669 | 0.927536 | -1 | 0.359503 |
| Q52KI8 | Serine/arginine repe | Srrm1 | 387 | S | 1.1 | 0.672516 | 0.944993 | -1 | 0.941067 |
| Q52KI8 | Serine/arginine repe | Srrm1 | 391 | S | 1.1 | 0.668254 | 0.940678 | -1 | 0.929989 |
| Q8BTI8 | Serine/arginine repe | Srrm2 | 1269 | S | -1.7 | 0.149164 | 0.571429 | 1.1 | 0.644683 |
| Q8BTI8 | Serine/arginine repe | Srrm2 | 1305 | S | 1 | 0.841351 | 0.988571 | -1 | 0.311872 |
| Q8BTI8 | Serine/arginine repe | Srrm2 | 1359 | S | 1 | 0.852114 | 0.988571 | -3 | 0.004497 |
| Q8BTI8 | Serine/arginine repe | Srrm2 | 1360 | S | -1.8 | 0.524077 | 0.894378 | 1.4 | 0.369657 |
| Q99KH8 | Serine/threonine-pr | Stk24 | 4 | S | 1.5 | 0.269383 | 0.714286 | 1 | 0.941018 |
| P54227 | Stathmin;Stathmin-: | Stmn1 | 46 | S | -1.3 | 0.207799 | 0.65894 | 1.1 | 0.956345 |
| P54227 | Stathmin | Stmn1 | 38 | S | 1.6 | 0.154944 | 0.577236 | 2.6 | 0.006101 |
| Q8C079 | Striatin-interacting p | Strip1 | 335 | S | 1.3 | 0.095556 | 0.47027 | -1 | 0.400375 |
| O55106 | Striatin | Strn | 245 | S | -1.6 | 0.434994 | 0.848425 | 1 | 0.70577 |
| P58404 | Striatin-4 | Strn4 | 223 | S | 1.5 | 0.09757 | 0.471795 | 1.4 | 0.12593 |
| Q9WUD1 | STIP1 homology an | Stub1 | 20 | S | 1.1 | 0.983506 | 0.993884 | -1 | 0.223415 |
| Q9WUD1 | STIP1 homology an | Stub1 | 15 | T | 1.2 | 0.877698 | 0.99118 | -2 | 0.590864 |
| O35526 | Syntaxin-1A | Stx1a | 14 | S | -2.4 | 0.047354 | 0.366412 | -1 | 0.839308 |
| O35526 | Syntaxin-1A | Stx1a | 10 | T | -1.3 | 0.582393 | 0.916272 | -1 | 0.660433 |
| P61264 | Syntaxin-1B | Stx1b | 14 | S | -4.9 | 0.015775 | 0.224138 | -1 | 0.839039 |
| P70452 | Syntaxin-4 | Stx4 | 15 | S | -1.2 | 0.557225 | 0.912975 | -1 | 0.864562 |
| O08599 | Syntaxin-binding pr | Stxbp1 | 593 | S | -1.2 | 0.221517 | 0.674923 | -1 | 0.688456 |
| O08599 | Syntaxin-binding pr | Stxbp1 | 594 | S | -2.3 | 0.132232 | 0.530973 | 1.2 | 0.539368 |
| O08599-2 | Syntaxin-binding pr | Stxbp1 | 590 | S | 1.9 | 0.065235 | 0.381944 | -1 | 0.205767 |
| Q8K400 | Syntaxin-binding pr | Stxbp5 | 760 | S | 1.9 | 0.360935 | 0.810811 | 3.2 | 0.033162 |
| P63166 | Small ubiquitin-relat | Sumo1 | 2 | S | 1.1 | 0.449653 | 0.859073 | -1 | 0.08873 |
| Q9JIS5 | Synaptic vesicle gly | Sv2a | 127 | S | -1.3 | 0.367996 | 0.818792 | -1 | 0.253155 |
| O88935 | Synapsin-1 | Syn1 | 67 | S | 1.5 | 0.277547 | 0.716535 | 3.8 | 0.039507 |
| O88935 | Synapsin-1 | Syn1 | 427 | S | 1.8 | 0.391337 | 0.832976 | 3.7 | 0.00241 |
| O88935 | Synapsin-1 | Syn1 | 438 | S | 1.3 | 0.895379 | 0.99118 | 2 | 0.198647 |
| O88935;Q | Synapsin-1;Synapsi | Syn1 | 705 | S | 1.1 | 0.770379 | 0.963975 | -1 | 0.597659 |
| O88935 | Synapsin-1 | Syn1 | 39 | S | -1.2 | 0.349901 | 0.799539 | 1.5 | 0.213653 |
| O88935 | Synapsin-1 | Syn1 | 551 | S | -1.2 | 0.513936 | 0.884817 | 2.2 | 0.114007 |
| O88935 | Synapsin-1 | Syn1 | 553 | S | 1.4 | 0.987543 | 0.995935 | 4 | 0.040424 |
| O88935 | Synapsin-1 | Syn1 | 568 | S | -2.2 | 0.854778 | 0.988571 | -2 | 0.509616 |
| O88935 | Synapsin-1 | Syn1 | 666 | S | 2.1 | 0.190081 | 0.64311 | 2.7 | 0.219414 |
| O88935;Q | Synapsin-1;Synapsi | Syn1 | 341 | S | 1.3 | 0.758886 | 0.963975 | 1.4 | 0.654495 |
| O88935 | Synapsin-1 | Syn1 | 512 | T | 1.8 | 0.20727 | 0.65894 | -1 | 0.881936 |
| O88935 | Synapsin-1 | Syn1 | 434 | S | -1 | 0.747151 | 0.963975 | -2 | 0.385441 |
| O88935 | Synapsin-1 | Syn1 | 438 | S | -1.3 | 0.783386 | 0.963975 | -2 | 0.201338 |
| Q64332 | Synapsin-2 | Syn2 | 426 | S | 1.4 | 0.359437 | 0.809524 | 2 | 0.009785 |
| Q64332 | Synapsin-2 | Syn2 | 422 | T | 2 | 0.86593 | 0.988571 | ### | 0.077763 |
| Q64332 | Synapsin-2 | Syn2 | 426 | S | -1.5 | 0.126039 | 0.522936 | -1 | 0.605627 |
| Q64332 | Synapsin-2 | Syn2 | 422 | T | 1 | 0.895273 | 0.99118 | -2 | 0.173193 |
| Q8JZP2 | Synapsin-3 | Syn3 | 461 | S | -1.4 | 0.279102 | 0.716535 | -2 | 0.027811 |
| Q8JZP2 | Synapsin-3 | Syn3 | 483 | S | 1 | 0.547527 | 0.902156 | -1 | 0.097504 |
| Q8JZP2 | Synapsin-3 | Syn3 | 538 | S | 1.5 | 0.194715 | 0.646853 | -3 | 0.051161 |
| Q8JZP2 | Synapsin-3 | Syn3 | 540 | S | 1.6 | 0.205593 | 0.65894 | -2 | 0.102981 |
| F6SEU4 | Ras/Rap GTPase-a | Syngap1 | 766 | S | -1.2 | 0.52359 | 0.894378 | -1 | 0.635881 |
| E9Q7S0 | Synaptojanin-1 | Synj1 | 1084 | S | 1.3 | 0.418929 | 0.848425 | -1 | 0.860711 |
| E9Q7S0 | Synaptojanin-1 | Synj1 | 1075 | T | 8.5 | 0.157121 | 0.589641 | 5.3 | 0.23946 |
| Q8CC35 | Synaptopodin | Synpo | 258 | S | 1.4 | 0.963081 | 0.993884 | 2.3 | 0.022767 |
| Q62277 | Synaptophysin | Syp | 81 | Y | -1.2 | 0.222804 | 0.674923 | 1.6 | 0.008168 |
| P46097 | Synaptotagmin-2 | Syt2 | 128 | T | -1.1 | 0.899514 | 0.99118 | 1.5 | 0.684788 |
| A2A690 | Protein TANC2 | Tanc2 | 400 | S | -1.1 | 0.90637 | 0.99118 | -2 | 0.411247 |
| A2A690 | Protein TANC2 | Tanc2 | 404 | S | -1.2 | 0.46231 | 0.868069 | -2 | 0.073603 |

| | | | | | | | | | |
|---|---|---|---|---|---|---|---|---|---|
| Q8BHL3 | TBC1 domain famil | Tbc1d10b | 695 | T | 1.2 | 0.638884 | 0.927536 | 2.5 | 0.543024 |
| E9Q1W7 | | Tbck | 890 | S | 1 | 0.928187 | 0.993884 | -1 | 0.935291 |
| Q8CCT4 | Transcription elong | Tceal5 | 120 | S | -1.1 | 0.499497 | 0.880143 | -1 | 0.416515 |
| Q8CCT4 | Transcription elong | Tceal5 | 117 | T | 1.5 | 0.488671 | 0.874773 | 1.3 | 0.300913 |
| Q569Z6 | Thyroid hormone re | Thrap3 | 243 | S | 1.3 | 0.276582 | 0.716535 | -1 | 0.178533 |
| Q569Z6 | Thyroid hormone re | Thrap3 | 679 | S | 1.1 | 0.740614 | 0.963975 | 1.4 | 0.043299 |
| Q569Z6 | Thyroid hormone re | Thrap3 | 572 | S | -1.1 | 0.925321 | 0.993884 | 1.8 | 0.503011 |
| Q99J36 | THUMP domain-co | Thumpd1 | 8 | S | 1.2 | 0.358489 | 0.809524 | -2 | 0.009869 |
| Q9Z0U1 | Tight junction prote | Tjp2 | 107 | S | 1.1 | 0.879648 | 0.99118 | 1.2 | 0.924232 |
| P83510 | Traf2 and NCK-inte | Tnik | 611 | S | 1.1 | 0.613291 | 0.926975 | -1 | 0.826188 |
| P83510 | Traf2 and NCK-inte | Tnik | 740 | S | -1 | 0.67876 | 0.95493 | 1.1 | 0.611246 |
| P58871 | 182 kDa tankyrase- | Tnks1bp1 | 1437 | S | -1.1 | 0.978311 | 0.993884 | -1 | 0.958789 |
| P58871 | 182 kDa tankyrase- | Tnks1bp1 | 1440 | S | -2 | 0.845052 | 0.988571 | -2 | 0.819266 |
| Q8BYI9 | Tenascin-R | Tnr | 727 | S | 1.2 | 0.944759 | 0.993884 | 1.5 | 0.561342 |
| Q5SRX1 | TOM1-like protein 2 | Tom1l2 | 479 | S | -2.7 | 0.016284 | 0.224138 | 1.7 | 0.023248 |
| Q5SRX1 | TOM1-like protein 2 | Tom1l2 | 160 | S | -1.5 | 0.178209 | 0.62963 | 3 | 0.410539 |
| Q9CPQ3 | Mitochondrial impor | Tomm22 | 15 | S | 2.2 | 0.05076 | 0.366412 | 4.6 | 0.004193 |
| Q9CZW5 | Mitochondrial impor | Tomm70a | 94 | S | -1.1 | 0.61714 | 0.926975 | 2.2 | 0.015427 |
| P70399 | Tumor suppressor | Tp53bp1 | 552 | S | -1.2 | 0.437797 | 0.848425 | -1 | 0.545319 |
| Q9CYZ2 | Tumor protein D54 | Tpd52l2 | 12 | S | 1 | 0.637937 | 0.927536 | -2 | 0.004641 |
| Q7TQD2 | Tubulin polymerizat | Tppp | 159 | S | 2.3 | 0.397929 | 0.836864 | 1.8 | 0.118806 |
| Q7TQD2 | Tubulin polymerizat | Tppp | 15 | T | 2 | 0.123191 | 0.520737 | 2.7 | 0.001342 |
| Q7TQD2 | Tubulin polymerizat | Tppp | 19 | S | 1.3 | 0.321853 | 0.753027 | -1 | 0.93834 |
| Q7TQD2 | Tubulin polymerizat | Tppp | 15 | T | 1.3 | 0.321853 | 0.753027 | -1 | 0.93834 |
| Q9DBS2 | Tumor protein p63-r | Tprg1l | 10 | S | 1.3 | 0.904259 | 0.99118 | 1.1 | 0.996696 |
| Q9DBS2 | Tumor protein p63-r | Tprg1l | 34 | T | 2.2 | 0.011032 | 0.195652 | 3.6 | 0.004206 |
| Q9DBS2 | Tumor protein p63-r | Tprg1l | 10 | S | -1.1 | 0.754224 | 0.963975 | -1 | 0.30151 |
| Q9DBS2 | Tumor protein p63-r | Tprg1l | 34 | T | -1.1 | 0.754224 | 0.963975 | -1 | 0.356044 |
| Q9ESN6 | Tripartite motif-cont | Trim2 | 428 | S | 1.4 | 0.227058 | 0.674923 | -1 | 0.980116 |
| Q62318 | Transcription interm | Trim28 | 23 | S | 1.3 | 0.436458 | 0.848425 | -1 | 0.69167 |
| E9Q6P5 | | Ttc7b | 160 | S | 1.4 | 0.218478 | 0.674923 | 1.4 | 0.268422 |
| E9Q6P5 | | Ttc7b | 678 | S | 1.2 | 0.423837 | 0.848425 | 1.6 | 0.023923 |
| Q3UDE2 | Tubulin--tyrosine lig | Ttll12 | 11 | S | -1.1 | 0.528506 | 0.894378 | -2 | 0.050275 |
| P05213;P6 | Tubulin alpha-1B ch | Tuba1b | 48 | S | 1.3 | 0.621992 | 0.926975 | 2 | 0.012159 |
| P68368;P( | Tubulin alpha-4A ch | Tuba4a | 340 | S | 1.8 | 0.133112 | 0.532751 | 1.9 | 0.221801 |
| Q02053 | Ubiquitin-like modifi | Uba1 | 835 | S | 1.2 | 0.441768 | 0.85098 | 2.2 | 0.003241 |
| Q6ZPJ3 | E2/E3 hybrid ubiqui | Ube2o | 836 | S | 1.9 | 0.237801 | 0.692771 | 2.6 | 0.04463 |
| P52479 | Ubiquitin carboxyl-te | Usp10 | 208 | S | 1.3 | 0.376177 | 0.824176 | 1.1 | 0.99614 |
| P52479 | Ubiquitin carboxyl-te | Usp10 | 205 | T | 1 | 0.776573 | 0.963975 | 1.3 | 0.728879 |
| Q9JMA1 | Ubiquitin carboxyl-te | Usp14 | 143 | S | 1.9 | 0.537855 | 0.900673 | 2 | 0.032128 |
| P56399 | Ubiquitin carboxyl-te | Usp5 | 783 | S | -1.2 | 0.284868 | 0.718346 | -1 | 0.676086 |
| P56399 | Ubiquitin carboxyl-te | Usp5 | 623 | T | 1.1 | 0.79534 | 0.966952 | 4.4 | 0.005815 |
| O70480 | Vesicle-associated | Vamp4 | 30 | S | -1.1 | 0.383692 | 0.828326 | 1.2 | 0.847583 |
| Q60932 | Voltage-dependent | Vdac1 | 117 | S | 1.4 | 0.146505 | 0.571429 | 1.6 | 0.048256 |
| Q8C0E2 | Vacuolar protein so | Vps26b | 302 | S | 1.9 | 0.010644 | 0.195652 | 1.6 | 0.028551 |
| Q9EQH3 | Vacuolar protein so | Vps35 | 7 | S | -1.8 | 0.41376 | 0.848425 | -2 | 0.650812 |
| Q3UVL4 | Vacuolar protein so | Vps51 | 18 | S | -1.2 | 0.280431 | 0.716535 | 1.2 | 0.930427 |
| Q6NVE8 | WD repeat-containi | Wdr44 | 50 | S | -1.3 | 0.226976 | 0.674923 | -2 | 0.077086 |
| Q6NVE8 | WD repeat-containi | Wdr44 | 405 | S | -1.1 | 0.723625 | 0.963975 | 1.6 | 0.217332 |
| P0C7L0 | WAS/WASL-interac | Wipf3 | 211 | S | 1.7 | 0.204776 | 0.65894 | 1.3 | 0.163197 |

| ROTS FDR |
|---|
| 0.103774 |
| 0.169811 |
| 0.958515 |
| 0.238298 |
| 0.368078 |
| 0.744681 |
| 0.456897 |
| 0.494845 |
| 0.224299 |
| 0.478495 |
| 0.37013 |
| 0.417683 |
| 0.503667 |
| 0.524887 |
| 0.310469 |
| 0.169811 |
| 0.955407 |
| 0.346801 |
| 0.471751 |
| 0.917397 |
| 0.378981 |
| 0.672694 |
| 0.962121 |
| 0.672694 |
| 0.776886 |
| 0.139706 |
| 0.485488 |
| 0.815249 |
| 0.708621 |
| 0.90781 |
| 0.503667 |
| 0.485488 |
| 0.238298 |
| 0.324042 |
| 0.793103 |
| 0.955407 |
| 0.793103 |
| 0.793103 |
| 0.070423 |
| 0.129032 |
| 0.744681 |
| 0.958515 |
| 0.070423 |
| 0.070423 |
| 0.90781 |
| 0.564211 |
| 0.503667 |
| 0.129032 |
| 0.623552 |
| 0.510638 |
| 0.856745 |
| 0.951807 |
| 0.46 |
| 0.955407 |
| 0.955703 |
| 0.955407 |
| 0.1875 |

| |
|---|
| 0.504762 |
| 0.955407 |
| 0.797214 |
| 0.785714 |
| 0.227679 |
| 0.437126 |
| 0.911728 |
| 0.475275 |
| 0.955407 |
| 0.958515 |
| 0.997976 |
| 0.052632 |
| 0.955407 |
| 0.953125 |
| 0.57438 |
| 0.052632 |
| 0.857341 |
| 0.142336 |
| 0.997976 |
| 0.445104 |
| 0.836415 |
| 0.052632 |
| 0.378981 |
| 0.564211 |
| 0.197861 |
| 0.911728 |
| 0.611328 |
| 0.564211 |
| 0.701209 |
| 0.955407 |
| 0.324042 |
| 0.985447 |
| 0.995889 |
| 0.058824 |
| 0.955407 |
| 0.975866 |
| 0.875676 |
| 0.611328 |
| 0.103774 |
| 0.878869 |
| 0.975866 |
| 0.968085 |
| 0.516355 |
| 0.955407 |
| 0.758958 |
| 0.49354 |
| 0.5 |
| 0.143885 |
| 0.744681 |
| 0.716724 |
| 0.169811 |
| 0.817251 |
| 0.955407 |
| 0.964668 |
| 0.327645 |
| 0.227679 |
| 0.997976 |
| 0.785256 |
| 0.590631 |

| |
|---|
| 0.955407 |
| 0.955407 |
| 0.929104 |
| 0.744681 |
| 0.136364 |
| 0.058824 |
| 0.058824 |
| 0.052632 |
| 0.070423 |
| 0.911728 |
| 0.911728 |
| 0.932681 |
| 0.347826 |
| 0.997976 |
| 0.147887 |
| 0.955407 |
| 0.564211 |
| 0.985447 |
| 0.089888 |
| 0.05 |
| 0.258964 |
| 0.644068 |
| 0.503667 |
| 0.564211 |
| 0.1875 |
| 0.5 |
| 0.823188 |
| 0.716724 |
| 0.955407 |
| 0.070423 |
| 0.070423 |
| 0.35 |
| 0.995881 |
| 0.304348 |
| 0.941606 |
| 0.504762 |
| 0.932681 |
| 0.975661 |
| 0.797214 |
| 0.817784 |
| 0.955407 |
| 0.701209 |
| 0.510638 |
| 0.227679 |
| 0.951807 |
| 0.545657 |
| 0.885906 |
| 0.649813 |
| 0.134921 |
| 0.945322 |
| 0.785714 |
| 0.785714 |
| 0.785714 |
| 0.327645 |
| 0.327645 |
| 0.227679 |
| 0.744681 |
| 0.949214 |
| 0.744681 |

| |
|---|
| 0.744681 |
| 0.552106 |
| 0.644068 |
| 0.950483 |
| 0.962366 |
| 0.597561 |
| 0.052632 |
| 0.744681 |
| 0.836415 |
| 0.836415 |
| 0.955407 |
| 0.6238 |
| 0.857341 |
| 0.05 |
| 0.955407 |
| 0.895833 |
| 0.955407 |
| 0.05 |
| 0.221698 |
| 0.997976 |
| 0.733447 |
| 0.266667 |
| 0.058824 |
| 0.823188 |
| 0.895833 |
| 0.807284 |
| 0.957143 |
| 0.955407 |
| 0.968085 |
| 0.844193 |
| 0.08642 |
| 0.76013 |
| 0.955407 |
| 0.836415 |
| 0.473389 |
| 0.478495 |
| 0.811377 |
| 0.238298 |
| 0.503667 |
| 0.503667 |
| 0.089888 |
| 0.089888 |
| 0.611111 |
| 0.955407 |
| 0.564211 |
| 0.05 |
| 0.917397 |
| 0.955407 |
| 0.997976 |
| 0.248963 |
| 0.564211 |
| 0.136364 |
| 0.997976 |
| 0.052632 |
| 0.205 |
| 0.744681 |
| 0.955407 |
| 0.05 |
| 0.697552 |

0.197861
0.569038
0.849372
0.814815
0.214634
0.661172
0.96463
0.849372
0.76013
0.680357
0.516355
0.503667
0.503667
0.503667
0.129032
0.407975
0.622824
0.478142
0.264822
0.504762
0.324042
0.058824
0.930521
0.492188
0.743676
0.169811
0.870166
0.836415
0.4375
0.611111
0.590631
0.744681
0.092784
0.955407
0.658627
0.289963
0.737288
0.117117
0.879032
0.672694
0.911728
0.478495
0.955407
0.227679
0.953125
0.467236
0.052632
0.990712
0.917397
0.90781
0.962366
0.962366
0.962366
0.829233
0.744681
0.776886
0.082192
0.258964
0.955407

| |
|---|
| 0.806697 |
| 0.576132 |
| 0.697552 |
| 0.524887 |
| 0.092784 |
| 0.05 |
| 0.05 |
| 0.895833 |
| 0.90781 |
| 0.534676 |
| 0.644068 |
| 0.070423 |
| 0.103774 |
| 0.143885 |
| 0.103774 |
| 0.874317 |
| 0.117117 |
| 0.762136 |
| 0.985447 |
| 0.485488 |
| 0.488189 |
| 0.975661 |
| 0.180233 |
| 0.960954 |
| 0.070423 |
| 0.10101 |
| 0.180233 |
| 0.070423 |
| 0.070423 |
| 0.05 |
| 0.258964 |
| 0.347826 |
| 0.556291 |
| 0.205 |
| 0.59919 |
| 0.052632 |
| 0.650558 |
| 0.803354 |
| 0.052632 |
| 0.272727 |
| 0.968085 |
| 0.56044 |
| 0.875676 |
| 0.05 |
| 0.177914 |
| 0.564211 |
| 0.799383 |
| 0.49354 |
| 0.486842 |
| 0.949214 |
| 0.244726 |
| 0.214634 |
| 0.955407 |
| 0.214634 |
| 0.955407 |
| 0.765751 |
| 0.136364 |
| 0.136364 |
| 0.4375 |

| |
|---|
| 0.932681 |
| 0.697552 |
| 0.189944 |
| 0.276119 |
| 0.911728 |
| 0.238298 |
| 0.356436 |
| 0.955407 |
| 0.81194 |
| 0.070423 |
| 0.449275 |
| 0.811377 |
| 0.673874 |
| 0.516355 |
| 0.90781 |
| 0.949214 |
| 0.92 |
| 0.917397 |
| 0.57438 |
| 0.823188 |
| 0.180233 |
| 0.650558 |
| 0.450867 |
| 0.803354 |
| 0.518605 |
| 0.358553 |
| 0.564211 |
| 0.129032 |
| 0.563043 |
| 0.59919 |
| 0.180233 |
| 0.776167 |
| 0.475275 |
| 0.932681 |
| 0.219048 |
| 0.475275 |
| 0.895833 |
| 0.504762 |
| 0.274436 |
| 0.375405 |
| 0.955407 |
| 0.815249 |
| 0.103774 |
| 0.504762 |
| 0.744681 |
| 0.524887 |
| 0.089888 |
| 0.525959 |
| 0.05 |
| 0.343537 |
| 0.812221 |
| 0.895833 |
| 0.81454 |
| 0.960954 |
| 0.955407 |
| 0.744681 |
| 0.875676 |
| 0.270992 |
| 0.270992 |

| |
|---|
| 0.803354 |
| 0.964592 |
| 0.955506 |
| 0.304348 |
| 0.955407 |
| 0.483957 |
| 0.90781 |
| 0.327645 |
| 0.955407 |
| 0.485488 |
| 0.797214 |
| 0.180233 |
| 0.744681 |
| 0.266667 |
| 0.264822 |
| 0.05 |
| 0.955407 |
| 0.955407 |
| 0.91195 |
| 0.315412 |
| 0.401869 |
| 0.246862 |
| 0.561135 |
| 0.052632 |
| 0.478495 |
| 0.672694 |
| 0.51963 |
| 0.534676 |
| 0.791798 |
| 0.839716 |
| 0.637405 |
| 0.129032 |
| 0.95719 |
| 0.606426 |
| 0.407407 |
| 0.685053 |
| 0.227679 |
| 0.25 |
| 0.327645 |
| 0.814264 |
| 0.785714 |
| 0.744681 |
| 0.270992 |
| 0.644068 |
| 0.956858 |
| 0.650558 |
| 0.052632 |
| 0.679211 |
| 0.871233 |
| 0.871233 |
| 0.955407 |
| 0.270992 |
| 0.449275 |
| 0.386076 |
| 0.430303 |
| 0.672694 |
| 0.985507 |
| 0.205 |
| 0.870523 |

| |
|---|
| 0.895833 |
| 0.478495 |
| 0.556291 |
| 0.898701 |
| 0.662409 |
| 0.564211 |
| 0.51963 |
| 0.637405 |
| 0.964554 |
| 0.857341 |
| 0.797214 |
| 0.412844 |
| 0.561135 |
| 0.644068 |
| 0.729131 |
| 0.304348 |
| 0.836415 |
| 0.680357 |
| 0.911728 |
| 0.911728 |
| 0.238298 |
| 0.175 |
| 0.378981 |
| 0.378981 |
| 0.504762 |
| 0.569038 |
| 0.194444 |
| 0.498715 |
| 0.959651 |
| 0.433735 |
| 0.227679 |
| 0.6238 |
| 0.955407 |
| 0.815249 |
| 0.476712 |
| 0.962366 |
| 0.5 |
| 0.503667 |
| 0.936508 |
| 0.90781 |
| 0.90781 |
| 0.955407 |
| 0.618217 |
| 0.103774 |
| 0.08642 |
| 0.955407 |
| 0.955407 |
| 0.10101 |
| 0.475275 |
| 0.849372 |
| 0.570833 |
| 0.070423 |
| 0.180233 |
| 0.93643 |
| 0.797214 |
| 0.849083 |
| 0.981191 |
| 0.089888 |
| 0.985447 |

| |
|---|
| 0.955703 |
| 0.129032 |
| 0.129032 |
| 0.205 |
| 0.070423 |
| 0.344595 |
| 0.955407 |
| 0.430303 |
| 0.955407 |
| 0.729592 |
| 0.08642 |
| 0.803354 |
| 0.875676 |
| 0.129032 |
| 0.836415 |
| 0.932681 |
| 0.585216 |
| 0.058824 |
| 0.975713 |
| 0.134921 |
| 0.052632 |
| 0.6593 |
| 0.895833 |
| 0.471751 |
| 0.475275 |
| 0.505938 |
| 0.449275 |
| 0.849372 |
| 0.848656 |
| 0.787975 |
| 0.811377 |
| 0.449275 |
| 0.483957 |
| 0.968085 |
| 0.386792 |
| 0.177914 |
| 0.90781 |
| 0.238298 |
| 0.1875 |
| 0.564211 |
| 0.955407 |
| 0.957143 |
| 0.08642 |
| 0.05 |
| 0.117117 |
| 0.709622 |
| 0.849372 |
| 0.836415 |
| 0.932681 |
| 0.227679 |
| 0.270992 |
| 0.503667 |
| 0.955407 |
| 0.070423 |
| 0.957143 |
| 0.435435 |
| 0.573805 |
| 0.836415 |
| 0.57438 |

| |
|---|
| 0.803354 |
| 0.205 |
| 0.90781 |
| 0.386792 |
| 0.517483 |
| 0.504762 |
| 0.775806 |
| 0.449275 |
| 0.614035 |
| 0.958515 |
| 0.792126 |
| 0.151724 |
| 0.542411 |
| 0.797214 |
| 0 |
| 0.960954 |
| 0.870879 |
| 0.936663 |
| 0.936663 |
| 0.686726 |
| 0.611328 |
| 0.955407 |
| 0.982255 |
| 0.611111 |
| 0.895722 |
| 0.955407 |
| 0.205 |
| 0.743676 |
| 0.205 |
| 0.697552 |
| 0.823188 |
| 0.955407 |
| 0.870879 |
| 0.955407 |
| 0.975891 |
| 0.524887 |
| 0.274436 |
| 0.524887 |
| 0.471751 |
| 0.177914 |
| 0.511792 |
| 0.356436 |
| 0.129032 |
| 0.246862 |
| 0.301471 |
| 0.569937 |
| 0.227679 |
| 0.615534 |
| 0.15894 |
| 0.219048 |
| 0.324042 |
| 0.324042 |
| 0.298893 |
| 0.892761 |
| 0.661792 |
| 0.755302 |
| 0.139706 |
| 0.456897 |
| 0.968119 |

| |
|---|
| 0.405573 |
| 0.238298 |
| 0.955703 |
| 0.849372 |
| 0.701209 |
| 0.997976 |
| 0.169811 |
| 0.701209 |
| 0.070423 |
| 0.197861 |
| 0.197861 |
| 0.398119 |
| 0.75 |
| 0.504762 |
| 0.675045 |
| 0.205 |
| 0.991753 |
| 0.219048 |
| 0.957143 |
| 0.575258 |
| 0.524887 |
| 0.449275 |
| 0.611328 |
| 0.169811 |
| 0.76013 |
| 0.962366 |
| 0.911728 |
| 0.911728 |
| 0.811377 |
| 0.564211 |
| 0.849582 |
| 0.849372 |
| 0.197802 |
| 0.925187 |
| 0.205 |
| 0.552106 |
| 0 |
| 0.955407 |
| 0.997976 |
| 0.197861 |
| 0.997976 |
| 0.949214 |
| 0.147887 |
| 0.147887 |
| 0.836415 |
| 0.836415 |
| 0.895833 |
| 0 |
| 0.111111 |
| 0.238298 |
| 0.189944 |
| 0.744681 |
| 0.637405 |
| 0.932681 |
| 0.569038 |
| 0.08642 |
| 0.56044 |
| 0.360656 |
| 0.9201 |

| |
|---|
| 0.475275 |
| 0.801233 |
| 0.092784 |
| 0.564211 |
| 0.875676 |
| 0.433735 |
| 0.815249 |
| 0.895833 |
| 0.787975 |
| 0.6238 |
| 0.797527 |
| 0.98759 |
| 0.238298 |
| 0.998992 |
| 0.698606 |
| 0.377419 |
| 0.401869 |
| 0.15894 |
| 0.15894 |
| 0.955407 |
| 0.180233 |
| 0.180233 |
| 0.817251 |
| 0.955407 |
| 0.092784 |
| 0.221698 |
| 0.478495 |
| 0.611328 |
| 0.67446 |
| 0.08642 |
| 0.219048 |
| 0.817784 |
| 0.070423 |
| 0.611111 |
| 0.644068 |
| 0.052632 |
| 0.858921 |
| 0.686726 |
| 0.258964 |
| 0.659259 |
| 0.659926 |
| 0.968085 |
| 0.988636 |
| 0.05 |
| 0.449275 |
| 0.895833 |
| 0.524887 |
| 0.997976 |
| 0.929104 |
| 0.270992 |
| 0.227679 |
| 0.811377 |
| 0.995889 |
| 0.968085 |
| 0.932681 |
| 0.744681 |
| 0.985507 |
| 0.895833 |
| 0.895833 |

| |
|---|
| 0.895833 |
| 0.895833 |
| 0.503667 |
| 0.058824 |
| 0.997976 |
| 0.092784 |
| 0.092784 |
| 0.895833 |
| 0.231111 |
| 0.562092 |
| 0.649813 |
| 0.930521 |
| 0.960954 |
| 0.248963 |
| 0.955407 |
| 0.932681 |
| 0.405573 |
| 0.955407 |
| 0.214634 |
| 0.05 |
| 0.524887 |
| 0.531532 |
| 0.524887 |
| 0.208955 |
| 0.895833 |
| 0.058824 |
| 0.058824 |
| 0.070423 |
| 0.224299 |
| 0.911728 |
| 0.661172 |
| 0.052632 |
| 0.052632 |
| 0.503667 |
| 0.384127 |
| 0.997976 |
| 0.649813 |
| 0.814264 |
| 0.219048 |
| 0.6593 |
| 0.797214 |
| 0.710616 |
| 0.407975 |
| 0.103774 |
| 0.797214 |
| 0.151724 |
| 0.611328 |
| 0.81194 |
| 0.238298 |
| 0.956906 |
| 0.955407 |
| 0.878869 |
| 0.960954 |
| 0.955506 |
| 0.25 |
| 0.089888 |
| 0.205 |
| 0.092784 |
| 0.897269 |

| |
|---|
| 0.606426 |
| 0.680927 |
| 0.156463 |
| 0.811377 |
| 0.611328 |
| 0.615534 |
| 0.955407 |
| 0.052632 |
| 0.955407 |
| 0.997976 |
| 0.743676 |
| 0.31295 |
| 0.611111 |
| 0.129032 |
| 0.709122 |
| 0.975738 |
| 0.960997 |
| 0.803354 |
| 0.71012 |
| 0.292593 |
| 0.815249 |
| 0.490862 |
| 0.473389 |
| 0.875676 |
| 0.954382 |
| 0.564211 |
| 0.564211 |
| 0.911728 |
| 0.697552 |
| 0.08642 |
| 0.957143 |
| 0.05 |
| 0.327645 |
| 0.815249 |
| 0.697552 |
| 0.324042 |
| 0.606426 |
| 0.129032 |
| 0.803354 |
| 0.894244 |
| 0.475275 |
| 0.129032 |
| 0.129032 |
| 0.139706 |
| 0.744681 |
| 0 |
| 0.504762 |
| 0.932681 |
| 0.082192 |
| 0.253061 |
| 0.932681 |
| 0.998992 |
| 0.811377 |
| 0.611111 |
| 0.955407 |
| 0.270992 |
| 0.244726 |
| 0.272727 |
| 0.317857 |

| |
|---|
| 0.503667 |
| 0.611328 |
| 0.070423 |
| 0.512941 |
| 0.15894 |
| 0.650558 |
| 0.975941 |
| 0.975661 |
| 0.874145 |
| 0.606061 |
| 0.058824 |
| 0.6593 |
| 0.975941 |
| 0.985462 |
| 0.070423 |
| 0.686726 |
| 0.90781 |
| 0.368078 |
| 0.504762 |
| 0.836415 |
| 0.955407 |
| 0.878869 |
| 0.955407 |
| 0.955556 |
| 0.895833 |
| 0.807284 |
| 0.49354 |
| 0.169811 |
| 0.276119 |
| 0.534676 |
| 0.180233 |
| 0.052632 |
| 0.485488 |
| 0.839716 |
| 0.503667 |
| 0.344595 |
| 0.1875 |
| 0.791469 |
| 0.503667 |
| 0.875676 |
| 0.958515 |
| 0.673285 |
| 0.489529 |
| 0.089888 |
| 0.258964 |
| 0.849083 |
| 0.46 |
| 0.151724 |
| 0.304348 |
| 0.205 |
| 0.323843 |
| 0.870523 |
| 0.955407 |
| 0.51963 |
| 0.136364 |
| 0.08642 |
| 0.895833 |
| 0.69808 |
| 0.25 |

| |
|---|
| 0.811377 |
| 0.975764 |
| 0.701209 |
| 0.590631 |
| 0.473389 |
| 0.189944 |
| 0.785714 |
| 0.089888 |
| 0.972399 |
| 0.955407 |
| 0.849372 |
| 0.985507 |
| 0.955407 |
| 0.815249 |
| 0.136364 |
| 0.697552 |
| 0.058824 |
| 0.111111 |
| 0.811377 |
| 0.058824 |
| 0.355482 |
| 0.05 |
| 0.975866 |
| 0.975866 |
| 0.99899 |
| 0.058824 |
| 0.590631 |
| 0.644068 |
| 0.997976 |
| 0.895833 |
| 0.561135 |
| 0.139706 |
| 0.205 |
| 0.092784 |
| 0.504762 |
| 0.052632 |
| 0.197802 |
| 0.998989 |
| 0.911839 |
| 0.169811 |
| 0.895833 |
| 0.070423 |
| 0.955407 |
| 0.205 |
| 0.156463 |
| 0.875676 |
| 0.975687 |
| 0.258964 |
| 0.503667 |
| 0.449275 |
